# Reproducibility of protein X-ray diffuse scattering and potential utility for modeling atomic displacement parameters

**DOI:** 10.1101/2021.01.24.428002

**Authors:** Zhen Su, Medhanjali Dasgupta, Frédéric Poitevin, Irimpan I. Mathews, Henry van den Bedem, Michael E. Wall, Chun Hong Yoon, Mark A. Wilson

## Abstract

Protein structure and dynamics can be probed using X-ray crystallography. Whereas the Bragg peaks are only sensitive to the average unit-cell electron density, the signal between the Bragg peaks -- diffuse scattering -- is sensitive to spatial correlations in electron-density variations. Although diffuse scattering contains valuable information about protein dynamics, the diffuse signal is more difficult to isolate from the background compared to the Bragg signal, and the reproducibility of diffuse signal is not yet well understood. We present a systematic study of the reproducibility of diffuse scattering from isocyanide hydratase (ICH) in three different protein forms. Both replicate diffuse datasets and datasets obtained from different mutants were similar in pairwise comparisons (Pearson correlation coefficient (CC) ≥0.8). The data were processed in a manner inspired by previously published methods using custom software with modular design, enabling us to perform an analysis of various data processing choices to determine how to obtain the highest quality data as assessed using unbiased measures of symmetry and reproducibility. The diffuse data then were used to characterize atomic mobility using a liquid-like motions (LLM) model. This characterization was able to discriminate between distinct anisotropic atomic displacement parameter (ADP) models arising from different anisotropic scaling choices that agreed comparably with the Bragg data. Our results emphasize the importance of data reproducibility as a model-free measure of diffuse data quality, illustrate the ability of LLM analysis of diffuse scattering to select among alternative ADP models, and offer insights into the design of successful diffuse scattering experiments.

## I. INTRODUCTION

In X-ray crystallography, the sharp Bragg reflections are the main source of information for structure determination; however, they only contain information about the average electron density of the unit cell. Diffuse scattering, on the other hand, contains information about the spatial correlations of electron density variations, and thus can, in principle, distinguish among different atomic motions that yield the same mean electron density.^1–3^ In addition, recent studies suggest that diffuse scattering might be used to extend the resolution of density maps beyond the resolution limit of the Bragg peaks,^4,5^ motivating further rigorous investigation of this possibility.^6^

Early studies of protein diffuse scattering focused on interpreting features in individual diffraction images.^7–15^ Since the development of modern diffuse data processing methods,^16,17^ protein diffuse scattering studies have mostly focused on working with three-dimensional (3D) datasets. In addition to improvements in light sources and detectors, notable developments in 3D data processing include finer sampling in reciprocal space to model long-range correlations,^18^ rescuing useful diffuse data from experiments designed for Bragg diffraction,^19^ extracting finely sampled 3D datasets from serial femtosecond X-ray crystallography (SFX) experiments with X-ray free-electron lasers (XFEL),^4^ increasing data quality via improved rejection of the solvent contribution and multivariate analysis methods,^20^ and a major advance in the scaling and merging of data from multiple crystals,^21^ yielding a substantial improvement in data quality.

Given the variety of approaches to data processing, and the emerging importance of diffuse scattering for modeling protein dynamics, we sought to gain more insight into some fundamental questions about protein diffuse scattering data: How reproducible are single-crystal diffuse datasets? What is the influence of point mutations on the diffuse signal? How do changes in the data translate into differences in a model? What are the consequences of different data processing choices for data quality? Can diffuse scattering data discriminate between different models of atomic mobility that agree equally well with the Bragg data?

Here we address each of these questions in a study of diffuse scattering from crystalline ICH. We selected the ICH system because it diffracts X-rays to atomic resolution at ambient temperature, has clearly visible diffuse features in ambient temperature X-ray diffraction datasets, and displays large concerted motion of an *α*-helix that is modulated by the chemical state of the active site nucleophile.^22^ Upon formation of the catalytic thioimidate intermediate, this helix becomes more mobile and permits water to enter the active site and complete the reaction. Because the extent of this concerted, functionally important *α*-helix motion can be controlled using various experimental tools, ICH is a very promising system for exploring the utility of diffuse scattering data for characterizing functional protein dynamics.

Specifically, we address the above questions using multiple datasets collected from wild-type (WT) ICH and two mutants (G150A, G150T) that affect helix motion. Using a modular data processing pipeline in Python that we developed, we assessed quantitatively the reproducibility of the data and the influence of various data processing choices on the final quality of the datasets. Because our processing pipeline is modular in construction, individual steps can be easily modified and their impact on data quality separately evaluated. In this workflow, we assessed the data quality using unbiased measures of the internal consistency (CC_1/2_)^23^ and reproducibility (CC_Rep_), which we compared with prior metrics such as CC_Laue_ and CC_Friedel_. Finally, we analyzed the diffuse data using simple phenomenological models of correlated protein motion: the LLM model^9,11^ using three different treatments of ADPs (B factors); and an independent rigid-body translational motions (RBT) model.^1,4^ This analysis yields insights into the impact of the various data processing choices on the model parameters and the agreement with the data.

Overall, the results of this study indicate that single-crystal diffuse datasets can be measured reproducibly from WT and mutant ICH crystals (CC_Rep_ ≥ 0.81 to 1.4Å resolution). Differences in diffuse scattering among different ICH mutants are small when assessed directly using the data, yet are still detectable using the LLM analysis. Importantly, the LLM analysis showed that diffuse scattering can discriminate between ADP models that fit the Bragg data equally well. In addition, the LLM models of ICH yield higher correlations with the data than the independent RBT models. Finally, a systematic investigation of the influence of data processing methods using our Python workflow yielded a matrix of data quality measures, revealing insights into best practices for data collection and processing. In particular, the results emphasize the importance of background subtraction for increasing data quality, and highlight the benefits of adding a step to remove some of the variation in the isotropic radial intensity profiles.^20^

## II. METHODS

### ICH protein expression and crystallization

WT, G150A, and G150T *Pseudomonas protegens* Pf-5 (formerly *Pseudomonas fluorescens*) ICH proteins were expressed in BL21(DE3) *E. coli*, purified by Ni^2+^-metal affinity chromatography, and crystallized by hanging drop vapor equilibration as previously described.^22,24^ Briefly, ICH crystals were grown at room temperature (~22°C) by mixing 2 μl of protein at 20 mg/ml with 2 μl of reservoir (22-24% PEG 3350, 100 mM Tris-HCl, pH 8.6, 200 mM magnesium chloride and 2 mM dithiothreitol (DTT)) and typically took one week to reach maximum size. Microseeding of drops equilibrated for 6-12 hours improved crystal size and morphology. As previously noted,^22^ G150T crystals form in a different space group (C2/I2) than WT and G150A crystals (P2_1_) even when seeded with WT crystals. The largest crystals were ~700×700×150 μm, although typically G150A and G150T ICH crystals grew with a more compact prismatic habit than WT ICH.

### Diffuse and Bragg X-ray data collection

To study the reproducibility of diffuse scattering in independent samples, data were collected from three crystals of each form of ICH. For simplicity, these datasets are denoted as WT-1, WT-2, WT-3, G150A-1, G150A-2, G150A-3, G150T-1, G150T-2, and G150T-3, indicating the WT, G150A, and G150T mutant ICH proteins. Crystals were mounted in 10 μm thick glass number 50 borosilicate capillaries (Hampton Research) ranging from 0.7 to 1.0 mm diameter and sealed with wax. Excess solution near the crystal was wicked away while retaining a small volume of reservoir solution in the end of the capillary to maintain vapor equilibrium. For WT ICH, the plate-like crystals were mounted “edge-on”, such that their shortest axis was roughly parallel to the capillary axis. In this geometry, the X-ray beam illuminates approximately equivalent volumes of the crystal during rotation about the spindle axis, which was parallel with the capillary axis. G150A and G150T ICH crystals had more prismatic habits than WT ICH and did not require special orientation for data collection.

Diffraction data were collected at 274 K on BL12-2 at the Stanford Synchrotron Radiation Lightsource (SSRL) using 16 keV incident X-rays and shutterless data collection with 0.5° rotation/image, 0.3 sec/exposure, and 98% attenuation. The data were recorded on a PILATUS 6M pixel array detector (PAD) with roughly 0.95Å resolution at the edge of the detector for each dataset. Absorbed doses were approximately 2-4×10^4^ Gy per crystal as calculated using https://bl831.als.lbl.gov/xtallife.html.^25^ Doses were kept low to minimize X-ray-induced oxidation of the catalytic Cys101 nucleophile to sulfenic acid, which has been previously reported.^22,24,26^ To allow subtraction of the capillary background scattering from the diffraction images, non-crystal background diffraction patterns were collected using identical parameters to those used for crystal data collection but by increasing the exposure time and slightly shifting the X-ray beam to the region of the capillary away from the crystal, as shown in Fig. 1. The exposure time was 1 second per image for the non-crystal background patterns in order to accumulate more scattered photons and reduce error in the background measurements. The background images were later scaled by the ratio of the exposure times to be equal to the data images.

**FIG. 1.**
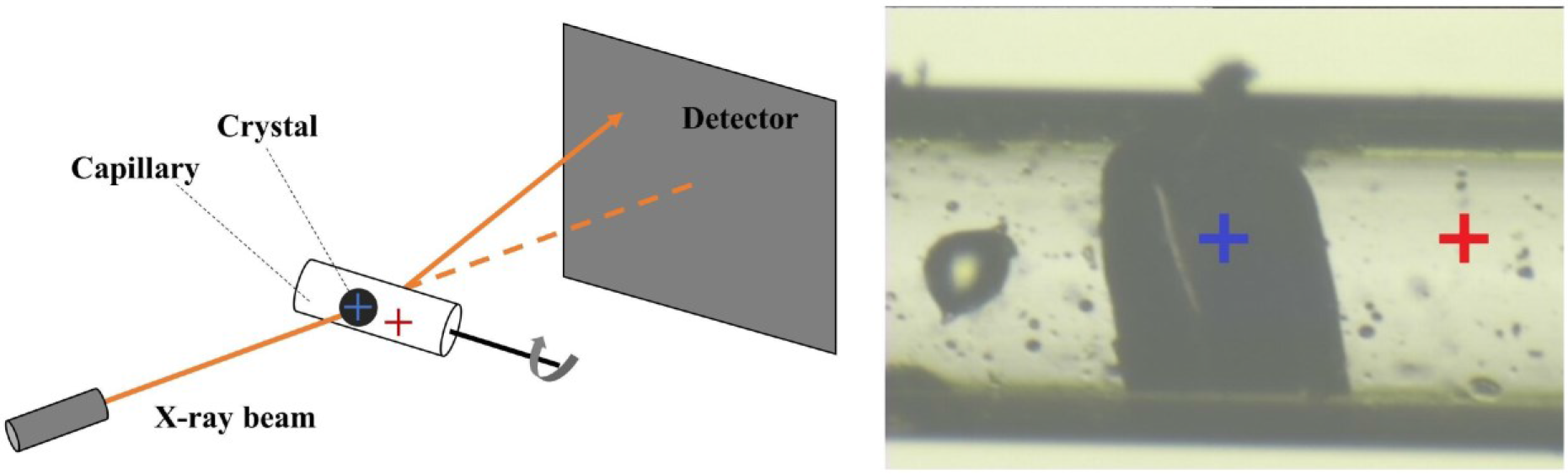
Illustration of distinction between crystal exposure and background exposure. (Left) experimental setup for diffuse data collection. (Right) the dark object in the center is the WT-1 crystal and the blue cross marks the X-ray beam position for crystal diffraction measurements. The crystal is hydrated by a buffer solution inside the capillary. The non-crystal background images were collected by translating the capillary so that the X-ray beam (red cross) only interacts with the capillary, buffer, and air bubbles. Crystal and background diffraction pattern pairs were collected in each orientation.

### Analysis of Bragg data

The Bragg data from each crystal were indexed and scaled using XDS,^27^ Pointless,^28^ and Aimless^29^ with statistics reported in Table S1. For G150T, the data were (equivalently) reindexed from C2 to I2, yielding unit cells more comparable to those of WT and G150A ICH datasets in space group P2_1_. Structures of WT, G150A, and G150T ICH were refined against these data in PHENIX (v1.17.1-3660)^30^ using riding hydrogen atoms and restrained anisotropic ADPs with weight optimization for coordinate and ADP refinements. Riding hydrogen atoms have their positions calculated from the geometry of the bonded heavier atoms upon which they “ride” and thus contribute to both the calculation of model structure factors and non-bonded contacts without adding additional refinement parameters. As noted previously,^22,24^ Ile152 is a Ramchandran outlier in all structures except G150T and is well-supported by the electron density maps in all cases. We also refined protein structures using the Refmac5 package (v5.8.0266)^31^ in the CCP4 suite of programs^32^ in order to compare the behavior of Refmac5- and PHENIX-refined models against the same datasets. The Refmac5 refinements used riding hydrogen atoms and restrained anisotropic ADPs with a matrix weight term of 0.2-0.4. This range for the matrix weight term produced bond length root mean square differences (RMSD) in Refmac5-refined models that were comparable to those of the PHENIX-refined models. These refinement protocols produced models with similar R_free_/R_work_ for the Bragg data (see Tables S2 and S3 for refined model statistics and PDB codes). Despite similar R_free_/R_work_ values, the anisotropic ADPs of the PHENIX-refined models have anisotropy ratios (the ratio of smallest to largest eigenvalues to the ADP variance-covariance matrix) that were lower (more anisotropic) than the Refmac5-refined models (Fig. S1), while the ADP magnitudes in both models are highly similar (Fig. S2). This difference in anisotropies was observed for all models, but was most pronounced in the WT datasets. Moving from isotropic to anisotropic ADPs decreased the R_free_ value by ~3-4% in all datasets in both Refmac5 (Table S2) and PHENIX (Table S3), confirming that anisotropic ADPs yield higher agreement with the Bragg diffraction data than isotropic displacements, and justifying the use of the additional parameters.

The differences in the anisotropic ADPs of models refined in Refmac5 and PHENIX against the same dataset were surprising initially; however, we were able to demonstrate that they are well explained by differences in the overall anisotropic scaling matrices. To demonstrate this, we obtained refined anisotropic scaling parameters from the headers of both the Refmac5 and PHENIX models after zero cycles of refinement against the same data in PDB-REDO.^33^ Using PDB-REDO in this way invokes the Refmac5 refinement engine to recover the anisotropic scale parameters and guarantees that all models are handled in an identical fashion. The resulting anisotropic scaling matrices for Refmac5 and PHENIX models are often different (see Table S4).

To determine whether differing anisotropic scale matrices are responsible for the different anisotropic ADP models obtained using Refmac5 and PHENIX refinement, we calculated difference anisotropic scaling matrices and used them to rescale the model ADPs (Supplementary Material section III). These difference matrices were added to the ANISOU records for each atom in the model after being made traceless by subtracting trace/3 from each diagonal element to ensure that B_eq_ would not be altered (Table S4). Using the difference matrices, we found that we were able to convert a PHENIX-refined anisotropic ADP model into one that resembles its Refmac5-refined counterpart, and vice versa (Fig. S3, S4, S5; Supplementary Material section III). Importantly, this rescaling of the models scarcely influenced the agreement with the Bragg data but could substantially influence the agreement of LLM models with the diffuse data (see below).

### Construction of 3D diffuse scattering maps

Our 3D diffuse map construction pipeline includes six image pre-processing steps followed by 3D merging and two volume processing steps (Fig. 2). The pre-processing steps were designed to convert the raw intensities into useful diffuse signals and to reject non-diffuse intensities such as Bragg peaks, bad pixels, random noise, and isotropic and anisotropic background. In order of application, these steps were: (1) detector masking; (2) bad pixel removal; (3) non-crystal background pattern subtraction; (4) pixel position and intensity corrections; (5) Bragg peak cleaning; and (6) image scaling and radial profile variance removal.^20^ Starting with raw diffraction patterns, step (1) was to mask out obvious bad pixels in the detector, including dead pixels, shadows, and grid lines between detector panels, pixels near the beamstop or with intensities that were either non-positive or greater than 10,000 photons. Step (2) was to perform a deeper cleaning of bad pixels, by masking pixels with intensities that are beyond 5 standard deviations from the mean value inside a 11×11 square window. Steps (1) and (2) were also applied to non-crystal background patterns in the same manner. In step (3), the filtered background patterns were scaled by the exposure time and subtracted frame-by-frame from the matching crystal diffraction patterns (see Methods). In step (4), pixel positions were corrected by the parallax broadening effect in the PILATUS 6M detector,^20^ and raw pixel intensities were converted to scattering intensities by applying polarization,^16^ solid-angle,^16^ and detector absorption corrections.^21^

**FIG. 2.**
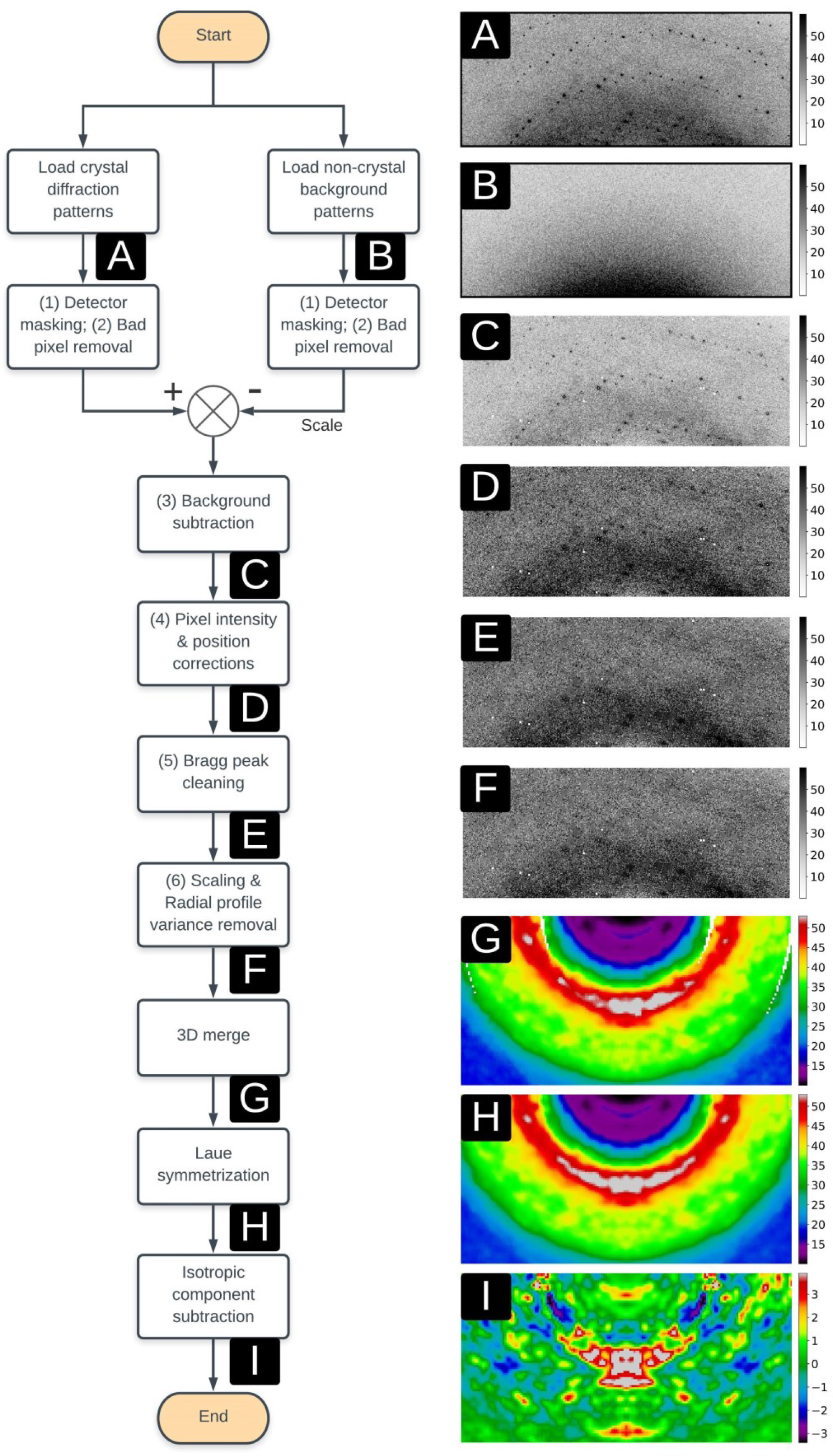
Data analysis pipeline from raw diffraction patterns to a Laue-symmetrized anisotropic diffuse map. Numbers (1)-(6) correspond to the same image pre-processing substeps as mentioned in Methods. Following this pipeline, the (A) crystal diffraction and (B) non-crystal background patterns are applied with the user-defined detector mask and a deeper bad pixel removal step based on pixel positions and intensities. The non-crystal background patterns are then scaled with the exposure time and subtracted from crystal diffraction patterns, giving rise to the (C) background subtracted patterns, followed by multiple pixel intensity and position corrections to produce the (D) corrected diffraction patterns. Bragg peaks are predicted in positions and then replaced with median intensities to generate (E) patterns without Bragg peaks, followed by image scaling and the radial profile variance removal method which end up with the final pre-processed diffraction patterns (F). These patterns are merged into a (G) 3D diffraction volume using indexing results and orientations from the goniometer. This 3D volume is then applied with Laue symmetrization to generate the (H) Laue-symmetrized diffraction volume, followed by the isotropic component subtraction step which produces the final (I) Laue-symmetrized anisotropic diffuse map. For improved visualization, panels (G)-(I) were created using more finely sampled diffraction volumes than were used in data quality evaluation and modelling.

In step (5), Bragg peaks were predicted in positions and further cleaned, although some peaks were already removed in step (2) due to their strong intensities. Pixels were mapped into reciprocal space and converted into fractional Miller indices (*h,k,l*) using the XDS^27^ indexing result. Intensities were identified as belonging to Bragg peaks if their indices (*h,k,l*) are all within 0.25 to the nearest integers. The intensity of each Bragg pixel was replaced with the median value in a 11×11 square window centered on this pixel. The order of filtering, background subtraction and correction steps described above is flexible, but Bragg peaks must be cleaned before image scaling and radial profile variance removal in step (6). The diffraction pattern after the previous five steps is considered as a combination of diffuse scattering, random noise, and isotropic signals from multiple sources such as the crystal, water, and air diffraction. Random noise can be averaged out later in the 3D merging stage, so dealing with the isotropic signal was the main focus in step (6). Firstly, the diffraction pattern was scaled using the radial intensity profile scale factor, calculated by minimizing the L2 distance between radial intensity profiles of the target diffraction pattern and a fixed reference diffraction pattern (the first pattern of each dataset in our method). Another radial profile variance removal step, first described in Peck *et al.*,^20^ was applied by performing principal component analysis (PCA) on the matrix of the scaled radial profiles and subtracting the contribution from the subspace of the three largest eigenvalues, as shown in Fig. S6.

Each diffraction pattern corresponds to the intersection of an Ewald sphere surface with the 3D diffraction volume. Diffraction patterns after six pre-processing steps were mapped into reciprocal space using crystal orientations and experiment parameters, including the X-ray wavelength, detector distance (*z_d_*), and pixel size. The orientation information was calculated from XDS^27^ indexing results (including the **A** matrix) as well as the relative rotation angles in the experiment. Each pixel located at (*x,y,z_d_*) on the detector corresponds to fractional Miller indices (*h,k,l*) in reciprocal space, which lies within a voxel in the 3D diffraction volume. The voxel value was measured as the average intensity of all pixels that were assigned to it. To avoid contamination arising from Bragg peaks, we rejected every pixel located within a 0.5×0.5×0.5 box centered on the nearest reciprocal-space point with integer Miller indices. This Bragg rejection step can be equivalently applied in the image pre-processing stage by masking pixels rather than replacing them with median intensities. In previous work, three different methods were mentioned regarding removal of Bragg pixels, by either filtering out Bragg pixels,^16–18^ replacing intensities,^34^ or preserving Bragg peak intensities together with diffuse scattering features.^21^ In this work, we chose to filter out all pixels in Bragg peak positions, as we were interested in large-scale diffuse features that vary on a length scale longer than the separation between Bragg peaks. The other two methods are useful for obtaining more finely sampled datasets and analyzing sharper diffuse scattering features.

The 3D diffraction volume obtained by merging all crystal diffraction patterns (denoted as the raw unsymmetrized map), was symmetrized according to its Laue/Friedel point group into a Laue-/Friedel-symmetrized map. For the ICH crystal, Friedel symmetrization averages 2 voxels related by an inversion symmetry, and Laue symmetrization averages 4 voxels related by the Laue group (2/m for all nine crystals). To remove the scattering from other sources such as water, air, and uncorrelated protein motions, the symmetrized map was further processed with an isotropic component subtraction step by subtracting the radially averaged 3D volume to get the symmetrized anisotropic diffuse scattering map (Fig. 2). The 3D anisotropic diffuse scattering map is called the diffuse map in this work, and is assumed to contain anisotropic diffuse scattering features arising from correlated motions in the crystal, although further analysis and modeling are still required to confirm this. The *dspack* package for the whole analysis pipeline, including image pre-processing steps, 3D merging, and volume operations, is available online: https://github.com/zhenwork/dspack.

### Evaluation of the quality of diffuse scattering maps

The diffuse map produced by our analysis pipeline contains both anisotropic diffuse scattering from correlated protein motions and any merging artifacts that have anisotropic features. Previous studies^16,18,20^ have used symmetry metrics such as CC_Laue_ and CC_Friedel_ (see Table 1) to assess the quality of 3D diffuse datasets, calculated using the function,

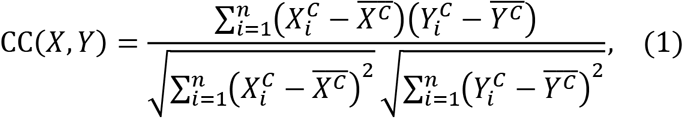

where *X^C^* and *Y^C^* represent two vectors sampled from *n* common voxels of unsymmetrized (*X*) and Laue-/Friedel-symmetrized anisotropic maps (*Y*), respectively, and 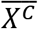 and 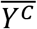 represent the mean values. The symmetrized maps were calculated by averaging related Laue/ Friedel voxels, as described in the previous section.

**TABLE 1.** The diffuse data quality statistics of each dataset. CC_Cross_ was not evaluated for G150T datasets due to the different space group.

| Sample | WT-1 | WT-2 | WT-3 | G150A-1 | G150A-2 | G150A-3 | G150T-1 | G150T-2 | G150T-3 |
| --- | --- | --- | --- | --- | --- | --- | --- | --- | --- |
| Compl<sup>1</sup> | 98.36 | 100.0 | 99.30 | 100.0 | 98.80 | 100.0 | 99.84 | 100.0 | 98.97 |
| $CC_{Friedel}$ | 0.93 | 0.91 | 0.91 | 0.93 | 0.91 | 0.91 | 0.91 | 0.92 | 0.94 |
| $CC_{Laue}$ | 0.90 | 0.87 | 0.87 | 0.88 | 0.86 | 0.85 | 0.86 | 0.88 | 0.91 |
| $CC_{1/2}$ | 0.85 | 0.78 | 0.81 | 0.81 | 0.76 | 0.77 | 0.82 | 0.84 | 0.89 |
| $CC_{Rep}$ | 0.86 | 0.84 | 0.85 | 0.82 | 0.81 | 0.82 | 0.88 | 0.87 | 0.89 |
| $CC_{Cross}$ | 0.83 | 0.84 | 0.85 | 0.84 | 0.84 | 0.85 | --- | --- | --- |

<sup>1</sup>Compl represents the completeness (%) of the diffuse data.

Here, we use two additional metrics for quality evaluation of diffuse maps: the data symmetry (CC_1/2_) and reproducibility (CC_Rep_). CC_1/2_ is an accepted metric for assessing the quality of Bragg diffraction data,^23^ and also has been used for diffuse scattering data.^21^ The CC_1/2_ metric was calculated using *phenix.merging_statistics*^30^ with the unsymmetrized anisotropic map as input. The CC_1/2_ measures whether the diffuse map follows the target symmetry, but it can be misleading if the diffuse map contains substantial anisotropic background features that partly obey the symmetry. To address this problem, we introduce another metric, CC_Rep_, which is the average CC between diffuse maps of the selected dataset and other independent datasets measured from different crystals of the same protein (see Table S5). For example, in this work the CC_Rep_ for the first dataset of WT ICH (WT-1) is measured as the average CC of CC(WT-1, WT-2) and CC(WT-1, WT-3), where WT-2 and WT-3 are two additional datasets measured from crystals of WT ICH.

### Modeling diffuse scattering data with the LLM model

We applied the LLM model using the refine_llm.py script in the Lunus software package,^17^ starting with inputs of the experimental Laue-symmetrized diffuse map and the corresponding PDB file refined from Bragg data of the same dataset (Table S2). The LLM model uses the following equation to describe the diffuse intensity *I_d_*(***q***):

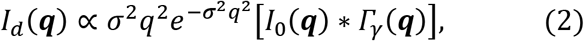

where *I_0_*(***q***) is the squared structure factor of the unperturbed crystal, and *Γ_γ_*(***q***) is the Fourier transform of the function describing the distance-dependence of the atomic displacement correlations. The LLM model has two refinable parameters: the average atomic displacement *σ*, which estimates the average amplitude of atomic motions, and the correlation length *γ*, which is the characteristic length scale of correlated atomic displacements.^9,11^ Before comparing *I_d_*(***q***) to the experimental data, *I_d_*(***q***) was Laue-symmetrized and the isotropic component was removed, to ensure that both maps were processed in a similar way. The parameters *σ* and *γ* of the LLM model were optimized using the Powell minimization method in scipy.optimize,^35^ using the CC between the model and the data as a target -- the highest value of the correlation is denoted as CC_LLM_.

In Eq. (2), *I_0_*(***q***) is computed after setting the individual B factors to zero. In addition to this model, here we consider models in which the individual B factors are preserved. Preserving the B factors yields the following equation for the LLM (Supplementary Material section IV):

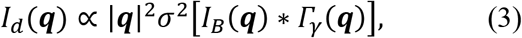

where *I_B_*(***q***) is the Bragg intensity computed using the individual ADPs in the PDB file, and *σ* is the amplitude of the correlated atomic displacements (assumed to be the same for all atoms). Eq. (3) is the same as Eq. (2), with *I_0_*(***q***) replaced by *I_B_*(***q***), and with the overall Debye-Waller factor 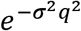 replaced by unity. Note that, whereas in Eq. (2), sufficiently high values of *σ* influence the resolution-dependence of the diffuse intensity, in Eq. (3), *σ* only influences the overall scale of the intensity. Because in our study the diffuse data are not placed on an absolute scale, and the CC target we use for optimization is not sensitive to the absolute scale, we cannot determine the value of *σ* using Eq. (3).

We used fits to Eq. (2) to assess whether the diffuse intensity is more accurately described using LLM models with individual ADPs. Eq. (2) was used directly for the case of zero ADPs, and *I_0_*(***q***) was replaced by *I_B_*(***q***) for the case of isotropic and anisotropic ADPs. In calculating *I_0_*(***q***) and *I_B_*(***q***), multiple conformations were handled by selecting only the A conformations and setting the occupancies to unity. In the case of zero ADPs, we interpret the value of *σ* after fitting the model as being indicative of the amplitude of motion of the atoms; however, in the case of individual ADPs, *σ* is smaller, for reasons described above, and the precise value is not as meaningful; in this case, we only consider whether the value of *σ* refines to nearly zero, making the overall Debye-Waller factor close to unity. In this limit, Eq. (2) reduces to Eq. (3), indicating that the model is consistent with the use of this equation. If *σ* does not refine to something close to zero (as is the case for some models we consider here), it indicates a possible inconsistency with Eq. (3).

The isotropic ADPs were calculated as B_eq_ values from the anisotropic ADPs in the input PDB file that were previously refined against the Bragg data. Anisotropic ADPs contain information about both the direction and the amplitude of atomic motion, while the isotropic ADPs contain only information about displacement amplitude. To further examine the utility of using the LLM model for diffuse data analysis, we also fit the diffuse data using a RBT model for comparison, as was performed in a previous study.^36^ The RBT model assumes that the only correlated motions are rigid-body translations of asymmetric units and does not include rigid-body rotations and/or correlations between rigid units. The RBT contains a single fitting parameter *σ* that describes the average translational displacement of the asymmetric unit. Lunus software^17^ was used to refine *σ* with respect to the CC of the model with the data. The best-fit correlation of the RBT model to the experimental data, denoted CC_RBT_, was compared with CC_LLM_ to determine which physical model was in better agreement with the processed diffuse maps.

### Determining the importance of various steps in the analysis pipeline

There are several reported methods^4,16,17,20,21,34^ for producing 3D protein diffuse scattering datasets, and they differ with respect to image pre-processing, scaling, and radial profile normalization techniques. In this work, we only focused on the most commonly used methods for processing single crystal synchrotron diffuse data^16,17,20,34^ as described in Methods, and then studied the effects of non-crystal background subtraction, pixel position and intensity corrections,^16,20,21^ radial profile variance removal, and per-image scale factors on the quality and reproducibility of the extracted diffuse scattering maps. We evaluated the impact of each of these processing steps on data quality by sequentially omitting each step in the standard pipeline as well as testing the influence of different scale factors on final data quality. Different processing choices were evaluated using multiple diffuse scattering quality metrics, including CC_1/2_ and CC_Rep_. A similar type of analysis was used by Meisburger *et al.*^21^ to assess different approaches to merged diffuse data using a CC_1/2_ statistic.

For the data processing choice analysis, we capitalized on the modular design of our developed program to turn on, turn off, or tune parameters in specific processing steps. For the present study, choices were assessed by eliminating individual data processing steps and determining the effect on the CC_Friedel_, CC_Laue_, CC_1/2_, CC_Rep_, and CC_LLM_ values. In total, we studied seven data processing choices, including (A) the standard pipeline, as well as processing that omits either (B) the non-crystal background image subtraction, (C) the polarization correction,^16^ (D) the radial profile variance removal,^20^ (E) the solid-angle correction,^16^ (F) the detector absorption correction,^21^ or (G) the parallax correction.^20^ The values of CC_Friedel_, CC_Laue_, CC_1/2_, CC_Rep_, and CC_LLM_ resulting from these processing choices is summarized in Table 2 and Table S6.

**TABLE 2.**
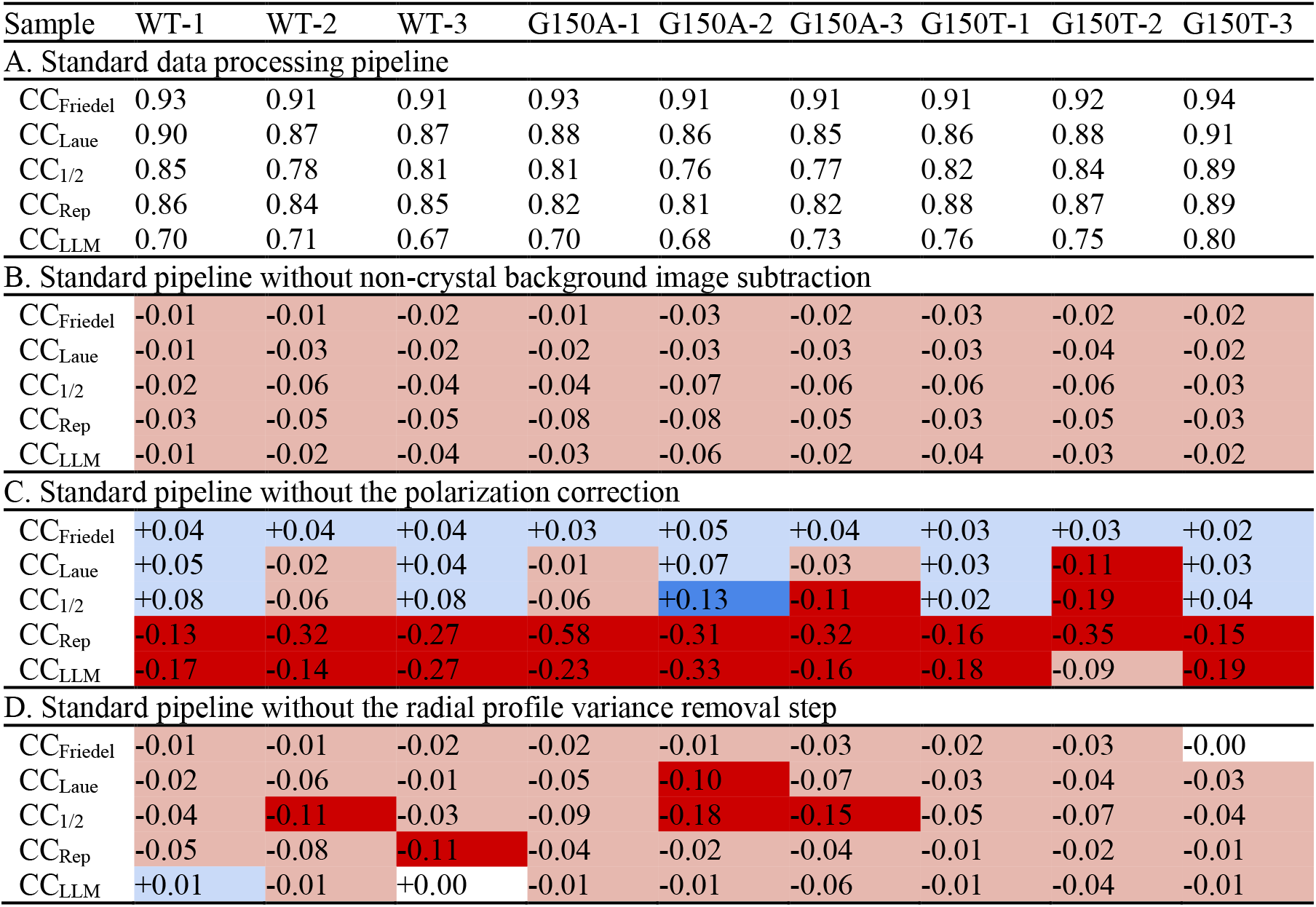
The CC statistics of each dataset are analyzed with different data processing choices. The diffuse map generated by each processing method was evaluated with five CC metrics: CC_Friedel_, CC_Laue_, CC_1/2_, CC_Rep_, and CC_LLM_ (anisotropic ADP model). Method A (standard processing pipeline) contains real CC values of each dataset up to 1.4Å, while other methods (B)-(D) are filled with relative CC changes compared to those in method A. Cells in (B)-(D) are colored with four different colors depending on the relative changes. A cell is colored as white if the relative CC change is ±0.00, as light blue/red if CC increases/decreases by less than 0.1, otherwise it will be colored as dark blue/red.

| Sample | WT-1 | WT-2 | WT-3 | G150A-1 | G150A-2 | G150A-3 | G150T-1 | G150T-2 | G150T-3 |
| --- | --- | --- | --- | --- | --- | --- | --- | --- | --- |
| **A. Standard data processing pipeline** |  |  |  |  |  |  |  |  |  |
| $CC_{Friedel}$ | 0.93 | 0.91 | 0.91 | 0.93 | 0.91 | 0.91 | 0.91 | 0.92 | 0.94 |
| $CC_{Laue}$ | 0.90 | 0.87 | 0.87 | 0.88 | 0.86 | 0.85 | 0.86 | 0.88 | 0.91 |
| $CC_{1/2}$ | 0.85 | 0.78 | 0.81 | 0.81 | 0.76 | 0.77 | 0.82 | 0.84 | 0.89 |
| $CC_{Rep}$ | 0.86 | 0.84 | 0.85 | 0.82 | 0.81 | 0.82 | 0.88 | 0.87 | 0.89 |
| $CC_{LLM}$ | 0.70 | 0.71 | 0.67 | 0.70 | 0.68 | 0.73 | 0.76 | 0.75 | 0.80 |
| **B. Standard pipeline without non-crystal background image subtraction** |  |  |  |  |  |  |  |  |  |
| $CC_{Friedel}$ | -0.01 | -0.01 | -0.02 | -0.01 | -0.03 | -0.02 | -0.03 | -0.02 | -0.02 |
| $CC_{Laue}$ | -0.01 | -0.03 | -0.02 | -0.02 | -0.03 | -0.03 | -0.03 | -0.04 | -0.02 |
| $CC_{1/2}$ | -0.02 | -0.06 | -0.04 | -0.04 | -0.07 | -0.06 | -0.06 | -0.06 | -0.03 |
| $CC_{Rep}$ | -0.03 | -0.05 | -0.05 | -0.08 | -0.08 | -0.05 | -0.03 | -0.05 | -0.03 |
| $CC_{LLM}$ | -0.01 | -0.02 | -0.04 | -0.03 | -0.06 | -0.02 | -0.04 | -0.03 | -0.02 |
| **C. Standard pipeline without the polarization correction** |  |  |  |  |  |  |  |  |  |
| $CC_{Friedel}$ | +0.04 | +0.04 | +0.04 | +0.03 | +0.05 | +0.04 | +0.03 | +0.03 | +0.02 |
| $CC_{Laue}$ | +0.05 | -0.02 | +0.04 | -0.01 | +0.07 | -0.03 | +0.03 | -0.11 | +0.03 |
| $CC_{1/2}$ | +0.08 | -0.06 | +0.08 | -0.06 | +0.13 | -0.11 | +0.02 | -0.19 | +0.04 |
| $CC_{Rep}$ | -0.13 | -0.32 | -0.27 | -0.58 | -0.31 | -0.32 | -0.16 | -0.35 | -0.15 |
| $CC_{LLM}$ | -0.17 | -0.14 | -0.27 | -0.23 | -0.33 | -0.16 | -0.18 | -0.09 | -0.19 |
| **D. Standard pipeline without the radial profile variance removal step** |  |  |  |  |  |  |  |  |  |
| $CC_{Friedel}$ | -0.01 | -0.01 | -0.02 | -0.02 | -0.01 | -0.03 | -0.02 | -0.03 | -0.00 |
| $CC_{Laue}$ | -0.02 | -0.06 | -0.01 | -0.05 | -0.10 | -0.07 | -0.03 | -0.04 | -0.03 |
| $CC_{1/2}$ | -0.04 | -0.11 | -0.03 | -0.09 | -0.18 | -0.15 | -0.05 | -0.07 | -0.04 |
| $CC_{Rep}$ | -0.05 | -0.08 | -0.11 | -0.04 | -0.02 | -0.04 | -0.01 | -0.02 | -0.01 |
| $CC_{LLM}$ | +0.01 | -0.01 | +0.00 | -0.01 | -0.01 | -0.06 | -0.01 | -0.04 | -0.01 |

To study the effect of the choice of merging approach on data quality, we computed diffraction image scale factors using four different signal sources: (A) the profile of the image intensity vs. the scattering vector length, (B) the average intensity in the isotropic ring, (C) the average intensity in the diffraction image, and (D) the Bragg peaks. For (A), the profile in each image was scaled to minimize the difference with respect to the profile in a reference image, using intensities within the resolution range (up to 1.4Å). For (B), the scale factor was computed as the ratio of the average pixel intensities within the water ring region (5Å-1.82Å). For (C), the scale factor was computed as the ratio of average pixel intensities within the resolution range. For (D), the Bragg intensity scale factors reported by *dials.scale*^37,38^ were used. They are denoted as the (A) radial profile, (B) water ring, (C) overall, and (D) Bragg scale factor, respectively. The standard pipeline in this work uses method (A). The effectiveness of a particular scale factor was evaluated with data quality metrics of the diffuse map processed using that scale factor. The data quality statistics of each type of scale factor are summarized in Table S7. This table also includes another four choices (E)-(H) where the radial profile variance removal step was turned off as the scale factor was switched from (A) to (D) successively.

## III. RESULTS

### WT and mutant ICH structures and helix motion

Prior work with ICH showed that X-ray photooxidation of Cys101 results in concerted motion of a helix near the active site that is also observed during formation of the catalytic thioimidate intermediate.^22^ These cysteine modification-activated motions in ICH^26^ occur owing to transient loss of negative charge on the catalytic cysteine thiolate and facilitate later steps in catalysis. Engineered mutations at residue 150 (e.g., G150A, G150T) also favor shifted conformations of the helix to varying degrees. Because the concerted motion of this helix can be modulated by mutation and the charge of the Cys101 S*γ* atom, ICH is an attractive system for exploring diffuse scattering as a probe of functional correlated protein motions.

In this work, structural models refined against replicate Bragg datasets that were collected simultaneously with the diffuse scattering data (see below) are essentially identical (0.02-0.03 Å C_*α*_ RMSD). The refined WT and G150A ICH models are also highly similar (~0.05-0.07 Å C_*α*_ RMSD). As observed before,^22^ the G150T mutation constitutively shifts the helix to the relaxed conformation and crystallizes in a different space group than WT or G150A ICH (see Methods). As expected based on these structural and space group changes, G150T ICH superimposes onto WT and G150A ICH with a larger C_*α*_ RMSD of ~0.8 Å (see Fig. 3). In addition, the six WT and G150A datasets show ~2*σ* difference (mFo-DFc) electron density features around the mobile helix that indicate a minor population (< 10% occupancy) of the shifted helix conformation. Consistent with our efforts to minimize radiation damage to the crystals, these difference map features are much lower than those observed when Cys101 is oxidized to Cys101-SOH.^22^ These minor difference map peaks near the helix could indicate either the basal level of helical mobility in ICH or a response to minor X-ray-induced Cys101 modification in these datasets, possibly including thiyl radical formation.

**FIG. 3.**
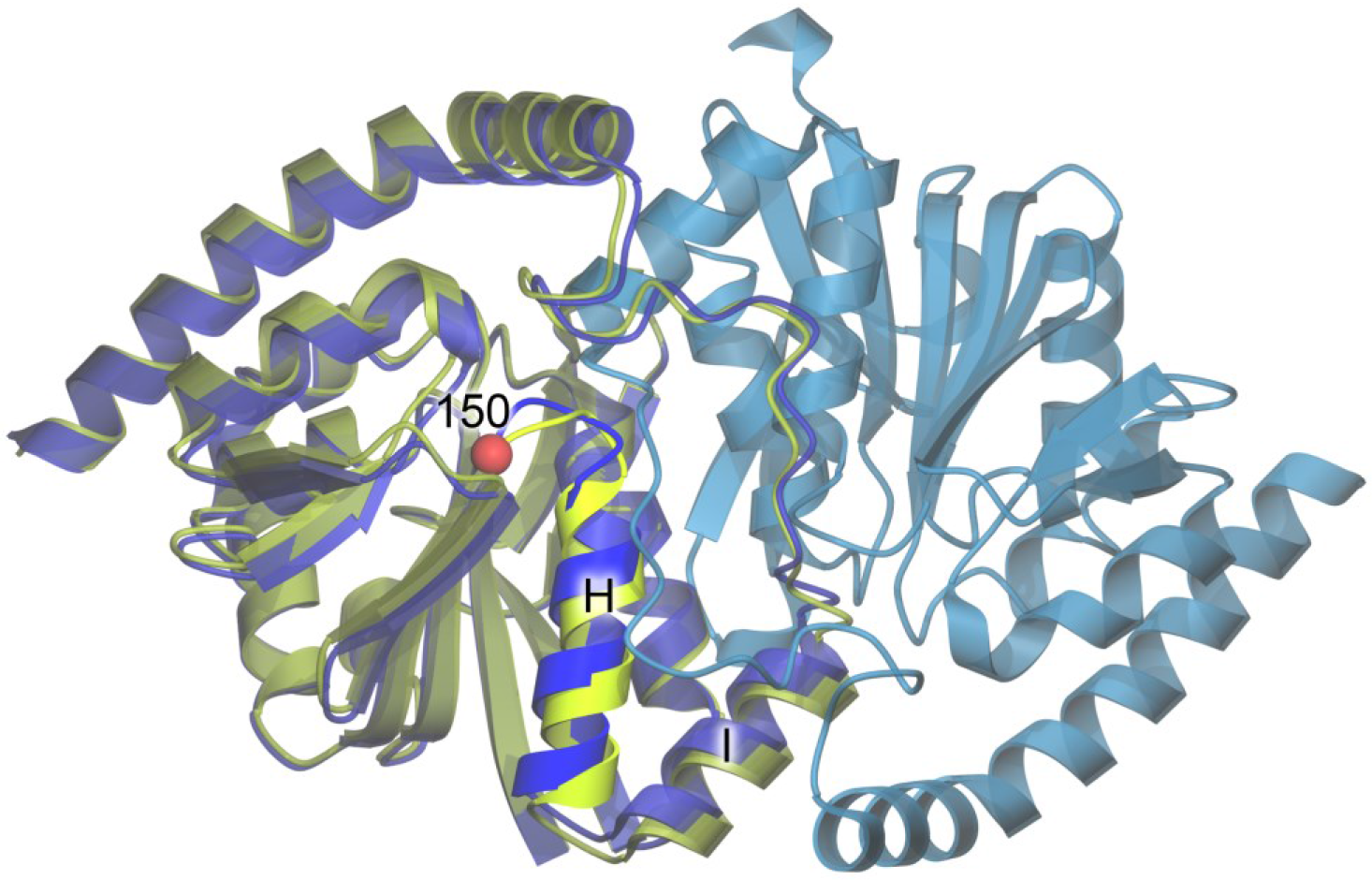
Structure of ICH. The ribbon diagram for the WT ICH dimer is shown in blue, with protomer A colored darker blue and protomer B lighter blue. The structure of G150T ICH (yellow-green) is superimposed on protomer A of WT ICH. The location of residue 150 is represented as a red sphere and the mobile helix is labeled H and shown in brighter colors.

### Quantifying the quality of the experimental diffuse scattering maps

Diffuse intensity is continuously distributed in reciprocal space and is weak compared to Bragg intensity; therefore, robust metrics for quantifying diffuse data quality are needed to avoid the introduction of noise or artifacts into the diffuse maps. Diffraction patterns were processed using our standard pipeline described in Methods to obtain 3D anisotropic diffuse scattering maps for all nine datasets. The diffraction volume was saved in a 3D lattice with 121×121×121 voxels sampled by integer Miller indices. The whole pipeline and visualization of each substep is displayed in Fig. 2. As shown in panel (F) of Fig. 2, anisotropic features were observable in processed diffraction patterns after the removal of Bragg peaks, although they were not as clear as those displayed in 3D diffraction volumes (panel (I)) after a deeper noise and isotropic component reduction. The average number of pixels that contribute to the intensity of each non-empty voxel in the diffraction volume is more than 1000 up to 1.4Å, as shown in Fig. S7, leading to a small standard error of the mean. In addition, the isotropic component is more than 10 times stronger than the anisotropic data (Fig. S8). Extracting large-scale anisotropic features from diffuse data therefore is challenging not only due to the high intensity of the Bragg peaks, but also due to the presence of a more intense isotropic component. The Laue-symmetrized anisotropic diffuse maps for all datasets are displayed as section cuts in Fig. 4 and 5 in the *q_y_* and *q_z_* directions, while other visualizations (in the *q_x_* direction) are shown in Fig. S9. Independent datasets of the same protein are very similar, as can be observed from their section cuts. This gives additional confidence that the diffuse maps produced by our pipeline contain *bona fide* protein diffuse scattering data and are not dominated by anisotropic background features or merging artifacts.

**FIG. 4.**
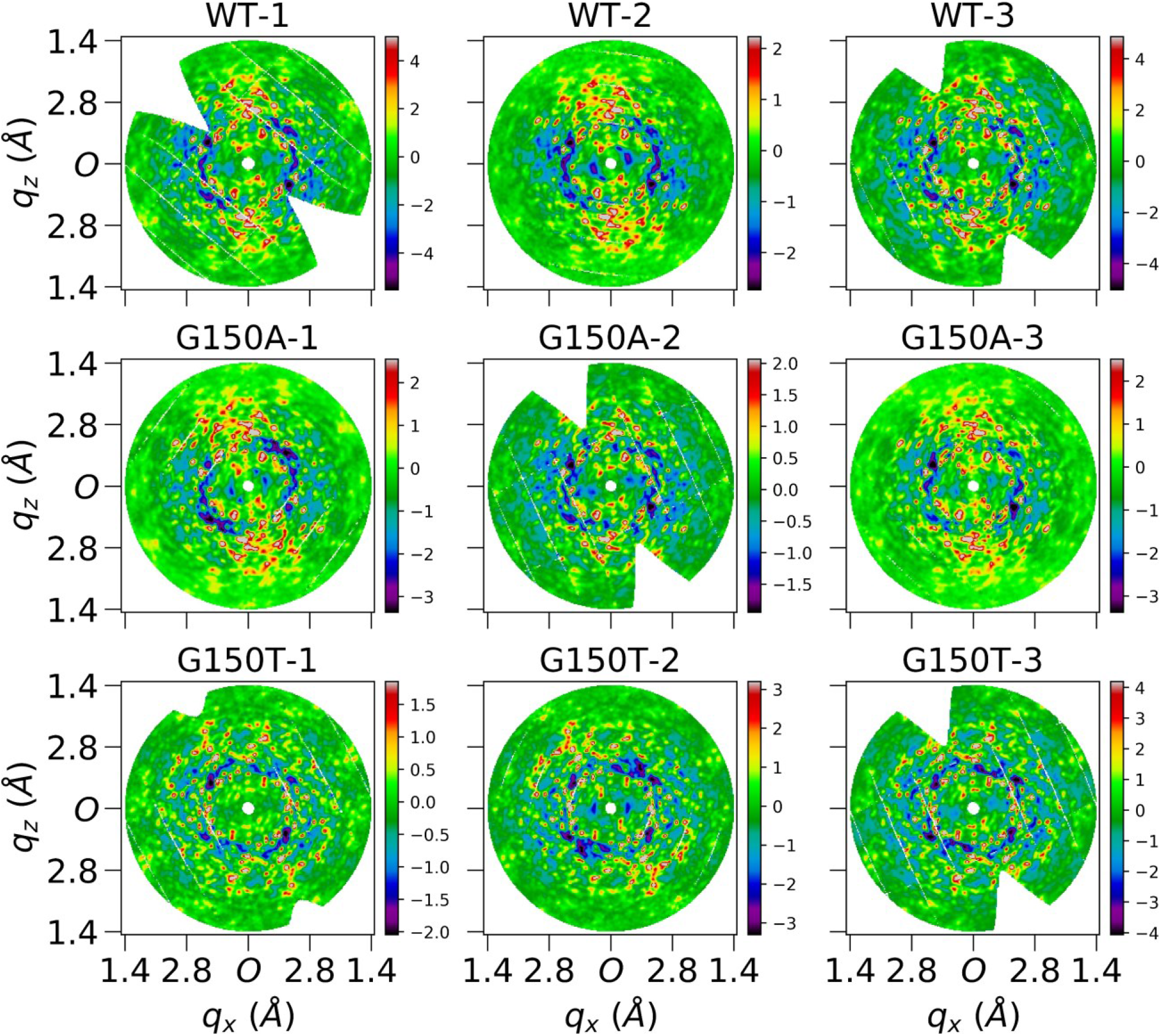
Central slices of Laue-symmetrized anisotropic diffuse maps (standard pipeline) of nine datasets perpendicular to *q_y_* direction. Each image is cut from the center of the corresponding diffuse map which is three-time finely sampled over Miller indices *H,K,L*. Each subfigure shows average voxels within a depth of 0.05Å^−1^ in *q_y_* direction, and 0.02 × 0.02Å^−1^ in *q_x_q_z_* plane. Both *q_x_* and *q_z_* axes extend to 1.4Å, and *0* represents the center in the reciprocal space. These finely sampled diffraction volumes were used for improved visualization only and were not used in data quality evaluation and modelling.

**FIG. 5.**
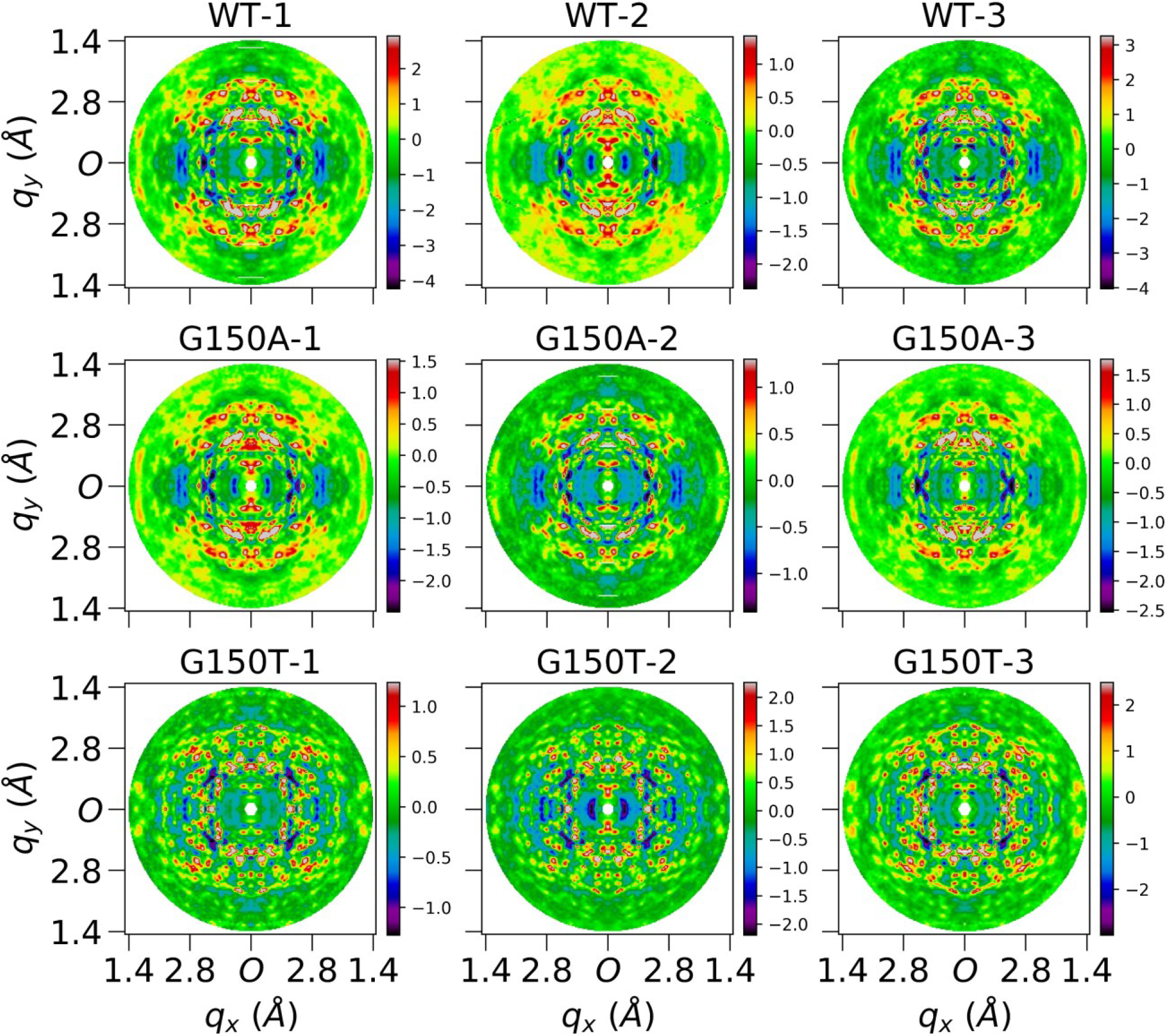
Central slices of Laue-symmetrized anisotropic diffuse maps (standard pipeline) of nine datasets perpendicular to *q_z_* direction. Each image is cut from the center of the corresponding diffuse map which is three-time finely sampled over Miller indices *H,K,L*. Each subfigure shows average voxels within a depth of 0.05Å^−1^ in *q_z_* direction, and 0.02 × 0.02Å^−1^ in *q_x_q_y_* plane. Both *q_x_* and *q_y_* axes extend to 1.4Å, and *0* represents the center in the reciprocal space.

In addition to using visual inspection, we assessed the quality of the extracted diffuse maps using quantitative metrics such as percent completeness, CC_Friedel_, and CC_Laue_ (Table 1). The resolution-dependent curves of these metrics up to 1.4Å are displayed in Fig. S10. Each dataset is > 95% complete in each resolution shell and > 98% complete over the entire resolution range. The CC_Friedel_ is > 0.7 in each resolution shell and > 0.9 overall. The CC_Laue_ is lower than CC_Friedel_, but it is still > 0.5 in each resolution shell and ≥ 0.85 in the overall resolution range. These numbers have been used to evaluate the data quality of diffuse maps before,^16,18,20^ however, in this work we find that the CC_Friedel_ and CC_Laue_ metrics are less sensitive to the data quality than CC_1/2_. For example, CC_1/2_ is roughly twice as sensitive as CC_Laue_ to changes in the diffuse map based on the observed decreases of both metrics in the analysis of different processing choices (Tables 2 and S6). In addition, as shown in Table 2, the CC_Laue_ is > 0.75 even without key processing steps such as the polarization correction or radial profile variance removal, where merging artifacts are clearly shown in section cuts of corresponding diffuse maps (Fig. S11 and S12). This suggests that CC_Laue_ fails to evaluate the data quality if there are contaminating background features in the images that roughly obey Friedel or Laue symmetry but are not the desired protein-derived diffuse signal. Based on these findings, we used CC_1/2_ to evaluate internal consistency of a diffuse map in this work, and increased emphasis on reproducibility to assess the data quality.

The CC_1/2_ values for each dataset are provided in Table 1, with the resolution-dependent curves shown in Fig. 6. CC_1/2_ varies from 0.76 to 0.89 for all datasets, indicating that the anisotropic diffuse features obey crystallographic point group symmetry reasonably well. CC_1/2_ is found to increase for the WT-1, WT-3, and G150A-2 datasets when the polarization correction is not used in the diffuse data processing pipeline (C in Table 2). This increase in correlation upon omitting an important correction is caused by the anisotropy in the diffraction pattern introduced by X-ray polarization that does not arise from the sample. Despite not representing crystal-derived diffuse features, these merging artifacts can greatly increase CC_1/2_ values when polarization-induced features happen to coincide with a crystal symmetry axis. As shown in Fig. S11, these polarization features are much stronger along some directions. Importantly, these artifacts are not reproducible between datasets, indicating that inter-dataset reproducibility may be a valuable additional data quality metric for diffuse scattering data.

**FIG. 6.**
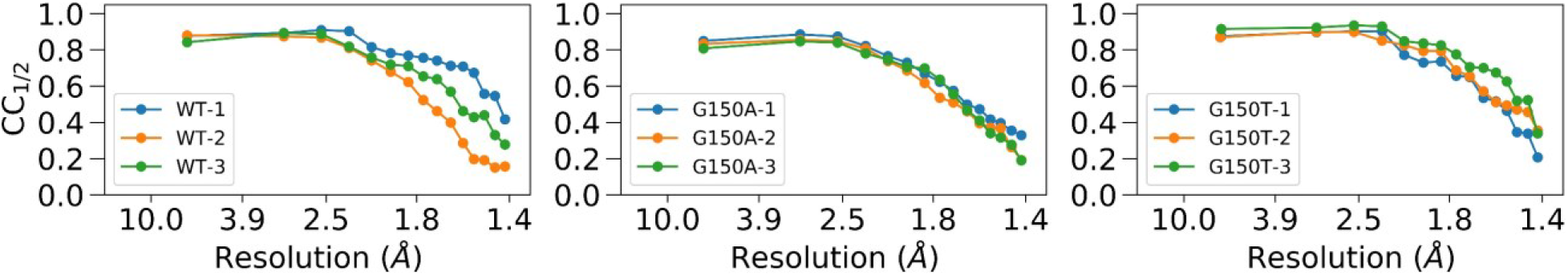
The resolution dependent CC_1/2_ curves for WT, G150A, and G150T datasets. Each curve was calculated using PHENIX up to 1.4Å, with the unsymmetrized anisotropic map as input.

Because anisotropic background features or artifacts can generate high values for CC_1/2_, another robust and unbiased quality metric for diffuse data is desired. To address this issue, we introduced CC_Rep_ as a measure of the reproducibility of anisotropic diffuse maps of the same protein collected from similar crystals. Collecting multiple datasets for the calculation of CC_Rep_ is not a large experimental burden, as PADs and shutterless data collection have reduced the time needed to collect a complete dataset to a few minutes at most synchrotron beamlines. The inter-dataset metric CC_Rep_ is valuable because it is not expected to be influenced as much as CC_1/2_ by artifacts or background scattering from the mount. Both metrics can be used together to increase confidence in the assessment of the quality of the anisotropic diffuse data. These two metrics also provide means to compare different data processing pipelines and to evaluate the effect of each submodule during processing, as we discuss below.

The CC_Rep_ statistics is summarized in Table 1, with the resolution dependent CC curves of dataset pairs of the same protein shown in Fig. 7. CC_Rep_ is > 0.8 for all datasets processed using the standard pipeline, and drops to lower values when important steps are omitted, as shown in Table 2. The detailed statistics of other diffuse data analysis choices is listed in Table S6. The CC_1/2_ value follows the same trend as CC_Laue_ although it is more sensitive to diffuse data quality, while the CC_Rep_ does not always follow the same trend as CC_1/2_. For example, the G150A-2 dataset processed without the polarization correction (C in Table 2) shows that its CC_Laue_ value increases by 0.07, and CC_1/2_ value increases by 0.13 due to the presence of merging artifacts with symmetrical features (Fig. S11). In contrast, CC_Rep_ decreases by 0.31, demonstrating that the improvement in CC_1/2_ might be due to background features or artifacts that are not reproducible in independent samples. Using the standard processing pipeline, all ICH datasets display substantial CC_1/2_ and CC_Rep_ values.

**FIG. 7.**
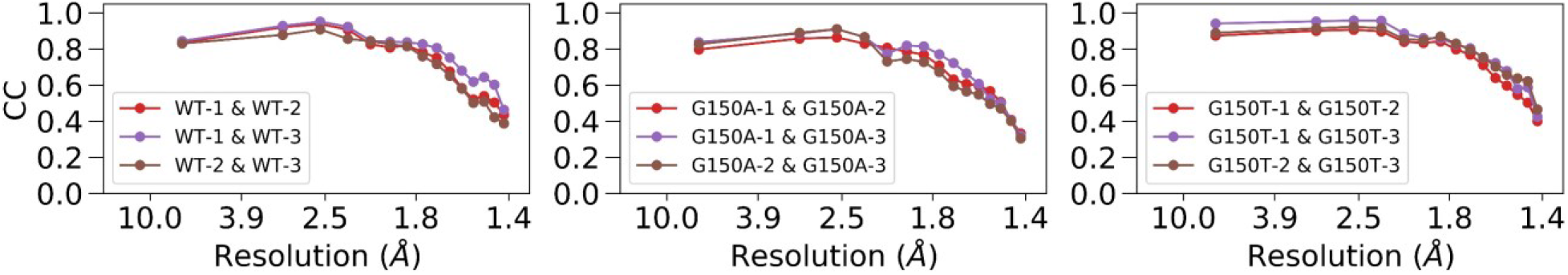
The resolution dependent CC curves of dataset pairs of the same protein. Each subfigure shows three CC curves between every two independent measurements for WT, G150A, and G150T, respectively. For example, the curve of WT-1 & WT-2 was calculated as the CC between Laue-symmetrized anisotropic diffuse maps of WT-1 and WT-2 datasets.

The near-identical WT and G150A ICH dimeric protein structures (C_*α*_ RMSD~0.06 Å) provide an opportunity to evaluate the cross-correlation coefficient (CC_Cross_) of their diffuse scattering maps. WT and G150A crystallize in the same space group, while G150T crystallizes in a different space group with a related cell to WT and G150A ICH (see Methods). The CC_Cross_ for WT-1, for example, can be calculated as the average CC of CC(WT-1, G150A-1), CC(WT-1, G150A-2), and CC(WT-1, G150A-3). We find that the CC_Cross_ is ≥ 0.83 for every WT and G150A dataset (Table 1), and each data pair within the set of replicate WT and G150A datasets also has CC ≥ 0.8, as shown in orange-colored cells in Table S5. The high cross correlation between WT and G150A diffuse datasets provides additional evidence that protein-derived diffuse scattering is the dominant feature in the processed diffuse anisotropic maps and is consistent with the minor differences in the crystal structures refined against the Bragg data.

### Evaluating models of protein motion using the LLM and RBT models

Much of the motivation for collecting diffuse data has been to develop models of correlated atomic motions. In this work, we develop LLM and independent RBT models as implemented in Lunus^17^ (see Methods). The traditional LLM model assumes that atomic motions in macromolecules have pairwise correlations that decay exponentially with a characteristic length *γ*, even across molecular and unit-cell boundaries.^9^ The magnitude of the atomic displacement is given by *σ*, which is refined as a single value for all of the atoms in the unit cell. In contrast, the RBT model assumes independent rigid body translation of the entire asymmetric unit.

Just as diffraction patterns can be mapped into reciprocal space to build 3D diffraction volumes, simulated diffraction images can be generated using diffraction volumes obtained either from experimental data or a model. This allows a direct visual comparison between the experimental and simulated diffraction patterns in the same orientation. One example is shown in Fig. S13, which compares the LLM model and the experimental data. Visual inspection of the simulated and experimental diffuse scattering shows agreement in many regions, although the simulated data display more detailed “granular” features, while the experimental data appear somewhat more “smeared”.

The individual atomic ADPs are set to zero in the standard LLM model (*I_0_*(***q***) in Eq. (2)).^16,17^ We wondered how well the diffuse data can discriminate between different models of atomic displacement, and whether using the refined ADPs (either isotropic or anisotropic) from the structural model might provide the LLM with a more accurate representation of variations in atomic positions in the protein. We therefore considered a variation of the standard LLM where *I_0_*(***q***) in Eq. (2) is replaced by *I_0_*(***q***), computed using either isotropic or anisotropic individual ADPs (Eq. (3) and Supplementary Material section IV). In addition to assessing the agreement with the data using the CC_LLM_, we considered whether the optimal values of *σ* were close to zero, consistent with the predictions of Eq. (3) (see Methods).

The results of the LLM analysis differed when using crystal structures refined using Refmac5 vs. PHENIX. For the Refmac5-refined PDB files, the LLM model parameters and CC_LLM_ for all ICH datasets using different ADP treatments are shown in Fig. 8 and summarized in Table S8; the resolution-dependent CC_LLM_ curves are shown in Fig. S14. In the case of the WT-1 dataset, the zero ADP model yields an overall CC_LLM_ of 0.67 to 1.4Å resolution, with a correlation length *γ* = 6.7Å and an overall atomic displacement *σ* = 0.40Å. The CC_LLM_ using the isotropic ADP model is higher (0.71), with a longer correlation length *γ* = 7.9Å and much smaller *σ* < 0.01Å, consistent with an overall Debye-Waller factor of unity as in Eq. (3). The anisotropic ADP model yields a value of CC_LLM_ that is comparable to the isotropic LLM (Table S8), despite being a superior model of the Bragg data. The other datasets show that the CC_LLM_ varies within 0.66-0.80 for the various ADP treatments. The highest CC_LLM_ is consistently achieved in the isotropic ADP LLM model, which varies within 0.70-0.80. The anisotropic ADP LLM model yields higher correlations than the zero ADP LLM model for all datasets. The correlation length *γ* is shortest (~ 7Å) in the zero ADP LLM model and longest (~ 8.5Å) in the anisotropic ADP model. We observe that the correlation length increases as the ADP model becomes more detailed in most (seven) datasets in this work.

**FIG. 8.**
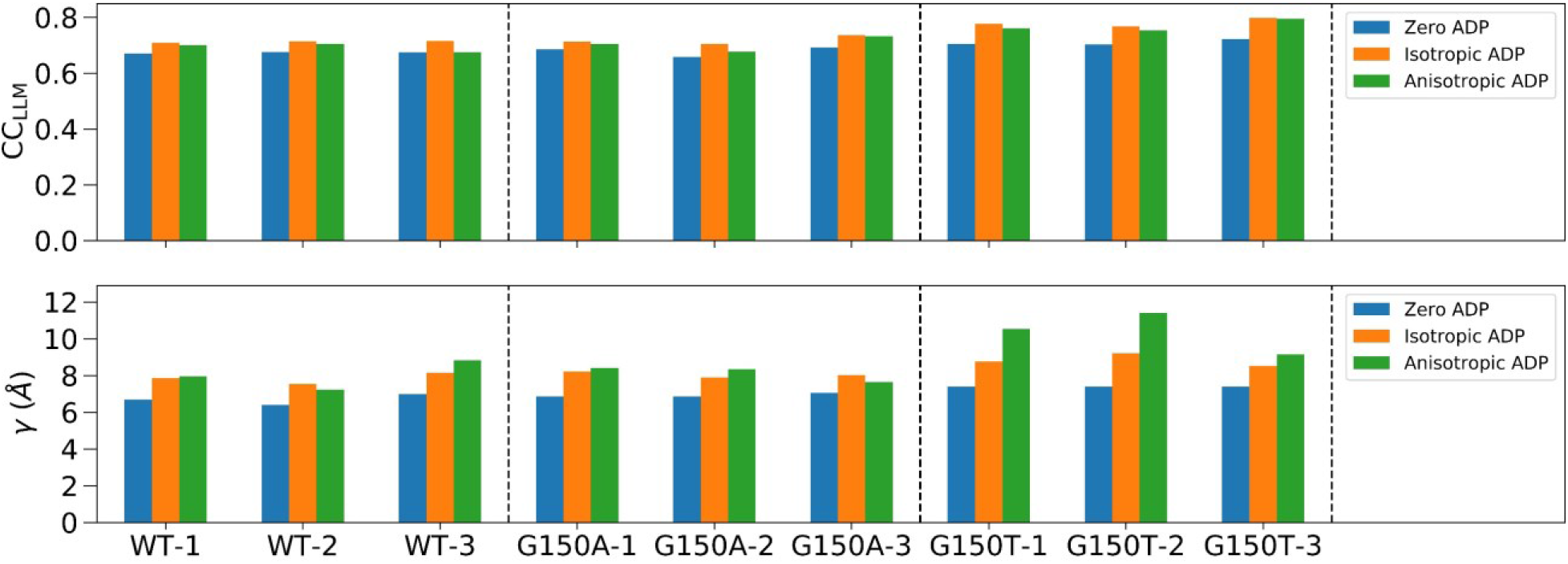
The LLM model statistics for all ICH datasets using Refmac5-refined PDB files with different ADP treatments. The two subfigures display the best-fit CC_LLM_ and average correlation length *γ*, respectively. Each dataset was analyzed using three different ADP models including the zero, isotropic, and anisotropic ADP, respectively. The dashed vertical line separates WT, G150A, and G150T datasets. The full LLM statistics are presented in Table S8.

Despite the Refmac5 and PHENIX models having comparable model statistics and agreement with the Bragg data, the PHENIX models have different distributions of ADP anisotropy (Fig. S1). In particular, for the WT PHENIX models, the distribution deviates from the “bell-shaped” distribution centered on ~0.45 that is typically observed in proteins (Fig. S1).^39,40^ In contrast, the Refmac5-refined models have anisotropy distributions that are closer to the average of other proteins, with fewer extreme anisotropy values (Fig. S1). The differing anisotropy values are not correlated with changes in the overall magnitude of the PHENIX- and Refmac5-refined ADPs, which are highly similar (Fig. S2). We determined that the difference in the anisotropic ADPs is due to different overall anisotropic scale parameters produced by the two programs (see Methods, Table S4, and Supplementary Material section III). We were able to use these different anisotropic scale matrices to convert the PHENIX-refined anisotropic ADPs into ones that closely resemble those in the Refmac5-refined model and vice versa (see Methods; Fig. S1, S3, S4, S5; Supplementary Material section III), confirming that the differences in the PHENIX and Refmac5 anisotropic ADP models are due predominantly to different anisotropic scaling parameters. This does not exclude the possibility of residual anisotropic ADP differences arising from different restraints in the two programs, which might be important for solvent atoms (see Fig. S1, S4, S5).

Although the different ADP models agreed equally well with the Bragg data (Table S1), this was not the case for the diffuse scattering data. Results of the LLM analysis using either the Refmac5- and PHENIX-refined input models are summarized in Tables S8 and S9. These two sets of models are comparable for all ADP treatments except anisotropic ADPs, which show marked differences. In general, the CC_LLM_ values are higher and *σ* values are lower for the Refmac5 anisotropic ADP models compared to those refined in PHENIX. The discrepancies between the Refmac5 and PHENIX models are clearest for the three replicate WT datasets, where the agreement with the data is lower for the PHENIX anisotropic ADP models (CC_LLM_ ~ 0.6) than the Refmac5-refined models (CC_LLM_ ~ 0.7); the PHENIX models also lead to higher *σ* values in the best-fit LLM (~ 0.2Å), suggesting an inconsistency with the predictions of Eq. (3). The difference in mean CC_LLM_ and *σ* between the Refmac5 and PHENIX models are larger than their standard deviations across three replicate WT datasets, supporting the significance of the discrepancies. However, the higher *σ* value for the WT-3 dataset indicates that there might be issues that remain in that Refmac5 anisotropic ADP model, or, alternatively, that there might be issues with the WT-3 diffuse data.

Compared to the low sensitivity of the Bragg data to anisotropic ADP differences as judged by the similar R_free_ values for the PHENIX and Refmac5 models (Tables S1, S10), the increased sensitivity of the diffuse data suggested that diffuse scattering might potentially be useful for modeling ADPs. However, the R factors are computed in a different way than CC_LLM_ and these two statistics are not directly comparable to each other. We therefore used a measure of the agreement with the Bragg data -- CC_Bragg_ -- that is computed in the same way as CC_LLM_, except using Bragg data. Specifically, CC_Bragg_ was computed as the Pearson correlation between the model and Bragg data intensities (as opposed to amplitudes) after subtracting the isotropic component, and, importantly, without applying overall anisotropic ADP scaling. Therefore, CC_Bragg_ and CC_LLM_ provide quantitatively comparable measures of model quality that can be used to assess the relative sensitivity of Bragg and diffuse scattering data to these different anisotropic ADP models. We compared CC_Bragg_ and CC_LLM_ values obtained for the Refmac5 and PHENIX models as well as the Refmac5 and PHENIX models that had been rescaled using the difference anisotropic scaling matrices (see above; Methods). The results are summarized in Fig. 9 and Table S11, and clearly indicate that the diffuse data are more sensitive to the differences in the ADPs than the Bragg data. For example, whereas CC_LLM_ for the PHENIX WT-1 model increases from 0.6 to 0.7 after ADP rescaling, the CC_Bragg_ value changes by a much smaller amount, from 0.893 to 0.895.

**FIG. 9.**
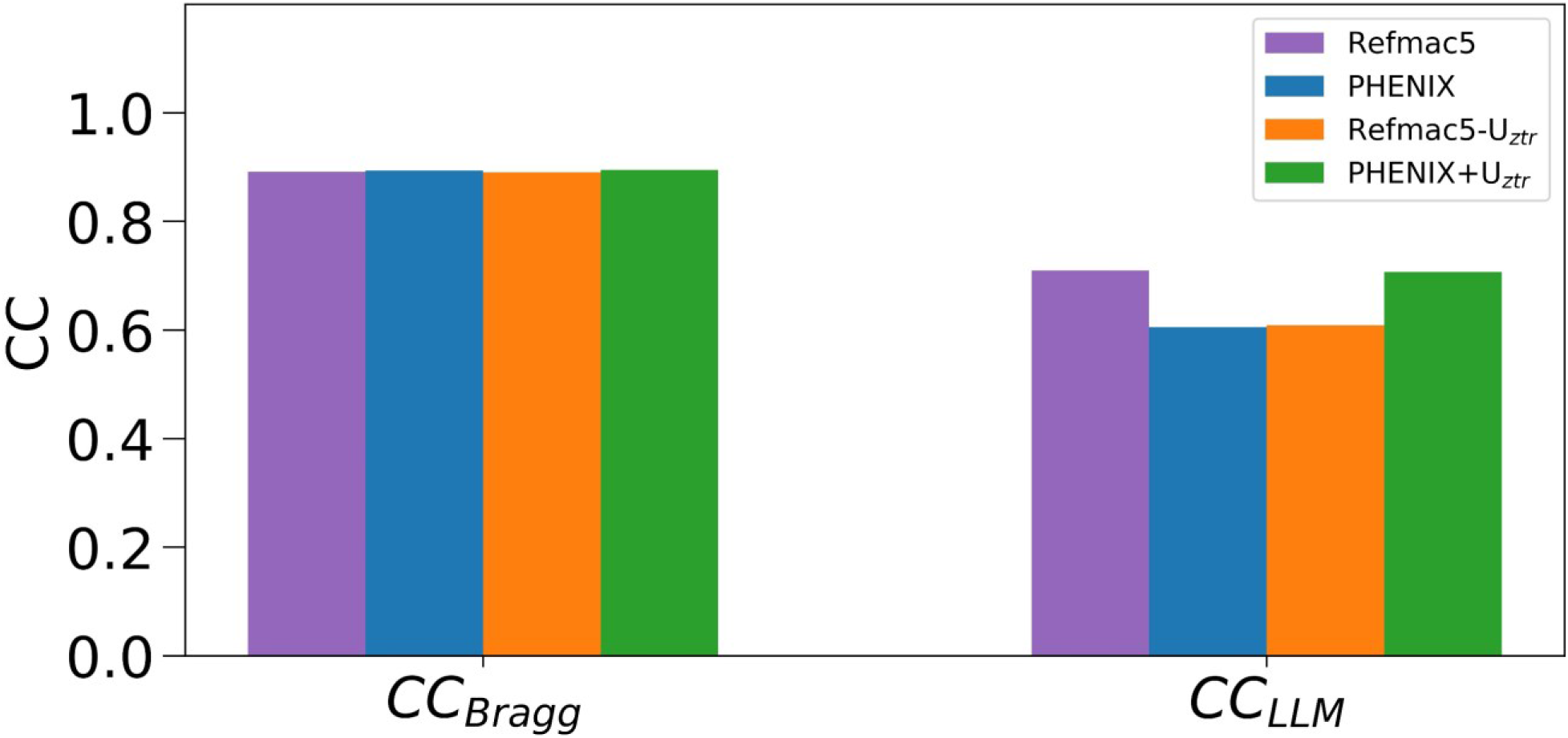
CC_Bragg_ and CC_LLM_ between experimental data and calculated Refmac5, PHENIX models with and without rescaling of the model B factors using Uztr for the WT-1 dataset. The sensitivity of CC_LLM_ to changes in the ADPs is much greater than that of CC_Bragg_, indicating that diffuse scattering data are more sensitive than Bragg data to anisotropic scale factor-related changes in ADPs. The displayed CC values are calculated with a low resolution cutoff of 10 Å because no bulk solvent correction was used. Both the Bragg and diffuse intensities have had the isotropic component removed as described in Methods.

Considered together, the improved anisotropic ADP CC_LLM_, the lack of change in CC_Bragg_ (Fig. 9 and Table S11), the lower values of *σ*, and the more typical distribution of anisotropies for the Refmac5-refined and rescaled PHENIX anisotropic ADP models show that diffuse scattering data favor anisotropic ADP models that possess more plausible features even when the Bragg models have similar R_free_/R_work_ and CC_Bragg_ values. The implications of this observation for using diffuse and Bragg data together to refine crystallographic models are discussed below.

Some studies have indicated that independent rigid-body motions of macromolecules are responsible for a significant portion of the diffuse scattering signal.^4,34^ To investigate this possibility for ICH, we implemented an independent RBT model in Lunus and used a metric equivalent to CC_LLM_, called CC_RBT_, as a target for optimization.^17^ The optimal CC_RBT_ and displacement parameter (*σ*) values are summarized in Table S12. The CC_RBT_ (~ 0.55) is lower than CC_LLM_ by about 0.1 for all datasets and ADPs treatments. The optimal *σ* values in the zero ADP RBT model are generally similar to those in the zero ADP LLM model. Interestingly, as in the LLM models, using the Refmac5-refined anisotropic ADP models (Table S12) produces higher CC_RBT_ values than the PHENIX-refined models (Table S13), although their differences are not as large as those for the LLM model.

### Studying the effects of various steps in the analysis pipeline

To determine which aspects of the diffuse scattering experiment and subsequent image processing have the greatest impact on final data quality, we systematically omitted each step in our pipeline, one at a time. Results of this analysis are partially shown in Table 2 and summarized in Table S6. Data quality assessed using CC_1/2_ and CC_Rep_ does not change greatly when the solid-angle, detector absorption, and parallax correction are omitted. In contrast, omitting the non-crystal background subtraction, polarization correction, or radial profile variance removal step substantially degrades the data quality. The omission of non-crystal background subtraction reduces the two quality metrics by 0.02-0.08 for all datasets, with the visualization only changed slightly (Fig. S15). The omission of the polarization correction reduces CC_Rep_ of all datasets by more than 0.1, with CC_1/2_ varying in a less informative way for each dataset. The omission of radial profile variance removal step reduces both quality metrics by 0.01-0.09 for most datasets and decreases a few of them by more than 0.1. The significant effects of these three steps are expected, as they are critical to remove contaminating anisotropic background intensity and to reduce merging artifacts (Fig. S11, S12). In contrast, other processing steps, such as the solid-angle, detector absorption, and parallax correction, only affect the radial intensity distribution in the diffraction pattern but do not introduce angular anisotropies. In addition, the omission of polarization correction increases the CC_1/2_ value because of strong anisotropic artifacts (see Fig. S11) that are introduced by X-ray polarization are not removed. In this case, also omitting the solid-angle correction can scale down the contribution of high resolution data to the calculation of correlations, leading to slight improvements for both CC_1/2_ and CC_Rep_ in the overall resolution range.

For the study of four different scale factors, the radial profile, overall, and water ring scale factors follow the same trend and only vary slightly (Fig. S16). However, the Bragg scale factor is significantly different from the other three, especially in the last half of each dataset where it increases more than the others (Fig. S16). This means that the last half of the images will be scaled to a much higher intensity level using the Bragg scale factor. The data quality metrics using each scale factor treatment are summarized in Table S7, where the radial profile variance removal step is turned on for (A)-(D) and turned off for (E)-(H). The radial profile, overall, and water ring scale factors with radial profile variance removal (A)-(D) produce the same CC_1/2_ and CC_Rep_ for all datasets, while the Bragg scale factor performs slightly worse. However, when the radial profile variance removal step is turned off (E)-(H), all four scale factors perform much worse, with the Bragg scale factor treatment producing very poor diffuse maps that are dominated by merging artifacts, as shown in Fig. S17. Interestingly, only the Bragg scale factor (H) has a measurable effect on the CC_LLM_ even though the data quality as quantified by CC_1/2_ and CC_Rep_ significantly decreases for other processing choices (D)-(G).

## IV. DISCUSSION

### Using multiple quality metrics to produce high quality diffuse scattering maps

Reliably extracting the relatively weak diffuse scattering signal from raw diffraction images is vital for generating useful diffuse scattering maps for downstream applications. Several different data quality metrics have been discussed in this article, including CC_Friedel_, CC_Laue_, and CC_1/2_ for evaluating internal consistency in diffuse datasets, and CC_Cross_ and CC_Rep_ for measuring inter-dataset reproducibility. CC_Laue_ and CC_Friedel_ evaluate whether the diffuse map follows the expected symmetry but they have behaviors that make them less desirable as data quality metrics. In particular, CC_Laue_ values change by about half as much as CC_1/2_ values when perturbations to the data processing are introduced (Tables S6, S7). Moreover, the value of CC_Laue_ can be rather high even for a diffuse map with obvious merging artifacts (see Fig. S11). This is in part because each voxel in a symmetrized map contains a contribution from the corresponding voxel in the unsymmetrized map, leading to a nonzero correlation even for random datasets. The correlation is highest for low-symmetry Laue groups: in P1, where CC_Laue_ corresponds to CC_Friedel_, the value is about 0.7 for a random dataset. Because of this, we favor CC_1/2_ as a quality metric.

Despite the benefits of using CC_1/2_ to assess data quality, a symmetry measure alone cannot fully describe the data quality of a diffuse map, especially when the map is dominated by anisotropic background features or artifacts which may approximately obey these symmetries. This consideration motivated our use of the metric CC_Rep_ to validate whether the anisotropic diffuse signal originates from protein crystal diffraction. The paucity of data quality metrics that can discriminate between anisotropic diffuse scattering from the sample and from the background is an important reason that different protocols for constructing diffuse maps have been reported.^16,20,34^ The combined use of CC_1/2_ and CC_Rep_ provides a more complete picture of data quality than CC_1/2_ alone. The use of these metrics also helped to assess quantitatively the performance of our data processing pipeline and enabled the processing choice analysis in this work. The CC_Cross_ is a special metric that can be used for two proteins with similar structures and unit cell dimensions, such as WT and G150A ICH in our experiment. It may have particular value when comparing changes in diffuse scattering between similar samples that have been subjected to perturbations such as temperature change, mutation, etc.

### Effects of each processing step in the diffuse map construction pipeline

Constructing and modeling the 3D diffuse map is now the standard method for diffuse scattering analysis. Although different versions share a similar general workflow, the details may vary. Benefiting from the introduction of two additional quality metrics, we are able to perform a detailed analysis of the variations, yielding insight into the impact of each processing step on the overall quality of the anisotropic diffuse map (Tables 2, S6, S7). The standard pipeline works satisfactorily for all datasets giving CC_1/2_ ≥ 0.76 and CC_Rep_ ≥ 0.81. Eliminating the parallax, solid-angle, and detector absorption corrections have small effects on both CC_1/2_ and CC_Rep_, perhaps related to the fact that they only modulate the radial intensity distribution in the diffraction pattern. In contrast, the non-crystal background subtraction, polarization correction, and radial profile variance removal have stronger effects on the data quality of extracted diffuse maps. The omission of these steps will affect the angular intensity distribution in the diffraction pattern and introduce strong artifacts or anisotropic background features to the diffuse map, which leads to systematic errors in the anisotropic diffuse intensity. It is important to note that the non-crystal background image subtraction requires acquiring matched background exposures at the time of data collection. The collection of non-crystal background patterns has not been consistently performed until recently^21^ despite its simplicity. When a shadow from the capillary or beamstop is visible it can be manually masked out from the detector image, but other anisotropic noise may not be visible by eye in the single diffraction pattern and thus can accumulate in the 3D diffuse map. We suggest collecting non-crystal background patterns in rotation method experiments. For SFX experiments, it might be possible to improve data quality by analyzing non-hit patterns and finding suitable background patterns for subtraction. The radial profile variance removal is another important step to avoid introducing merging artifacts in the diffuse map (Fig. S12). An alternative^34^ to radial profile variance removal is to subtract the radially averaged profile from each diffraction pattern before the 3D merging step; indeed, in implementing our removal method, we found that the difference in image radial profiles is similar to the first principal component. The diffuse map construction pipeline is flexible to some extent and the main focus is to remove anisotropic noise and avoid merging artifacts. Any steps that can introduce errors in the angular intensity distribution in the diffraction pattern deserve careful attention.

In addition to the processing choice analysis, four different types of per-image scale factors were also evaluated by comparing the data quality of diffuse maps processed by corresponding scale factors. As shown in Results, the radial profile, overall, and water ring scale factors generate similar results according to our data quality metrics, and perform moderately better than the Bragg scale factor which was adopted similarly by Peck *et al.*^20^ for systems other than ICH, using scale factors from XDS. When the radial profile variance removal step is turned off, all four scale factors give much worse results than the standard pipeline and also perform differently, although Fig. S16 shows that curves of the radial profile, overall, and water ring scale factors only vary slightly for all datasets. This indicates that data quality of the diffuse map is very sensitive to changes in the scale factor when radial profile variance removal is absent, while the radial profile variance removal step greatly reduces the impact of scale factors. In any case, for our ICH data, the Bragg scale factor always behaves worse than the others, which can be inferred from its distinctive curve that differs from the others especially for the last half images of each dataset. The Bragg scale factor increases to higher values than the other three scale factors and this is probably induced by the decrease of Bragg intensities in the last half diffraction patterns. The radial profile scale factor therefore is preferred for extracting high-quality diffuse maps from ICH diffraction images.

### Analysis of diffuse scattering using the LLM model

Using the standard LLM model with zero ADPs^16,17^ (Eq. (2)), the agreement with the data (CC_LLM_ of ~ 0.7), the value of the atomic displacement *σ* (~ 0.4Å), and the value of the correlation length *γ* (~ 7Å) are all comparable to previous studies of other protein crystals using coarse-grained diffuse data.^16,18,36^ Using isotropic ADPs in the calculation of *I_0_*(***q***) in Eq. (2), the optimal LLM models yielded slightly higher correlations with the data than using zero ADPs, and the differences exceed the standard deviations of three replicate datasets for all protein forms. The CC_LLM_ of 0.80 for G150T-3 is in the high end compared to the correlations reported from some previous work.^16,18,36^ The fitted values of *σ* for this model are very close to zero, indicating that the ADPs from the Bragg analysis are consistent with the pattern of diffuse intensity predicted by Eq. (3). The fact that including isotropic ADPs in the LLM leads to a low value of *σ* lends additional support to the utility of using a LLM model to analyze the ICH diffuse data. We consistently found that the refined correlation lengths *γ* were longer for the isotropic (~ 8Å) and anisotropic ADP models (~ 8.5Å) than in the zero ADP model (~ 7Å) for all nine datasets. The dependence of the correlation length on the complexity of the atomic displacement model was unexpected. However, we note that the LLM used here involves only a single correlation length, whereas it is more likely that displacements with multiple correlation lengths contribute to the actual diffuse signal.^11^

Because atomic motions result in the loss of Bragg intensity and increased diffuse scattering, there has been long-standing interest in combining Bragg and diffuse scattering data to improve models of atomic motion in crystal structures.^2,6,41^ By using LLM models that incorporate different anisotropic ADP models for the same structural model, we found that diffuse scattering data can discriminate between more and less plausible representations of anisotropic atomic motion, even when these models have similar R_free_/R_work_ and CC_Bragg_ values and thus cannot be distinguished easily based on Bragg data alone. Both Refmac5- and PHENIX-refined models agree well with the Bragg data, however the PHENIX models consistently refine to lower anisotropy values (corresponding to more anisotropic motion) than the Refmac5 refinements (see Tables S2, S3) and sometimes have anisotropy distributions that deviate from the “bell-shaped” curve centered on ~ 0.45 that is typically observed (Fig. S1).^39,40^ We showed that the origin of this effect is that these two widely used refinement programs can produce different anisotropic scaling parameters even when the starting model and the datasets are identical. This results in different anisotropy in the final model ADPs, even though the ADP magnitudes (i.e., B_eq_) are nearly identical. This difference is understandable because the total anisotropy in the diffraction data contains contributions from the crystal as a whole (anisotropic scaling parameters) and from individual atomic motions (ADPs), whose values are highly correlated and thus they are refined separately.^42^ Therefore, if different anisotropic scale parameters are initially refined by different programs using otherwise identical starting models and datasets, there will be subsequent compensatory changes in the refined anisotropic ADPs of the final models, as we have observed. In addition, we find that when ICH LLM models that already include individual ADPs also have substantial *σ* values, the models tend to agree less well with the diffuse data; it is possible that LLM analysis of *σ* values might be used for other systems as a general indicator of when ADPs deserve additional scrutiny. Interestingly, LLM models with anisotropic ADPs have CC_LLM_ values that are comparable to or lower than models using isotropic ADPs. The lack of improvement going from the isotropic to anisotropic ADP model was unexpected because anisotropic ADPs contain information about both the preferred directions and amplitudes of motion and substantially improve the agreement of the refined models with the Bragg data (see Methods; Table S10). While there are several lines of future investigation suggested by our results, the ability of diffuse scattering data to discriminate between models of anisotropic atomic motion that are equally consistent with the Bragg data indicates that joint refinement of models against Bragg and diffuse scattering data -- an idea long discussed in the literature^41^ -- is promising and might result in more accurate representations of atomic motion in proteins. We note that because ICH exhibits controllable concerted helical motion, it makes an ideal system in which to explore the ability of diffuse scattering data to discriminate between various representations of correlated secondary structure motions in the future.

Recent articles^4,34^ have suggested that independent rigid-body translations, like those in our RBT model, are responsible for the majority of the diffuse signal in protein X-ray diffraction. For ICH, we found that the LLM model agrees better with the diffuse data distributed between the Bragg peaks than the RBT model for all datasets in all ADP models (CC_LLM_ and CC_RBT_ values in Tables S8 and S12). This result indicates that the large-scale diffuse features in ICH are more accurately described using liquid-like rather than independent translational rigid-body motions. As the values of *γ* from the LLM fits are much smaller than the size of the protein, our results suggest that the correlation lengths inherent in the RBT model might be too long. Note that we did not consider rigid-body rotations, and that our findings do not exclude the possibility that rigid-body motions coupled across molecular and unit-cell boundaries are important for modeling the sharper diffuse features in the neighborhood of the Bragg peak.^21^

It is important to interpret data quality metrics (such as CC_1/2_ and CC_Rep_) and model quality metrics (CC_LLM_, CC_RBT_) in their appropriate contexts. Data quality metrics pertain only to the measured signal and are independent of model quality metrics, which quantify agreement between a representation of the data and the measured signal. However, better data processing approaches are expected to result in more accurate models. A prominent example is the development of paired model refinement in concert with CC_1/2_ for processing Bragg data, which uses the model R_work_ and R_free_ values obtained from refinements against datasets processed to different resolution limits in order to determine the maximal resolution at which meaningful signal is present.^23^ Although we did not use a full paired refinement-like workflow, we found that the CC_LLM_ values for the refined LLMs were not sensitive to even serious degradation in the quality of the diffuse maps, unlike the data quality metrics CC_1/2_ and CC_Rep_. For example, CC_LLM_ does not change significantly even when the diffuse data quality is severely reduced, such as in the WT-3 dataset processed without the radial profile variance removal step (D in Table 2). In this case, the data quality as quantified by CC_Rep_ decreases by 0.11 while CC_LLM_ does not change. Therefore, we do not currently recommend using model CC values as a metric for evaluating diffuse scattering data processing decisions, although this may change with improved models of correlated motions.

### Lessons about experimental best practices for the collection of macromolecular diffuse scattering data

Our detailed analysis of the influence of various processing steps on the quality of diffuse maps provides insights into important experimental aspects of collecting diffuse scattering data. The weak intensity values of diffuse scattering compared to Bragg diffraction places a premium on experimental approaches that reduce background scattering,^21^ and our results underscore the importance of careful treatment of the background. Because the speed of modern data collection makes collecting multiple datasets straightforward, we suggest collecting non-crystal background images which, in the case of a rotation series, match the spindle angles of the crystal exposures. There is broad agreement that the sample-derived signal should be maximized by using large crystals and by reducing sources of scattering in the beamline setup. However, the best choice of the sample mount is still debated. In this study, we used thin-walled borosilicate glass capillaries that are expected to have nearly isotropic background scattering. However, glass scatters X-rays ~10 times more strongly than plastics such as kapton,^43^ and thus will produce an intrinsically higher background that obscures weak diffuse scattering signals. In addition, depending on the diffracted beam path through the capillary walls, the greater absorption of glass might lead to anisotropy in the absorption of scattered X-rays. While most plastic mounts enjoy the advantage of lower scattering, they generate an anisotropic background owing to scattering by partially oriented molecules that compose the plastic. Our work and those of others^20,21^ indicate that combining the collection and careful subtraction of background non-crystal images with PCA analysis allows for effective removal of contaminating anisotropic background signals; however, a model of the capillary would be required to account for anisotropic absorption effects. This suggests that plastic capillaries with lower scattering may be preferable for diffuse scattering experiments despite their more anisotropic background. An important consideration with plastic capillaries is that the loop that is typically used to support the crystal in these mounts can generate a large anisotropic background signal. Therefore, it is advisable to use a loop that is smaller than the crystal and to aim the X-ray beam into portions of the crystal that are fully outside the loop throughout the entire rotation range. This is important because it is difficult to collect well-matched non-crystal background images that include empty loop scattering for later subtraction from the diffraction images.

Prior diffuse scattering work has used large, well-diffracting crystals with comparable thickness in all three dimensions.^6,16,19,21^ Such crystals are advantageous for diffuse scattering because they place comparable volumes of the crystal in the X-ray beam in all orientations during data collection, resulting in images with similar diffraction intensity throughout the dataset. In contrast, WT ICH crystals grew with a difficult, plate-shaped habit that required careful mounting in order to orient the short axis of the plate co-linearly with the capillary axis so that the X-ray beam illuminated similar thicknesses of crystal during rotation. Our initial inspection of diffuse data collected from crystals that were not so carefully oriented indicated that the data quality suffered when the X-ray beam illuminated very different thicknesses of the crystal during data collection. We note that rods do not present this problem so long as the long axis of the rod is roughly collinear with the rotation axis, which is their naturally preferred orientation during capillary mounting. Although it is clear that diffuse scattering researchers previously appreciated the importance of crystal size and shape for data quality, crystal morphology should be considered by experimentalists when planning a diffuse scattering experiment, particularly if plate-shaped crystals are being used.

Our use of the reproducibility metric CC_Rep_ showed that there was a much larger amount of contaminating anisotropic intensity in the WT-3 dataset compared to the other two replicates, which may not have been obvious had we not collected the other two datasets for comparison. The radial profile variance removal approach was able to suppress these problematic features and resulted in a usable final dataset that compared well with its replicates after processing based on the quality metrics. However, the LLM model of WT-3 still stands out as an outlier with a much larger *σ* in the anisotropic ADP model. The absence of comparable contaminating anisotropic features in WT-1 and WT-2 excludes beamline components, detector issues, or other sources that would be common to all three datasets. It is possible that the culprit is contaminating detritus (e.g., lint, a fiber from the wick, etc.) that may have adhered to the crystal used to collect the WT-3 dataset during mounting. This illustrates the sensitivity of diffuse scattering data to minor sources of non-crystalline scattering that make a negligible contribution to the Bragg data and demonstrates the value of collecting multiple datasets.

The intrinsic weakness of diffuse scattering data presents detection challenges that are tempting to solve by increasing the X-ray dose. However, because diffuse scattering data are typically collected from crystals at ambient (i.e., non-cryogenic) temperatures, radiation damage is a major concern. In this regard, the ICH system was especially valuable, as it contains a radiation-sensitive active site cysteine nucleophile (Cys101) that is readily photooxidized to cysteine-sulfenic acid at X-ray doses lower than the typically quoted 3×10^5^ Gy dose limit for ambient temperature Bragg data collection.^22,44^ We did not see strong evidence of Cys101 oxidation in these datasets, although we cannot exclude that some minor oxidation occurred. The minimal radiation damage in these sensitive crystals indicates that PADs, rapid shutterless data collection, and the use of large beams (~100-200 μm) can limit radiation damage and allow the collection of usable diffuse scattering data from moderately radiation-sensitive protein crystals. As in prior work,^19^ we collected usable Bragg and diffuse scattering data simultaneously, and it is possible that such combined Bragg/diffuse datasets could be used for the global refinement of macromolecular structure, atomic mobility, and correlated motions in the future.

## V. CONCLUSION

In this work, we have developed an open-source data analysis pipeline *dspack* to extract diffuse scattering features from X-ray diffraction patterns. Detailed studies were performed to validate the effectiveness of this pipeline and demonstrate how each submodule and different analysis variables can affect the data quality of extracted diffuse maps. We described our systematic study of the reproducibility of diffuse scattering from isocyanide hydratase (ICH) with nine datasets of three different protein forms demonstrating that the replicate diffuse datasets were similar in pairwise comparisons (Pearson correlation coefficient (CC) ≥0.8). In particular, these studies emphasized the importance for data quality of non-crystal background pattern subtraction, radial profile variance removal of radial intensity profiles, and the approach to calculating per-image scale factors. We introduced two unbiased and robust metrics (CC_1/2_ and CC_Rep_) to evaluate the data quality of diffuse maps. We conclude that using CC_1/2_ alone can lead to artificially high assessments of data quality, and that including CC_Rep_ can help to obtain a more reasonable assessment of data quality. We found that diffuse scattering data are more sensitive than Bragg data to different models of anisotropic atomic motion resulting from distinct anisotropic scaling parameters, and that diffuse scattering data favor models with more typical distributions of atomic anisotropy. In a comparison of the LLM and independent RBT models of protein motions inside the ICH crystal, we found that the agreement with the data is higher for the LLM model than for the RBT model, and that the LLM model agreement is in the high end among those reported in some other studies.^16,18,36^ Overall, this study provides a new set of computational tools for the analysis of diffuse scattering data, demonstrates the potential value of diffuse scattering for evaluating some types of ADP models, and indicates that ICH is an excellent system for future diffuse scattering studies.

## SUPPLEMENTARY MATERIAL

See the supplementary material for additional figures, tables, and detailed descriptions of the individual B factor LLM model.

## ACKNOWLEDGMENTS

The authors are grateful to an anonymous referee for comments that led to substantial improvements in the manuscript, and insights that enabled us to determine the source of the difference in ADPs between the PHENIX and Refmac5 models. Z.S. would like to thank Prof. Mike Dunne for his guidance and support, and Prof. James Holton for the discussion on X-ray cross-sections of glasses and plastics. M.A.W. acknowledges support from NIH R01GM139978.

H.v.d.B. was supported by NIH R01GM123159, and by a Mercator Fellowship from the Deutsche Forschungsgemeinschaft (LE 1841/5-1). M.E.W. was supported by the Exascale Computing Project (No. 17-SC-20-SC), a collaborative effort of the U.S. Department of Energy Office of Science and the National Nuclear Security Administration, and the University of California Laboratory Fees Research Program (No. LFR-17-476732). Use of the Stanford Synchrotron Radiation Lightsource and Linac Coherent Light Source (LCLS), SLAC National Accelerator Laboratory, is supported by the U.S. Department of Energy, Office of Science, Office of Basic Energy Sciences under Contract No. DE-AC02-76SF00515. The SSRL Structural Molecular Biology Program is supported by the DOE Office of Biological and Environmental Research, and by the National Institutes of Health, National Institute of General Medical Sciences (P30GM133894). The contents of this publication are solely the responsibility of the authors and do not necessarily represent the official views of NIGMS or NIH. The Los Alamos National Laboratory technical release number of this article is LA-UR-22724.

## DATA AVAILABILITY

The data that support findings in this work are available on https://proteindiffraction.org, and the data accession codes are: 10.18430/m3.irrmc.5696 (WT-1),^45^ 10.18430/m3.irrmc.5695 (WT-2),^46^ 10.18430/m3.irrmc.5694 (WT-3),^47^ 10.18430/m3.irrmc.5693 (G150A-1),^48^ 10.18430/m3.irrmc.5692 (G150A-2),^49^ 10.18430/m3.irrmc.5691 (G150A-3),^50^ 10.18430/m3.irrmc.5690 (G150T-1),^51^ 10.18430/m3.irrmc.5689 (G150T-2),^52^ and 10.18430/m3.irrmc.5688 (G150T-3).^53^

