## Supplementary Material for "Reproducibility of protein X-ray diffuse scattering and potential utility for modeling atomic displacement parameters"

### I. Additional figures

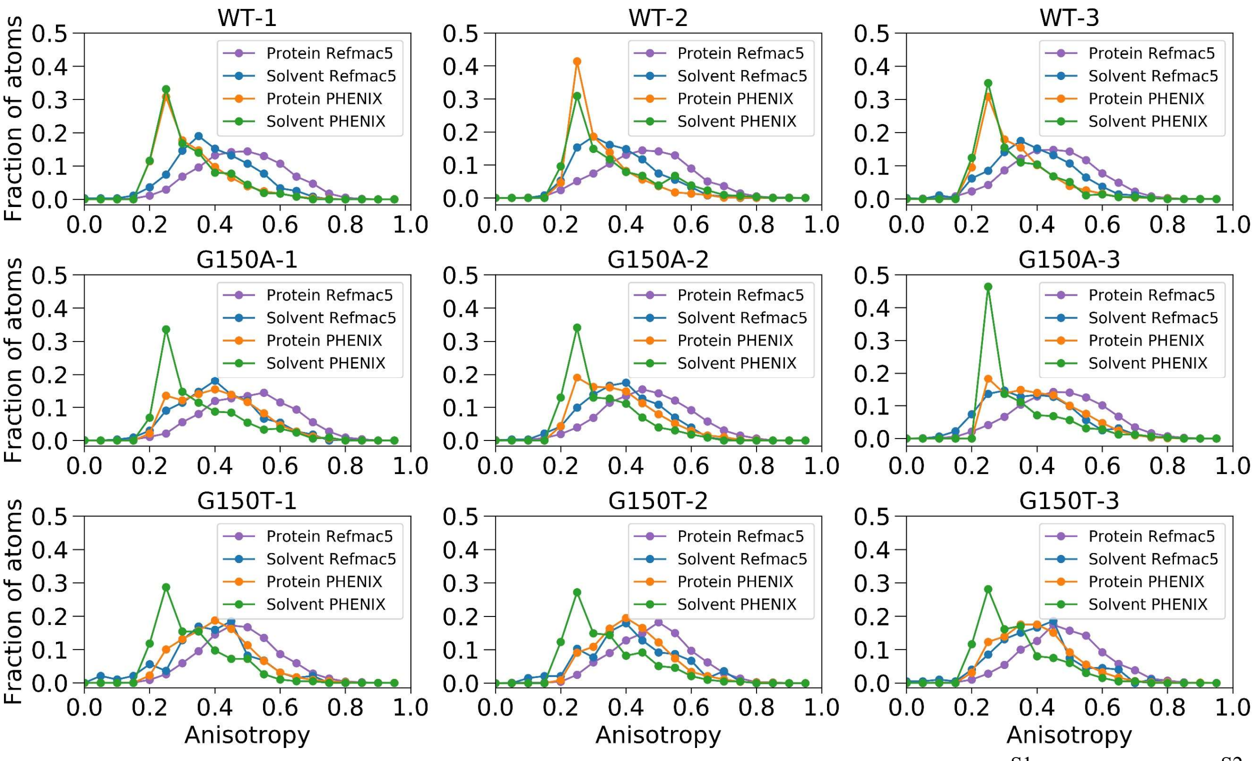


**FIG. S1.** The anisotropy distribution of Bragg models refined by PHENIX[S1] and Refmac5[S2]. Anisotropy is defined as the ratio of the smallest to the largest eigenvalues of the anisotropic ADP variance-covariance matrix.[S3]

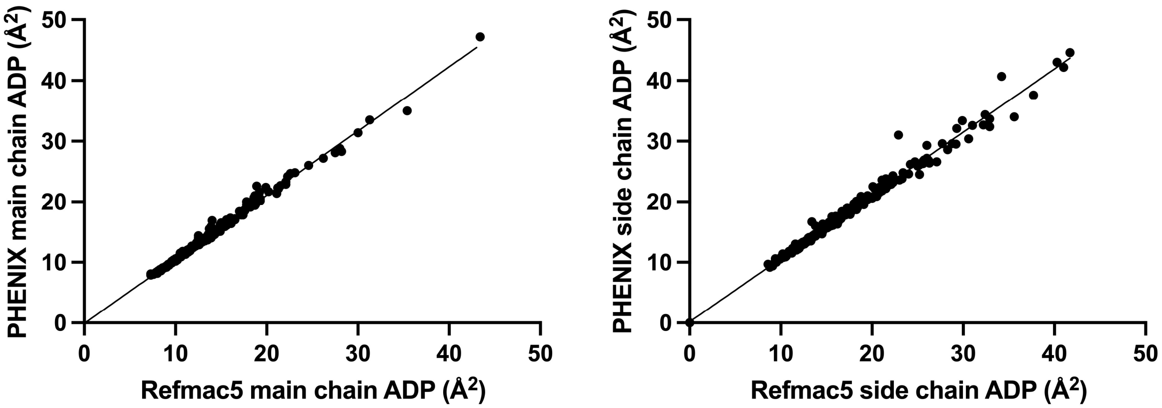


**FIG. S2.** ADP magnitudes are highly similar in the PHENIX- and Refmac5-refined models. Scatter plots of the ADP magnitudes for the main chain (left) and side chain (right) atoms of the WT-1 models are shown. ADP magnitudes in the other ICH models are comparably similar. In both cases, the ADP values refined by PHENIX and Refmac5 agree closely, with $R^2$ values of the best-fit line that exceed 0.99 and slopes near 1.

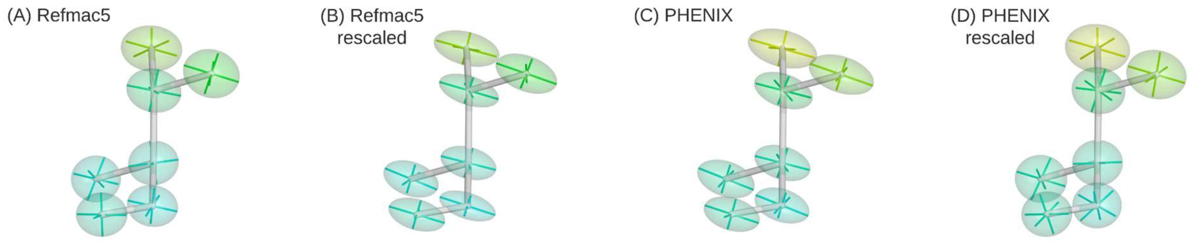


**FIG. S3.** Anisotropic rescaling can interconvert ADPs in the PHENIX- and Refmac5-refined models. Anisotropic ADP ellipsoids for Val100 in WT-1 ICH are represented at the 50% probability level and colored by $B_{eq}$, with warmer colors corresponding to larger $B_{eq}$ values. Panels (A) and (C) show the models as output by Refmac5 and PHENIX. Panel (B) shows the Refmac5-$U_{ztr}$ rescaled ADPs, which closely resemble the original PHENIX ADPs. Panel (D) shows the PHENIX+$U_{ztr}$ rescaled ADPs, which closely resemble the original Refmac5 ADPs. The figure was made with POVScript+[S4].

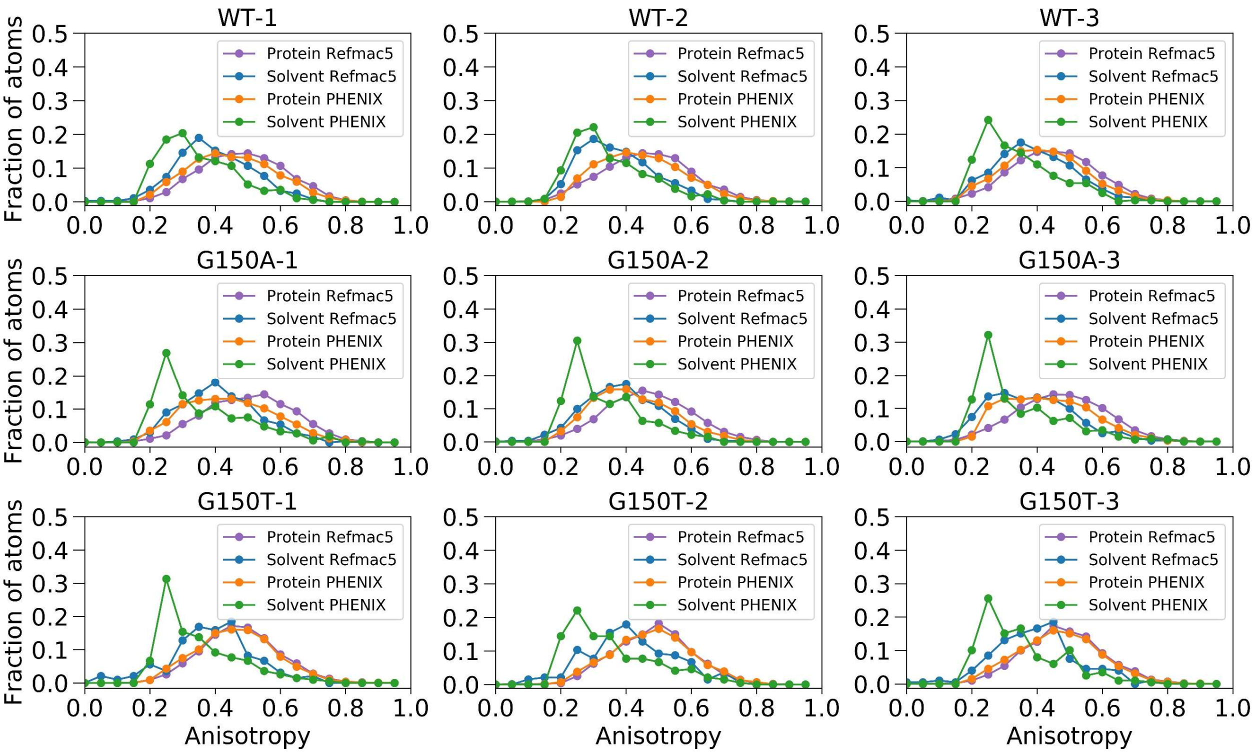


**FIG. S4.** The ADP anisotropy distribution for Refmac5 and PHENIX + $U_{ztr}$ rescaled models. After transforming the PHENIX model to a Refmac5-like model by addition of the difference anisotropic scaling matrix ($U_{ztr}$) to the ADPs, (see Methods), the anisotropy distributions (especially for protein atoms) agree closely.

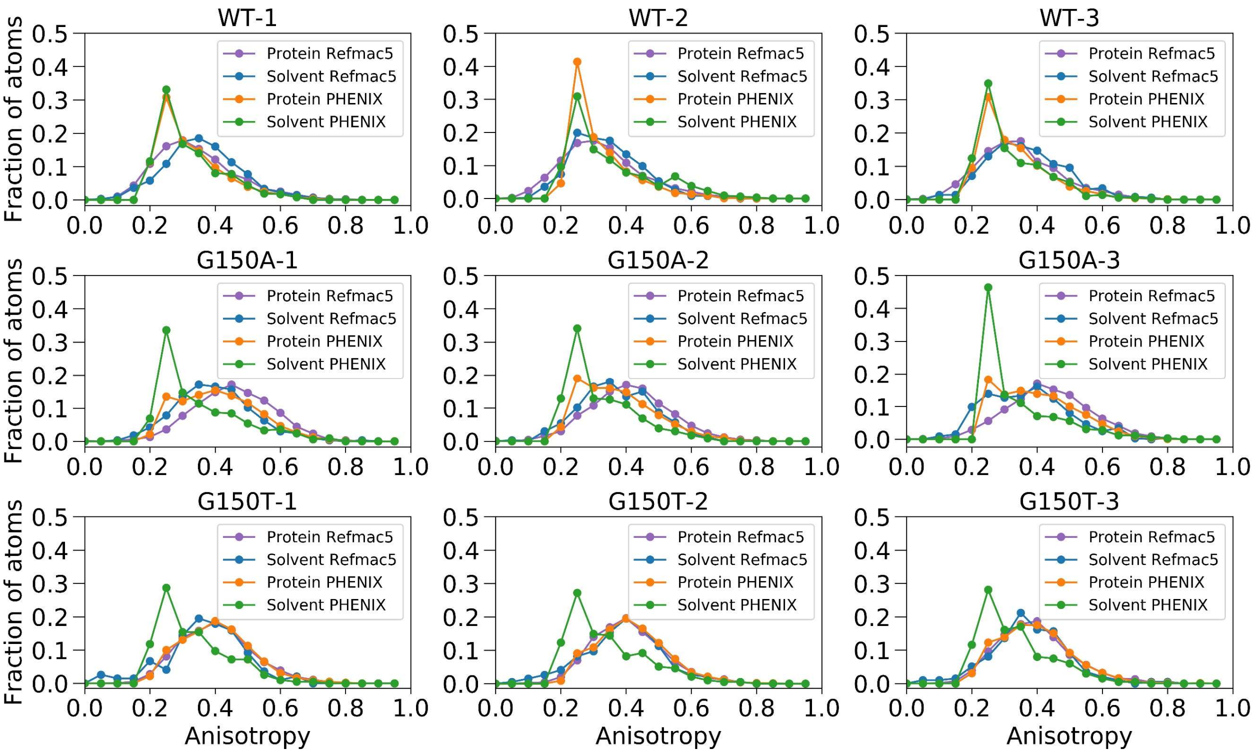


**FIG. S5.** The ADP anisotropy distribution for PHENIX and Refmac5 - $U_{ztr}$ rescaled models. After transforming the Refmac5 model to a PHENIX-like model by subtraction of the difference anisotropic scaling matrix ($U_{ztr}$) to the ADPs, (see Methods), the anisotropy distributions (especially for protein atoms) agree closely.

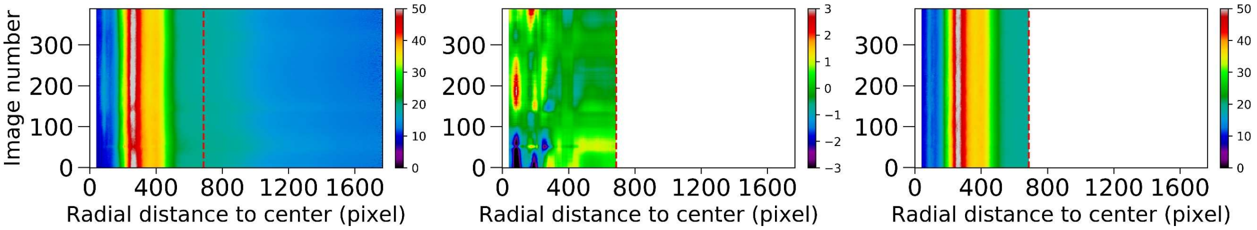


**FIG. S6.** Illustration of the radial profile variance removal method.[S5] The left side shows the scaled radial intensity profile stack of the whole WT-1 dataset before the radial profile variance removal method. Three largest PCA components of this radial profile stack are used to derive the radial profile variance as shown in the middle figure. This radial profile variance is subtracted as the isotropic background from each image, giving rise to variance removed diffraction patterns, whose new radial profile stack is shown in the right-side figure. The x axis is the distance of the radial shell to the detector center in pixels, and y axis is the image number. The red vertical line is at 1.4Å, above which data will not be used.

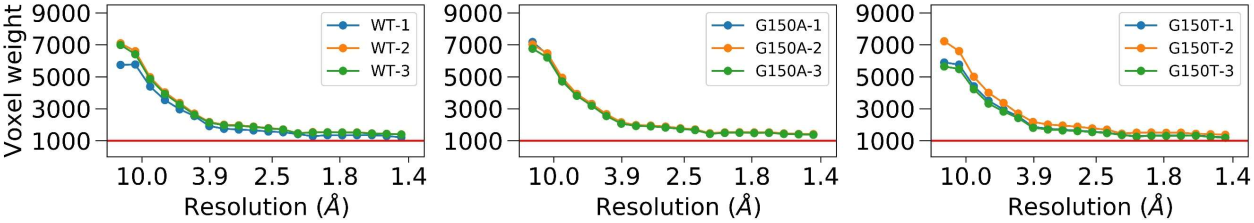


**FIG. S7.** Curves of the radially averaged number of pixels assigned to each non-empty voxel in the merged diffraction volume. Each curve is calculated using the diffraction volume which is one-time sampled over Miller indices. The x axis represents the resolution shell up to 1.4Å, and y axis represents the radially averaged number of pixels assigned per voxel in each resolution shell. The red line at 1000 shows that all non-empty voxels have at least 1000 contributing pixels on average in each resolution shell.

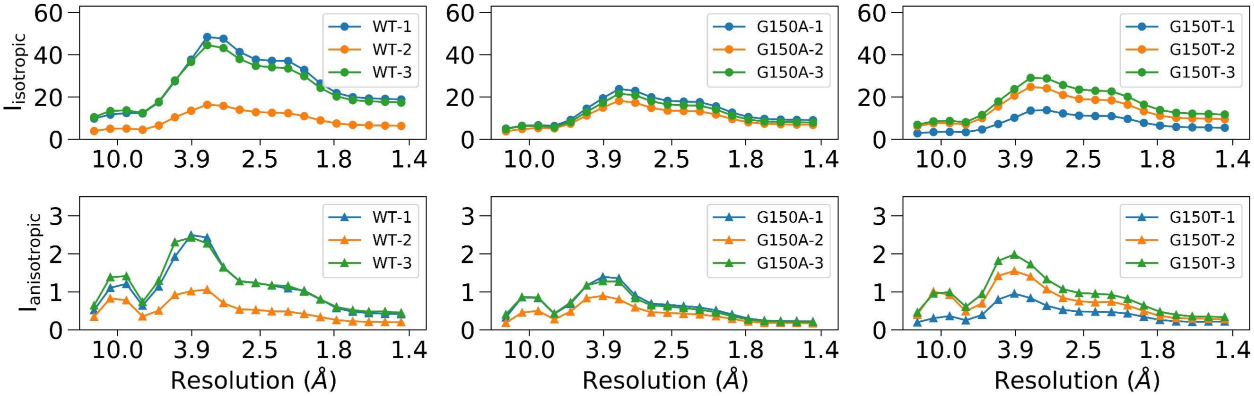


**FIG. S8.** Radially averaged voxel intensity profiles of the isotropic and anisotropic components in the diffraction volume. The first row shows the isotropic voxel intensity profile ($I_{isotropic}$) of each diffraction volume merged with the standard pipeline, and the second row shows the voxel intensity profile ($I_{anisotropic}$) of the anisotropic component. $I_{anisotropic}$ was calculated using absolute voxel intensities after $I_{isotropic}$ is subtracted from the diffraction volume. The x axis is the resolution shell up to 1.4Å, and y axis is the radially averaged voxel intensity in each resolution shell.

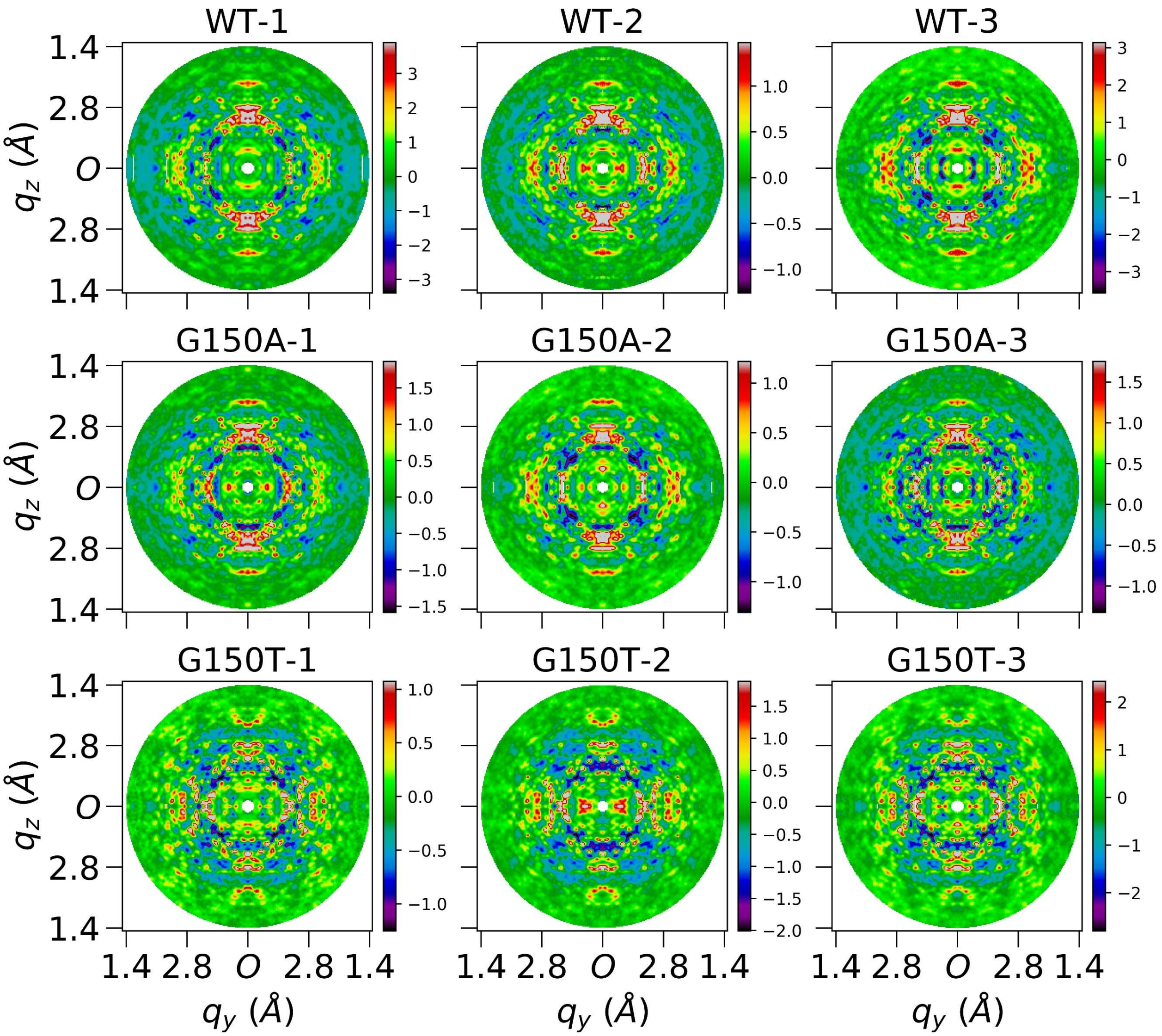


**FIG. S9.** Central slices of Laue-symmetrized anisotropic diffuse maps (standard pipeline) of nine datasets perpendicular to $q_x$ direction. Each image is cut from the center of the corresponding diffuse map which is three-time finely sampled over Miller indices *H,K,L*. Each subfigure shows average voxels within a depth of 0.05Å$^{-1}$ in $q_x$ direction, and 0.02 × 0.02Å$^{-1}$ in $q_yq_z$ plane. Both $q_y$ and $q_z$ axes extend to 1.4Å, and *O* represents the center in the reciprocal space. These finely sampled diffraction volumes were used for improved visualization only and were not used in data quality evaluation and modelling.

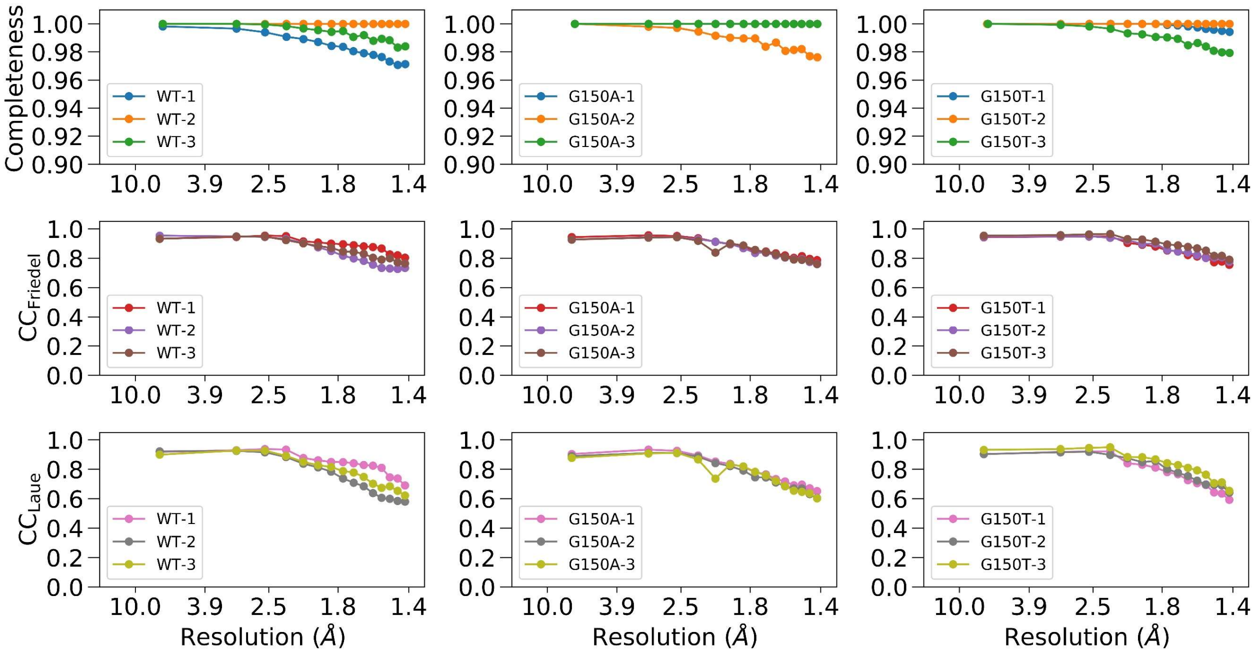


**FIG. S10.** The resolution-dependent completeness, CC$_{Friedel}$, and CC$_{Laue}$ curves of each dataset. The x axis is the resolution shell up to 1.4Å, and the y axis is the statistics in each resolution shell of the unsymmetrized anisotropic map of each dataset. The top, middle, and bottom rows show the completeness, CC$_{Friedel}$, and CC$_{Laue}$, respectively. The completeness was calculated using PHENIX[S1]. The completeness curve of G150A-1 overlaps with that of G150A-3.

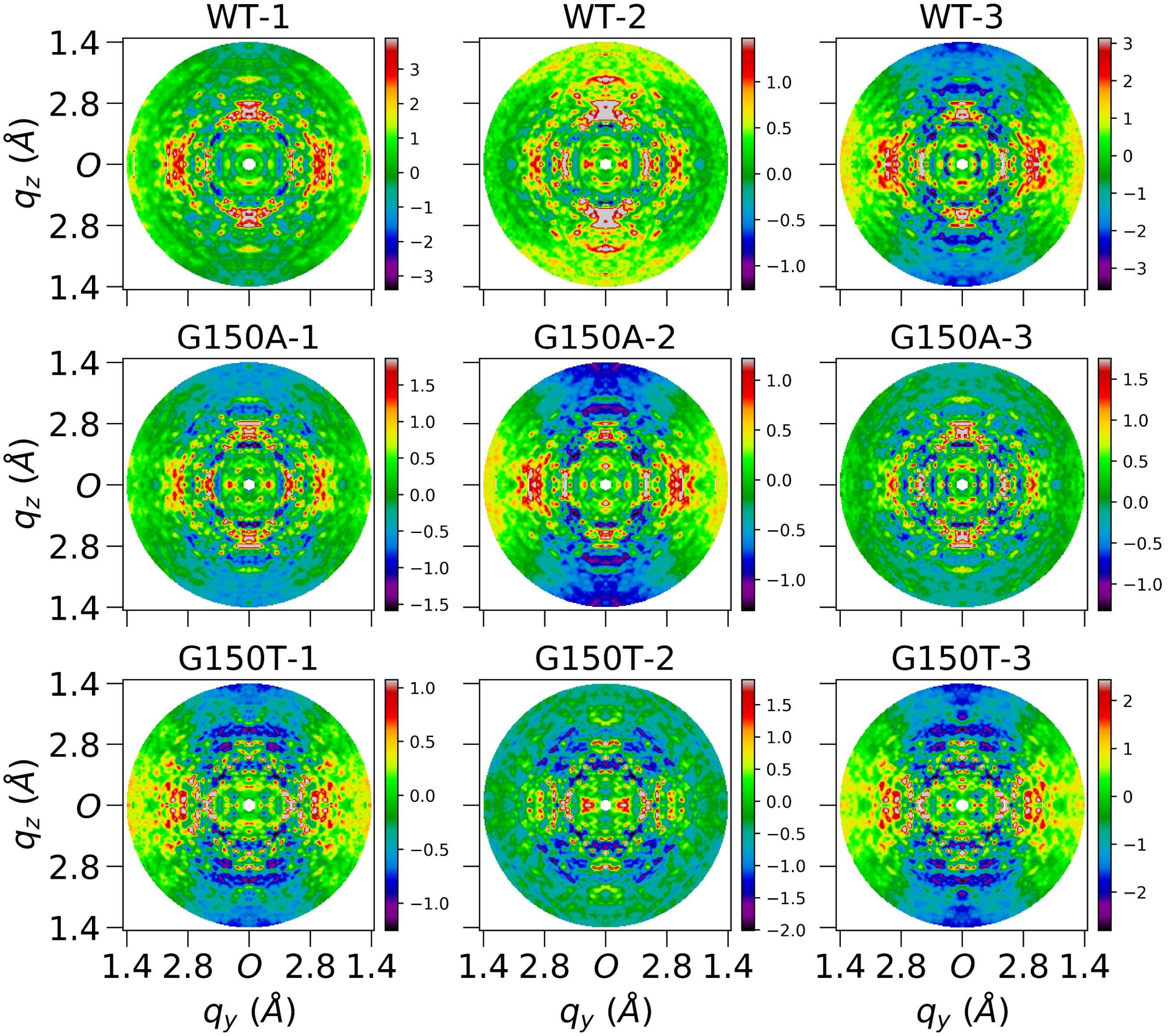


**FIG. S11.** Central slices of Laue-symmetrized anisotropic diffuse maps (without the polarization correction) of nine datasets perpendicular to $q_x$ direction. Each image is cut from the center of the corresponding diffuse map which is three-time finely sampled over Miller indices *H,K,L*. Each subfigure shows average voxels within a depth of 0.05Å$^{-1}$ in $q_x$ direction, and $0.02 \times 0.02$Å$^{-1}$ in $q_yq_z$ plane. Both $q_y$ and $q_z$ axes extend to 1.4Å, and *O* represents the center in the reciprocal space.

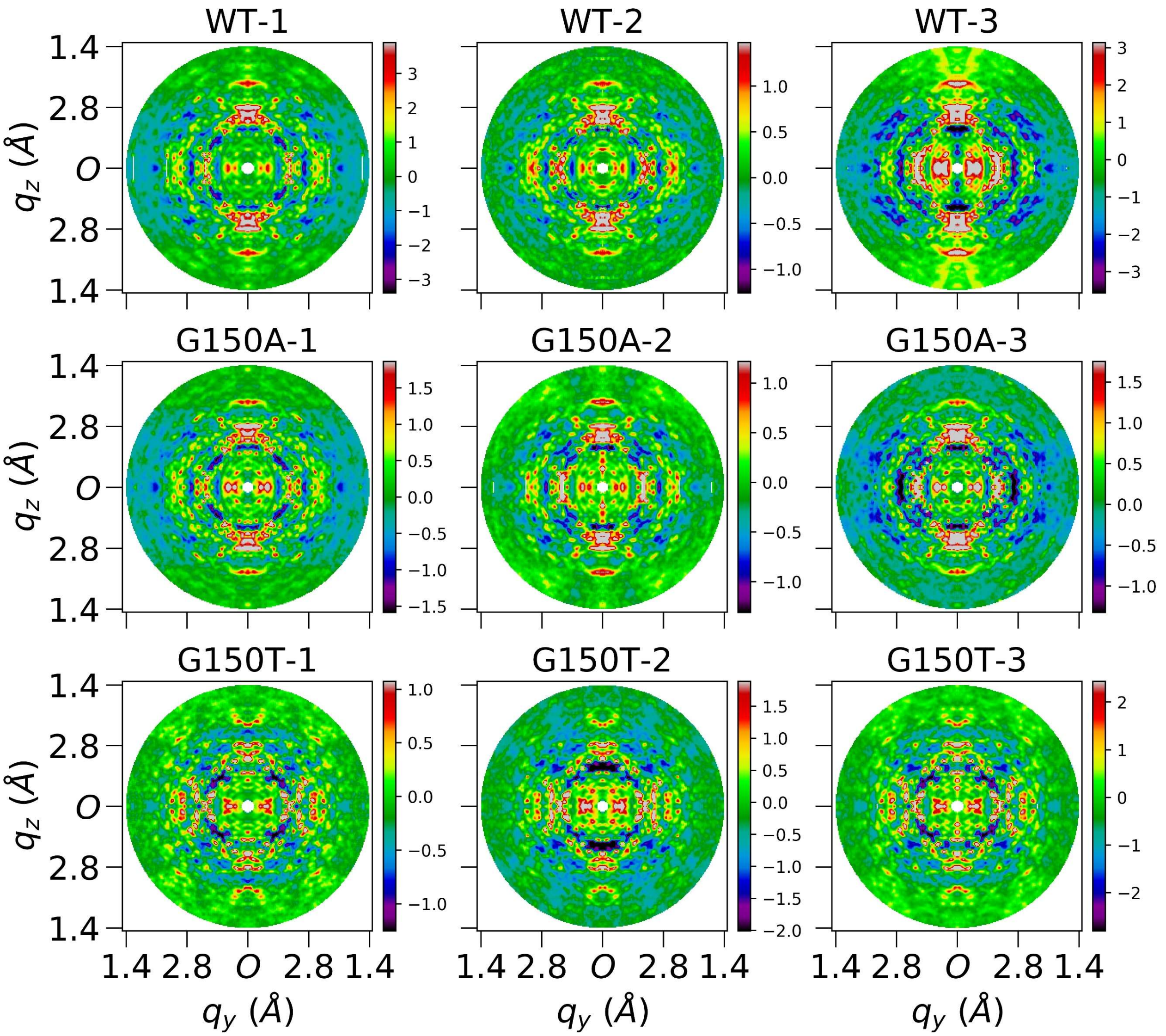


**FIG. S12.** Central slices of Laue-symmetrized anisotropic diffuse maps (without the radial profile variance removal step) of nine datasets perpendicular to $q_x$ direction. Each image is cut from the center of the corresponding diffuse map which is three-time finely sampled over Miller indices *H,K,L*. Each subfigure shows average voxels within a depth of 0.05Å$^{-1}$ in $q_x$ direction, and 0.02 × 0.02Å$^{-1}$ in $q_yq_z$ plane. Both $q_y$ and $q_z$ axes extend to 1.4Å, and *O* represents the center in the reciprocal space.

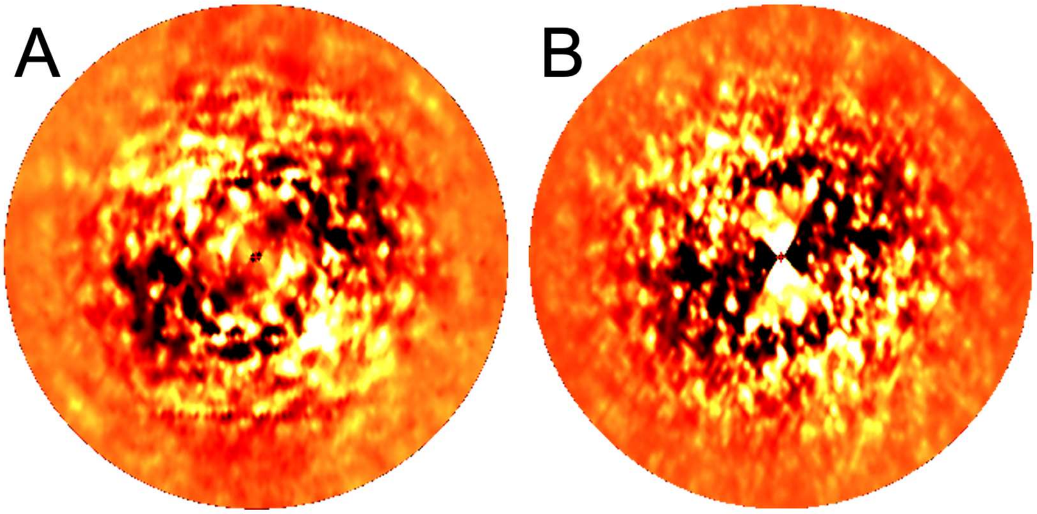


**FIG. S13.** Simulated diffraction patterns derived from the (A) experimental diffuse map and the (B) simulated diffuse map using the liquid-like motions (LLM) model. The orientation corresponds to the first image in the WT-1 dataset. The images are displayed using a heat map with progression black -> red -> yellow -> white as intensity increases from the lower to the upper value in the range (the range was chosen by eye to emphasize anisotropic diffuse features). The high resolution cut-off for both figures is 1.4Å.

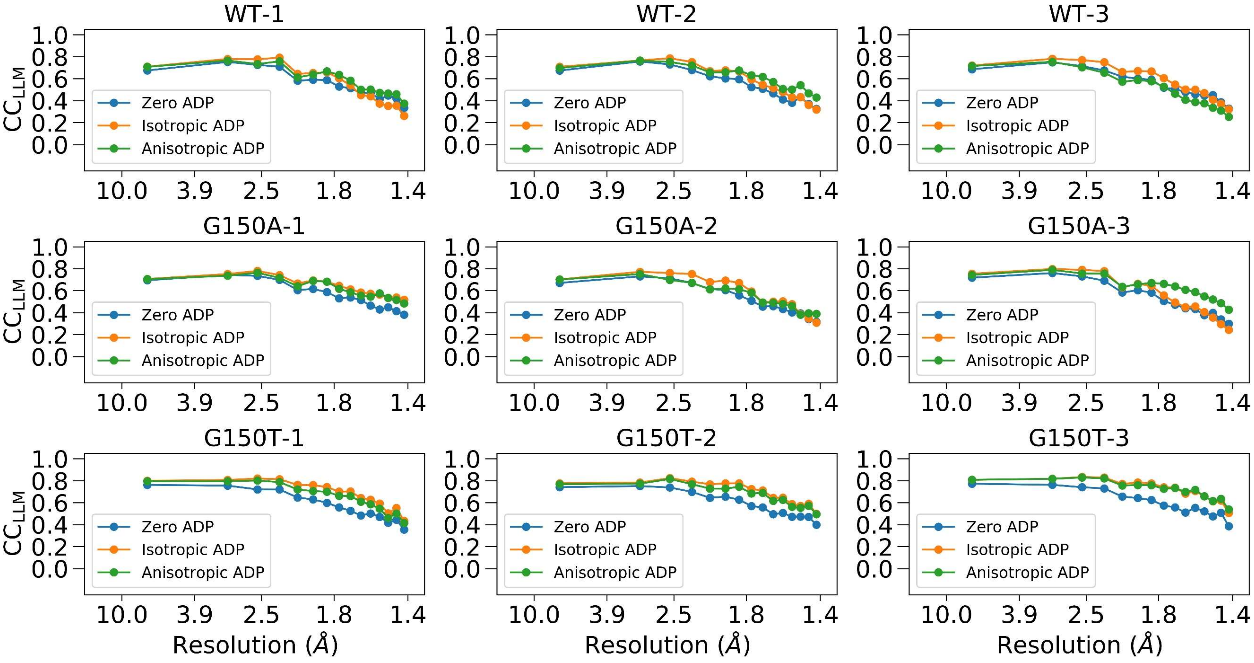


**FIG. S14.** The resolution dependent $CC_{LLM}$ curves of each dataset. The $CC_{LLM}$ was calculated as the CC between experimental and LLM simulated diffuse maps up to 1.4Å. The LLM model was performed using the Refmac5-refined protein structures with three types of ADP models: the zero, isotropic, and anisotropic ADP, respectively.

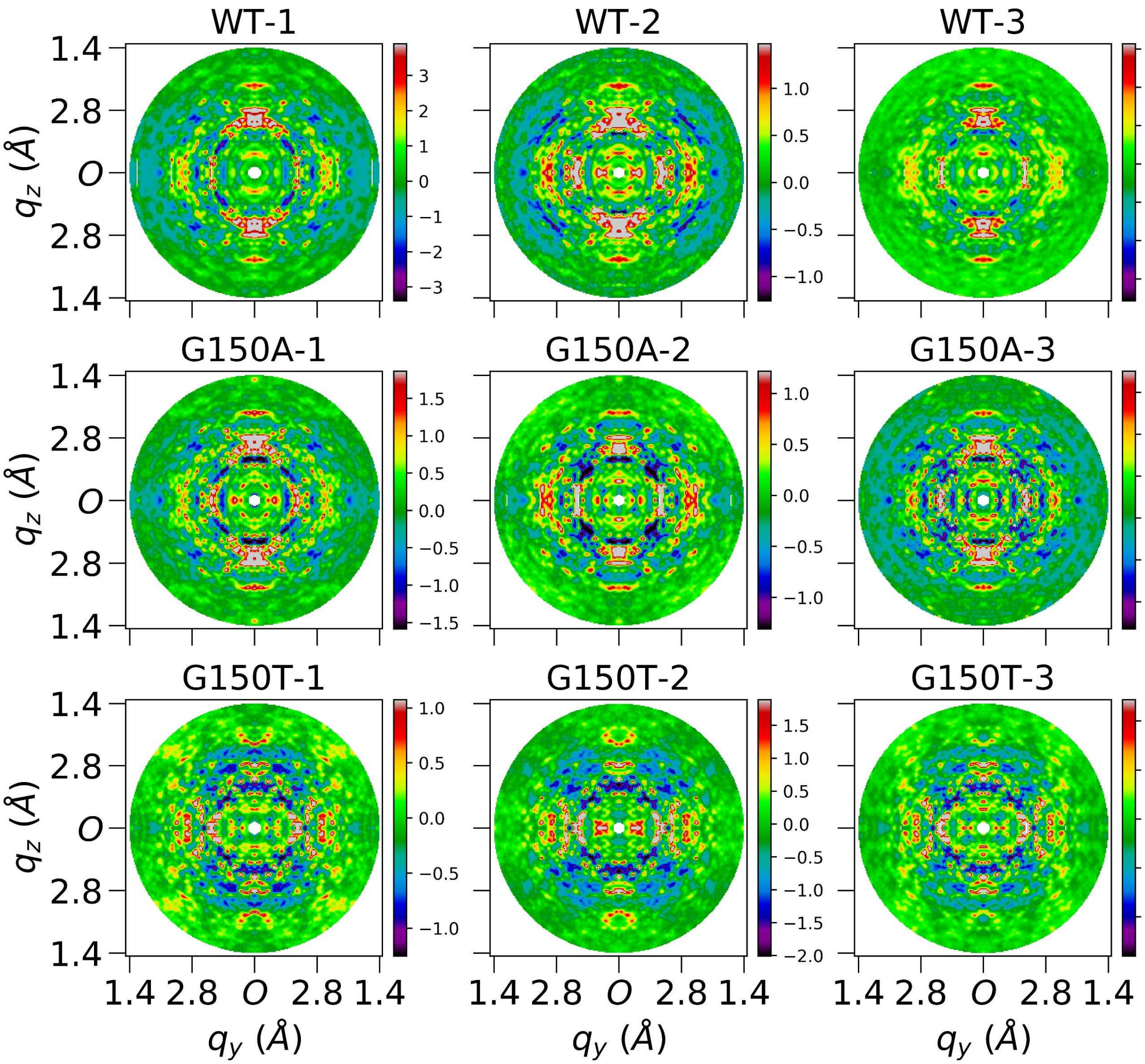


**FIG. S15.** Central slices of Laue-symmetrized anisotropic diffuse maps (without the non-crystal background image subtraction step) of nine datasets perpendicular to $q_x$ direction. Each image is cut from the center of the corresponding diffuse map which is three-time finely sampled over Miller indices *H,K,L*. Each subfigure shows average voxels within a depth of 0.05Å$^{-1}$ in $q_x$ direction, and 0.02 × 0.02Å$^{-1}$ in $q_yq_z$ plane. Both $q_y$ and $q_z$ axes extend to 1.4Å, and *O* represents the center in the reciprocal space.

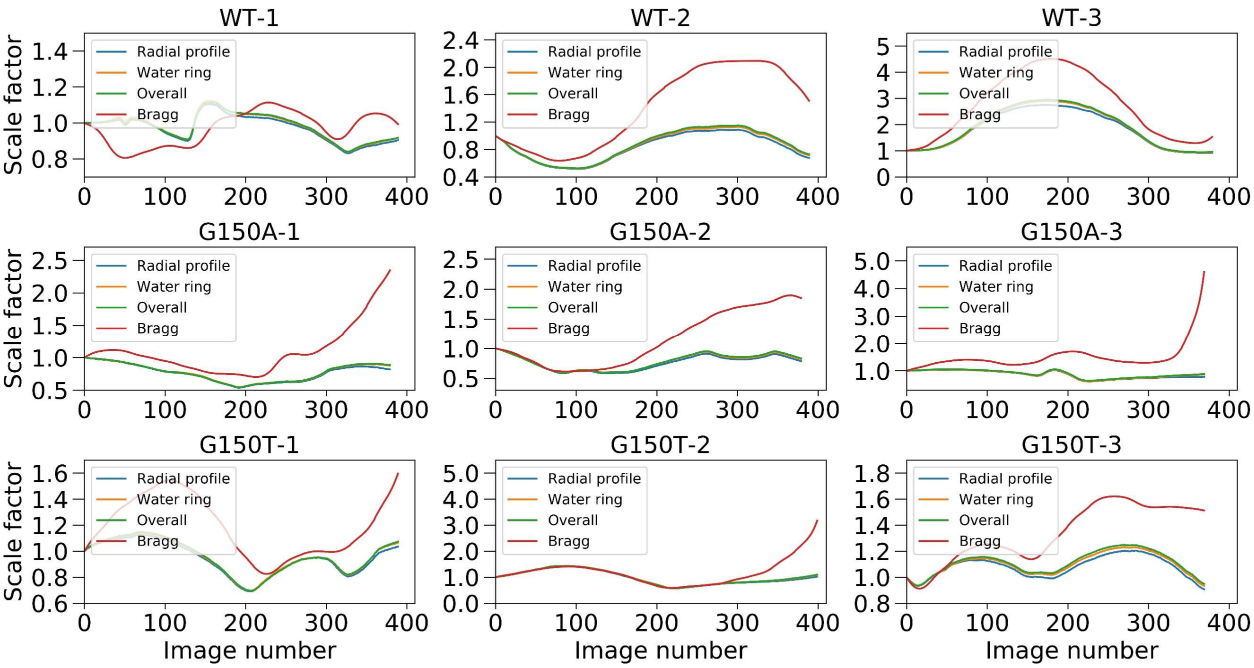


**FIG. S16.** Curves of four different scale factors of each dataset. These four scale factors are radial profile, water ring, overall, and Bragg intensity scale factor[S6], respectively. The x axis represents the image number in each dataset, and y axis is the per-image scale factor with the first image as the reference. The standard processing pipeline uses the radial profile scale factor.

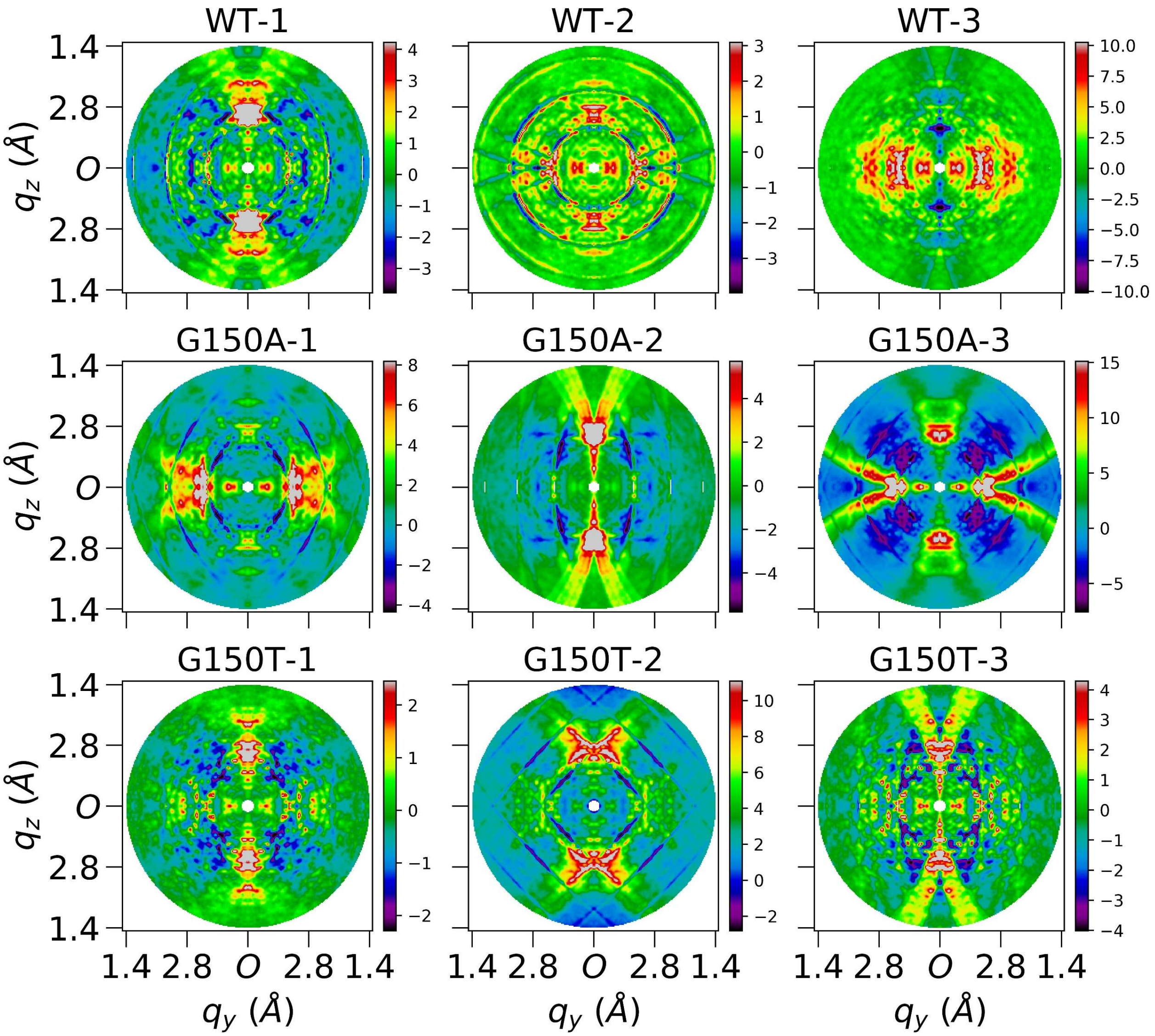


**FIG. S17.** Central slices of Laue-symmetrized anisotropic diffuse maps (with Bragg intensity scale factor and without the radial profile variance removal step) of nine datasets perpendicular to $q_x$ direction. Each image is cut from the center of the corresponding diffuse map which is three-time finely sampled over Miller indices *H,K,L*. Each subfigure shows average voxels within a depth of 0.05Å$^{-1}$ in $q_x$ direction, and 0.02 × 0.02Å$^{-1}$ in $q_yq_z$ plane. Both $q_y$ and $q_z$ axes extend to 1.4Å, and *O* represents the center in the reciprocal space.

### II. Additional tables

**TABLE S1.** Crystallographic data statistics

| Sample | WT-1 | WT-2 | WT-3 | G150A-1 | G150A-2 | G150A-3 | G150T-1 | G150T-2 | G150T-3 |
|---|---|---|---|---|---|---|---|---|---|
| Diffraction source | SSRL 12-2 | SSRL 12-2 | SSRL 12-2 | SSRL 12-2 | SSRL 12-2 | SSRL 12-2 | SSRL 12-2 | SSRL 12-2 | SSRL 12-2 |
| Wavelength (Å) | 0.775 | 0.775 | 0.775 | 0.775 | 0.775 | 0.775 | 0.775 | 0.775 | 0.775 |
| Temperature (K) | 274 | 274 | 274 | 274 | 274 | 274 | 274 | 274 | 274 |
| Detector | Pilatus 6M | Pilatus 6M | Pilatus 6M | Pilatus 6M | Pilatus 6M | Pilatus 6M | Pilatus 6M | Pilatus 6M | Pilatus 6M |
| Space group | $P2_1$ | $P2_1$ | $P2_1$ | $P2_1$ | $P2_1$ | $P2_1$ | I2 | I2 | I2 |
| a, b, c (Å) | 57.31 58.19 69.01 | 57.30 58.22 69.05 | 57.31 58.22 69.03 | 57.25 58.13 69.09 | 57.15 57.87 68.99 | 57.32 58.24 69.10 | 56.30 59.72 69.50 | 56.37 59.73 69.55 | 56.17 59.66 69.45 |
| α, β, γ (°) | 90.00 112.81 90.00 | 90.00 112.82 90.00 | 90.00 112.81 90.00 | 90.00 112.78 90.00 | 90.00 112.74 90.00 | 90.00 112.74 90.00 | 90.00 110.88 90.00 | 90.00 110.94 90.00 | 90.00 110.84 90.00 |
| Mosaicity (°) | 0.10 | 0.09 | 0.10 | 0.17 | 0.11 | 0.10 | 0.10 | 0.14 | 0.08 |
| Resolution range (Å) | 39.11-1.15 (1.17-1.15) | 38.67-1.20 (1.22-1.20) | 39.15-1.20 (1.22-1.20) | 39.11-1.30 (1.32-1.30) | 39.00-1.25 (1.27-1.25) | 39.17-1.35 (1.38-1.35) | 39.50-1.15 (1.17-1.15) | 35.22-1.20 (1.22-1.20) | 35.22-1.10 (1.12-1.10) |
| Total no. of observations | 544953 (24022) | 345556 (15550) | 469239 (20608) | 294077 (13740) | 328127 (14614) | 175502 (8066) | 280457 (12317) | 254082 (11584) | 302439 (13882) |
| No. of unique observations | 138486 (6553) | 127238 (6125) | 126107 (5967) | 100703 (4869) | 109380 (5115) | 85576 (4083) | 74206 (3528) | 66132 (3293) | 82868 (3983) |
| Completeness (%) | 93.5 (89.3) | 97.4 (94.3) | 96.5 (91.9) | 98.0 (95.8) | 95.6 (90.1) | 93.0 (90.0) | 97.4 (94.3) | 98.3 (95.8) | 95.7 (93.4) |
| Multiplicity | 3.9 (3.7) | 2.7 (2.5) | 3.7 (3.5) | 2.9 (2.8) | 3.0 (2.9) | 2.1 (2.0) | 3.8 (3.5) | 3.8 (3.5) | 3.6 (3.5) |
| $\langle I/\sigma(I) \rangle$ | 12.8 (1.4) | 6.8 (0.8) | 7.1 (1.1) | 10.4 (1.2) | 7.0 (0.8) | 6.0 (0.9) | 12.0 (0.9) | 7.3 (1.1) | 11.8 (1.0) |
| $CC_{1/2}$[1] | 0.999 (0.483) | 0.998 (0.257) | 0.997 (0.309) | 0.998 (0.405) | 0.998 (0.317) | 0.998 (0.264) | 0.999 (0.366) | 0.997 (0.334) | 0.998 (0.334) |
| $R_{meas}$[3] | 0.058 (1.280) | 0.068 (1.768) | 0.069 (1.737) | 0.057 (1.259) | 0.071 (1.851) | 0.064 (1.384) | 0.052 (1.720) | 0.078 (2.243) | 0.057 (1.839) |

[1]$CC_{1/2}$ [S7] was used to determine the high resolution cutoff.

**TABLE S2.** Refmac5 crystallographic refinement statistics

| Sample | WT-1 | WT-2 | WT-3 | G150A-1 | G150A-2 | G150A-3 | G150T-1 | G150T-2 | G150T-3 |
|---|---|---|---|---|---|---|---|---|---|
| PDB code | 7L9Q | 7L9S | 7L9W | 7L9Z | 7LA0 | 7LA3 | 7LAV | 7LAX | 7LB9 |
| Temperature (K) | 274 | 274 | 274 | 274 | 274 | 274 | 274 | 274 | 274 |
| Refinement program | Refmac 5.8.0266 | Refmac 5.8.0266 | Refmac 5.8.0266 | Refmac 5.8.0267 | Refmac 5.8.0267 | Refmac 5.8.0267 | Refmac 5.8.0266 | Refmac 5.8.0267 | Refmac 5.8.0267 |
| Resolution range (Å) | 39.11-1.15 (1.18-1.15) | 38.67-1.20 (1.23-1.20) | 39.15-1.20 (1.23-1.20) | 39.11-1.30 (1.33-1.30) | 39.00- 1.25 (1.28-1.25) | 39.17-1.35 (1.38-1.35) | 39.50-1.15 (1.18-1.15) | 35.22-1.20 (1.23-1.20) | 35.22-1.10 (1.13-1.10) |
| Completeness (%) | 93.25 (89.59) | 97.25 (94.96) | 96.35 (93.97) | 97.88 (96.77) | 94.28 (83.40) | 91.95 (85.63) | 97.15 (94.42) | 98.03 (93.84) | 95.29 (91.89) |
| No. of reflections | 138457 (9500) | 127216 (8890) | 126085 (8791) | 100597 (7138) | 108127 (6800) | 84895 (5658) | 74206 (5135) | 66132 (4501) | 82864 (5756) |
| No. of reflections, test set | 4137 (281) | 3838 (271) | 3768 (268) | 3076 (206) | 3309 (237) | 2602 (176) | 2238 (161) | 1994 (132) | 2457 (151) |
| $R_{work}$ | 0.1209 (0.2490) | 0.1234 (0.2920) | 0.1244 (0.2670) | 0.1191 (0.2980) | 0.1237 (0.3310) | 0.1201 (0.2970) | 0.1272 (0.2830) | 0.1241 (0.2710) | 0.1267 (0.2810) |
| $R_{free}$ | 0.1440 (0.2480) | 0.1523 (0.3070) | 0.1510 (0.2880) | 0.1463 (0.3100) | 0.1524 (0.3180) | 0.1554 (0.3340) | 0.1425 (0.3070) | 0.1447 (0.2570) | 0.1407 (0.2830) |
| No. of non-H atoms | | | | | | | | | |
| Protein | 3997 | 3965 | 3983 | 4363 | 4384 | 4380 | 1908 | 1884 | 1884 |
| Water | 363 | 366 | 355 | 334 | 334 | 323 | 195 | 195 | 199 |
| Total | 4360 | 4331 | 4338 | 4697 | 4718 | 4703 | 2103 | 2079 | 2083 |
| Average R.M.S. deviations | | | | | | | | | |
| Bonds (Å) | 0.008 | 0.009 | 0.009 | 0.007 | 0.008 | 0.008 | 0.009 | 0.009 | 0.008 |
| Angles (°) | 1.493 | 1.520 | 1.529 | 1.434 | 1.473 | 1.466 | 1.510 | 1.492 | 1.466 |
| Average B factors ($<B_{iso}>$;Å$^2$) | | | | | | | | | |
| Protein | 17.01 | 16.96 | 17.64 | 19.72 | 18.42 | 18.41 | 17.29 | 17.45 | 15.97 |
| Water | 33.95 | 34.08 | 34.78 | 36.38 | 35.63 | 33.96 | 36.13 | 35.67 | 34.59 |
| Average ADP anisotropy[1] | | | | | | | | | |
| Protein | 0.508 | 0.488 | 0.473 | 0.521 | 0.465 | 0.492 | 0.501 | 0.519 | 0.506 |
| Water | 0.420 | 0.386 | 0.415 | 0.417 | 0.395 | 0.404 | 0.415 | 0.439 | 0.420 |
| MolProbity clashscore | 1.5 | 1.5 | 1.2 | 3.5 | 3.2 | 2.9 | 2.1 | 1.8 | 1.6 |
| Ramachandran plot | | | | | | | | | |
| Outliers (%) | 0.44 | 0.44 | 0.44 | 0.44 | 0.44 | 0.88 | 0.00 | 0.00 | 0.00 |
| Allowed (%) | 1.33 | 1.33 | 1.33 | 1.99 | 1.99 | 1.33 | 0.89 | 0.89 | 0.45 |
| Favored (%) | 98.23 | 98.23 | 98.23 | 97.57 | 97.57 | 97.79 | 99.11 | 99.11 | 99.55 |

[1]Anisotropy is defined as the ratio of the smallest to largest eigenvalue of the ADP tensor and was calculated using PARVATI[S3]

**TABLE S3.** PHENIX crystallographic refinement statistics

| Sample | WT-1 | WT-2 | WT-3 | G150A-1 | G150A-2 | G150A-3 | G150T-1 | G150T-2 | G150T-3 |
|---|---|---|---|---|---|---|---|---|---|
| PDB code | 7LBH | 7LBI | 7LCX | 7LD6 | 7LD7 | 7LDB | 7LDM | 7LDI | 7LDO |
| Temperature (K) | 274 | 274 | 274 | 274 | 274 | 274 | 274 | 274 | 274 |
| Refinement program | PHENIX 1.17.1 | PHENIX 1.17.1 | PHENIX 1.17.1 | PHENIX 1.17.1 | PHENIX 1.17.1 | PHENIX 1.17.1 | PHENIX 1.17.1 | PHENIX 1.17.1 | PHENIX 1.17.1 |
| Resolution range (Å) | 31.81-1.15 (1.16-1.15) | 34.58-1.20 (1.22-1.20) | 38.64-1.20 (1.22-1.20) | 34.59-1.30 (1.32-1.30) | 38.49- 1.25 (1.27-1.25) | 34.63-1.35 (1.37-1.35) | 35.20-1.15 (1.17-1.15) | 35.20-1.20 (1.23-1.20) | 35.20-1.10 (1.12-1.10) |
| Completeness (%) | 93.28 (88.00) | 96.69 (91.00) | 96.42 (94.00) | 97.82 (94.00) | 94.30 (79.00) | 91.98 (84.00) | 97.16 (94.00) | 97.99 (93.00) | 95.37 (92.00) |
| No. of reflections | 138442 (4201) | 126307 (4274) | 125975 (4424) | 100616 (4274) | 108107 (3619) | 84886 (3898) | 74185 (4358) | 66081 (4324) | 82831 (4318) |
| No. of reflections, test set | 4137 (124) | 3808 (123) | 3767 (129) | 3075 (135) | 3308 (132) | 2602 (120) | 2237 (133) | 1993 (122) | 2457 (123) |
| $R_{work}$ | 0.1168 (0.2571) | 0.1207 (0.2723) | 0.1211 (0.2629) | 0.1198 (0.2848) | 0.1182 (0.2550) | 0.1193 (0.2539) | 0.1221 (0.2941) | 0.1190 (0.2590) | 0.1181 (0.2668) |
| $R_{free}$ | 0.1431 (0.2744) | 0.1526 (0.2992) | 0.1508 (0.3025) | 0.1434 (0.2959) | 0.1433 (0.2847) | 0.1535 (0.2690) | 0.1387 (0.3108) | 0.1373 (0.2571) | 0.1363 (0.2961) |
| No. of non-H atoms | | | | | | | | | |
| Protein | 3997 | 3965 | 3983 | 4363 | 4384 | 4380 | 1908 | 1884 | 1884 |
| Water | 363 | 366 | 355 | 334 | 334 | 323 | 195 | 195 | 199 |
| Total | 4360 | 4331 | 4338 | 4697 | 4718 | 4703 | 2103 | 2079 | 2083 |
| Average R.M.S. deviations | | | | | | | | | |
| Bonds (Å) | 0.008 | 0.007 | 0.007 | 0.004 | 0.004 | 0.008 | 0.005 | 0.007 | 0.009 |
| Angles (°) | 0.949 | 0.898 | 0.922 | 0.799 | 0.815 | 0.986 | 0.805 | 0.959 | 1.018 |
| Average B factors ($<B_{iso}>$;$Å^2$) | | | | | | | | | |
| Protein | 18.51 | 19.05 | 18.82 | 20.26 | 19.10 | 19.55 | 18.41 | 18.62 | 17.05 |
| Water | 35.72 | 36.69 | 36.57 | 39.11 | 37.91 | 37.43 | 38.53 | 38.42 | 36.97 |
| Average ADP anisotropy[1] | | | | | | | | | |
| Protein | 0.348 | 0.354 | 0.351 | 0.434 | 0.398 | 0.422 | 0.429 | 0.439 | 0.416 |
| Water | 0.345 | 0.343 | 0.341 | 0.369 | 0.345 | 0.359 | 0.355 | 0.364 | 0.355 |
| MolProbity clashscore | 1.1 | 1.3 | 0.9 | 1.8 | 1.8 | 2.1 | 1.5 | 1.3 | 1.6 |
| Ramachandran plot | | | | | | | | | |
| Outliers (%) | 0.44 | 0.44 | 0.44 | 0.44 | 0.44 | 0.88 | 0.00 | 0.00 | 0.00 |
| Allowed (%) | 1.33 | 1.33 | 1.33 | 1.99 | 1.99 | 1.33 | 0.89 | 0.89 | 0.45 |
| Favored (%) | 98.23 | 98.23 | 98.23 | 97.57 | 97.57 | 97.79 | 99.11 | 99.11 | 99.55 |

[1]Anisotropy is defined as the ratio of the smallest to largest eigenvalue of the ADP tensor and was calculated using PARVATI[S3]

**TABLE S4.** The anisotropic scaling matrices for Refmac5- and PHENIX-refined PDB files and the traceless $U_{ztr}$ matrix for three example datasets. The anisotropic scaling matrices (in the unit cell basis) are read from the header in PDB files after zero cycles of refinement (i.e., only scaling) in PDB-REDO[S8]. $U_{ztr}$ is their difference matrix in the PDB Cartesian basis after trace removal and constant $2\pi^2$ scaling.

| Sample | Refmac5 | PHENIX | $U_{ztr}$[1] |
|---|---|---|---|
| WT-1 | | | |
| | $\begin{bmatrix} 0.024 & 0 & 0.107 \\ 0 & -0.040 & 0 \\ 0.107 & 0 & -0.055 \end{bmatrix}$ | $\begin{bmatrix} 0.06 & 0 & -0.27 \\ 0 & -0.60 & 0 \\ -0.27 & 0 & 0.56 \end{bmatrix}$ | $\begin{bmatrix} 0.016 & 0 & -0.034 \\ 0 & -0.038 & 0 \\ -0.034 & 0 & 0.022 \end{bmatrix}$ |
| G150A-1 | | | |
| | $\begin{bmatrix} 0.025 & 0 & 0.231 \\ 0 & -0.265 & 0 \\ 0.231 & 0 & 0.034 \end{bmatrix}$ | $\begin{bmatrix} -0.04 & 0 & -0.22 \\ 0 & -0.21 & 0 \\ -0.22 & 0 & 0.32 \end{bmatrix}$ | $\begin{bmatrix} 0.007 & 0 & -0.031 \\ 0 & -0.009 & 0 \\ -0.031 & 0 & 0.002 \end{bmatrix}$ |
| G150T-1 | | | |
| | $\begin{bmatrix} -0.127 & 0 & -0.129 \\ 0 & 0.029 & 0 \\ -0.129 & 0 & 0.152 \end{bmatrix}$ | $\begin{bmatrix} -0.59 & 0 & -0.35 \\ 0 & 0.17 & 0 \\ -0.35 & 0 & 0.53 \end{bmatrix}$ | $\begin{bmatrix} -0.019 & 0 & -0.019 \\ 0 & 0.003 & 0 \\ -0.019 & 0 & 0.015 \end{bmatrix}$ |

[1]$U_{ztr}$ is the zero-trace U matrix that relates Refmac5- and PHENIX-refined PDBs.

**TABLE S5.** The data quality metrics of the anisotropic diffuse map of each dataset. This table contains the $CC_{1/2}$ (blue) of each dataset, the intra-protein correlations (green) between every two independent datasets of the same protein, and the inter-protein cross correlations (orange) between datasets of WT and G150A. All CC values were calculated using all data up to 1.4Å. The $CC_{Rep}$ for each dataset reported in the paper is the average of two intra-protein CC values in this matrix, and the final $CC_{Cross}$ is the average of three inter-protein CC in each row.

| Sample | WT-1 | WT-2 | WT-3 | G150A-1 | G150A-2 | G150A-3 | G150T-1 | G150T-2 | G150T-3 |
|---|---|---|---|---|---|---|---|---|---|
| WT-1 | 0.85 | 0.85 | 0.87 | 0.82 | 0.83 | 0.85 | | | |
| WT-2 | | 0.78 | 0.83 | 0.87 | 0.80 | 0.85 | | | |
| WT-3 | | | 0.81 | 0.83 | 0.88 | 0.85 | | | |
| G150A-1 | | | | 0.81 | 0.80 | 0.83 | | | |
| G150A-2 | | | | | 0.76 | 0.81 | | | |
| G150A-3 | | | | | | 0.77 | | | |
| G150T-1 | | | | | | | 0.82 | 0.85 | 0.91 |
| G150T-2 | | | | | | | | 0.84 | 0.88 |
| G150T-3 | | | | | | | | | 0.89 |

**TABLE S6.** The CC statistics of each dataset analyzed with seven different data processing choices. The diffuse map generated by each processing method was evaluated with five CC metrics: $CC_{Friedel}$, $CC_{Laue}$, $CC_{1/2}$, $CC_{Rep}$, and $CC_{LLM}$ (anisotropic ADP model). Method A (standard processing pipeline) contains CC values of each dataset up to 1.4Å resolution, while other methods (B)-(G) show the relative CC changes compared to those in method A. Cells in (B)-(G) are colored with four different colors depending on the relative CC changes. A cell is colored as white if the relative CC change is 0.00 or ±0.00, as light blue/red if CC increases/decreases by less than 0.1, otherwise it will be colored as dark blue/red.

| Sample | WT-1 | WT-2 | WT-3 | G150A-1 | G150A-2 | G150A-3 | G150T-1 | G150T-2 | G150T-3 |
|---|---|---|---|---|---|---|---|---|---|
| A. Standard data processing pipeline | | | | | | | | | |
| $CC_{Friedel}$ | 0.93 | 0.91 | 0.91 | 0.93 | 0.91 | 0.91 | 0.91 | 0.92 | 0.94 |
| $CC_{Laue}$ | 0.90 | 0.87 | 0.87 | 0.88 | 0.86 | 0.85 | 0.86 | 0.88 | 0.91 |
| $CC_{1/2}$ | 0.85 | 0.78 | 0.81 | 0.81 | 0.76 | 0.77 | 0.82 | 0.84 | 0.89 |
| $CC_{Rep}$ | 0.86 | 0.84 | 0.85 | 0.82 | 0.81 | 0.82 | 0.88 | 0.87 | 0.89 |
| $CC_{LLM}$ | 0.70 | 0.71 | 0.67 | 0.70 | 0.68 | 0.73 | 0.76 | 0.75 | 0.80 |
| B. Standard pipeline without non-crystal background image subtraction | | | | | | | | | |
| $CC_{Friedel}$ | -0.01 | -0.01 | -0.02 | -0.01 | -0.03 | -0.02 | -0.03 | -0.02 | -0.02 |
| $CC_{Laue}$ | -0.01 | -0.03 | -0.02 | -0.02 | -0.03 | -0.03 | -0.03 | -0.04 | -0.02 |
| $CC_{1/2}$ | -0.02 | -0.06 | -0.04 | -0.04 | -0.07 | -0.06 | -0.06 | -0.06 | -0.03 |
| $CC_{Rep}$ | -0.03 | -0.05 | -0.05 | -0.08 | -0.08 | -0.05 | -0.03 | -0.05 | -0.03 |
| $CC_{LLM}$ | -0.01 | -0.02 | -0.04 | -0.03 | -0.06 | -0.02 | -0.04 | -0.03 | -0.02 |
| C. Standard pipeline without the polarization correction | | | | | | | | | |
| $CC_{Friedel}$ | +0.04 | +0.04 | +0.04 | +0.03 | +0.05 | +0.04 | +0.03 | +0.03 | +0.02 |
| $CC_{Laue}$ | +0.05 | -0.02 | +0.04 | -0.01 | +0.07 | -0.03 | +0.03 | -0.11 | +0.03 |
| $CC_{1/2}$ | +0.08 | -0.06 | +0.08 | -0.06 | +0.13 | -0.11 | +0.02 | -0.19 | +0.04 |
| $CC_{Rep}$ | -0.13 | -0.32 | -0.27 | -0.58 | -0.31 | -0.32 | -0.16 | -0.35 | -0.15 |
| $CC_{LLM}$ | -0.17 | -0.14 | -0.27 | -0.23 | -0.33 | -0.16 | -0.18 | -0.09 | -0.19 |
| D. Standard pipeline without the radial profile variance removal step | | | | | | | | | |
| $CC_{Friedel}$ | -0.01 | -0.01 | -0.02 | -0.02 | -0.01 | -0.03 | -0.02 | -0.03 | -0.00 |
| $CC_{Laue}$ | -0.02 | -0.06 | -0.01 | -0.05 | -0.10 | -0.07 | -0.03 | -0.04 | -0.03 |
| $CC_{1/2}$ | -0.04 | -0.11 | -0.03 | -0.09 | -0.18 | -0.15 | -0.05 | -0.07 | -0.04 |
| $CC_{Rep}$ | -0.05 | -0.08 | -0.11 | -0.04 | -0.02 | -0.04 | -0.01 | -0.02 | -0.01 |
| $CC_{LLM}$ | +0.01 | -0.01 | +0.00 | -0.01 | -0.01 | -0.06 | -0.01 | -0.04 | -0.01 |
| E. Standard pipeline without the solid-angle correction | | | | | | | | | |
| $CC_{Friedel}$ | +0.00 | +0.01 | +0.01 | +0.01 | +0.01 | +0.01 | +0.01 | +0.01 | +0.01 |
| $CC_{Laue}$ | +0.01 | +0.02 | +0.01 | +0.01 | +0.01 | +0.01 | +0.01 | +0.01 | +0.01 |
| $CC_{1/2}$ | +0.01 | +0.03 | +0.02 | +0.02 | +0.02 | +0.02 | +0.03 | +0.02 | +0.01 |
| $CC_{Rep}$ | +0.01 | +0.01 | +0.01 | +0.01 | +0.01 | +0.01 | +0.01 | +0.01 | +0.01 |
| $CC_{LLM}$ | +0.01 | +0.00 | +0.01 | +0.01 | +0.01 | +0.01 | +0.01 | +0.00 | +0.00 |
| F. Standard pipeline without the detector absorption correction | | | | | | | | | |
| $CC_{Friedel}$ | -0.00 | -0.00 | -0.00 | -0.00 | -0.00 | -0.00 | -0.00 | -0.00 | -0.00 |
| $CC_{Laue}$ | -0.00 | -0.00 | -0.00 | -0.00 | -0.00 | -0.00 | -0.00 | -0.00 | -0.00 |
| $CC_{1/2}$ | -0.00 | -0.01 | -0.01 | -0.01 | -0.01 | -0.01 | -0.01 | -0.00 | -0.00 |
| $CC_{Rep}$ | -0.00 | -0.00 | -0.00 | -0.00 | -0.00 | -0.00 | -0.00 | -0.00 | -0.00 |
| $CC_{LLM}$ | -0.00 | -0.00 | -0.00 | -0.00 | -0.00 | -0.00 | -0.00 | -0.00 | -0.00 |
| G. Standard pipeline without the parallax correction | | | | | | | | | |
| $CC_{Friedel}$ | +0.00 | -0.00 | +0.00 | -0.00 | -0.00 | -0.00 | -0.00 | +0.00 | -0.00 |
| $CC_{Laue}$ | +0.00 | -0.00 | +0.00 | -0.00 | -0.00 | -0.00 | -0.00 | +0.00 | -0.00 |
| $CC_{1/2}$ | +0.00 | 0.00 | 0.00 | -0.00 | 0.00 | +0.00 | -0.00 | -0.00 | -0.00 |
| $CC_{Rep}$ | +0.00 | +0.00 | +0.00 | -0.00 | -0.00 | -0.00 | +0.00 | +0.00 | -0.00 |
| $CC_{LLM}$ | +0.00 | +0.00 | +0.00 | -0.00 | +0.00 | -0.00 | +0.00 | -0.00 | +0.00 |

**TABLE S7.** The CC statistics of each dataset processed with different image scale factors and with/without the radial profile variance removal method. The diffuse map generated by each processing method was evaluated with five CC metrics: $CC_{Friedel}$, $CC_{Laue}$, $CC_{1/2}$, $CC_{Rep}$, and $CC_{LLM}$ (anisotropic ADP model). Method A (standard processing pipeline) contains CC values of each dataset up to 1.4Å resolution, while other methods (B)-(H) show the relative CC changes compared to those in method A. (A)-(D) uses the radial profile variance removal method, while (E)-(H) turns it off. Each cell is colored in the same manner as Table S6.

| Sample | WT-1 | WT-2 | WT-3 | G150A-1 | G150A-2 | G150A-3 | G150T-1 | G150T-2 | G150T-3 |
|---|---|---|---|---|---|---|---|---|---|
| A. Radial profile scale factor, with the radial profile variance removal step (standard pipeline) | | | | | | | | | |
| $CC_{Friedel}$ | 0.93 | 0.91 | 0.91 | 0.93 | 0.91 | 0.91 | 0.91 | 0.92 | 0.94 |
| $CC_{Laue}$ | 0.90 | 0.87 | 0.87 | 0.88 | 0.86 | 0.85 | 0.86 | 0.88 | 0.91 |
| $CC_{1/2}$ | 0.85 | 0.78 | 0.81 | 0.81 | 0.76 | 0.77 | 0.82 | 0.84 | 0.89 |
| $CC_{Rep}$ | 0.86 | 0.84 | 0.85 | 0.82 | 0.81 | 0.82 | 0.88 | 0.87 | 0.89 |
| $CC_{LLM}$ | 0.70 | 0.71 | 0.67 | 0.70 | 0.68 | 0.73 | 0.76 | 0.75 | 0.80 |
| B. Water ring scale factor, with the radial profile variance removal step | | | | | | | | | |
| $CC_{Friedel}$ | +0.00 | +0.00 | -0.00 | -0.00 | -0.00 | -0.00 | -0.00 | -0.00 | -0.00 |
| $CC_{Laue}$ | +0.00 | +0.00 | -0.00 | -0.00 | +0.00 | -0.00 | -0.00 | -0.00 | +0.00 |
| $CC_{1/2}$ | 0.00 | +0.00 | -0.00 | 0.00 | 0.00 | -0.00 | -0.00 | -0.00 | 0.00 |
| $CC_{Rep}$ | -0.00 | -0.00 | -0.00 | -0.00 | -0.00 | -0.00 | -0.00 | +0.00 | -0.00 |
| $CC_{LLM}$ | -0.05 | -0.00 | -0.00 | +0.00 | +0.00 | +0.00 | +0.00 | +0.00 | +0.00 |
| C. Overall scale factor, with the radial profile variance removal step | | | | | | | | | |
| $CC_{Friedel}$ | +0.00 | +0.00 | -0.00 | +0.00 | +0.00 | -0.00 | +0.00 | -0.00 | +0.00 |
| $CC_{Laue}$ | +0.00 | +0.00 | -0.00 | +0.00 | +0.00 | -0.00 | +0.00 | -0.00 | +0.00 |
| $CC_{1/2}$ | 0.00 | +0.00 | -0.00 | +0.00 | +0.00 | -0.00 | +0.00 | 0.00 | +0.00 |
| $CC_{Rep}$ | -0.00 | -0.00 | -0.00 | -0.00 | +0.00 | -0.00 | -0.00 | -0.00 | -0.00 |
| $CC_{LLM}$ | +0.00 | -0.00 | +0.00 | +0.00 | +0.00 | -0.00 | +0.00 | +0.00 | +0.00 |
| D. Bragg intensity scale factor, with the radial profile variance removal step | | | | | | | | | |
| $CC_{Friedel}$ | -0.00 | -0.01 | -0.01 | -0.00 | -0.01 | -0.02 | -0.00 | -0.01 | -0.00 |
| $CC_{Laue}$ | -0.00 | -0.01 | -0.01 | -0.01 | -0.01 | -0.03 | -0.00 | -0.02 | -0.00 |
| $CC_{1/2}$ | -0.00 | -0.02 | -0.02 | -0.02 | -0.04 | -0.10 | -0.00 | -0.03 | -0.01 |
| $CC_{Rep}$ | -0.01 | -0.01 | -0.01 | -0.02 | -0.03 | -0.04 | -0.01 | -0.01 | -0.01 |
| $CC_{LLM}$ | -0.00 | -0.01 | -0.00 | -0.01 | -0.00 | -0.03 | +0.00 | +0.01 | +0.00 |
| E. Radial profile scale factor, without the radial profile variance removal step | | | | | | | | | |
| $CC_{Friedel}$ | -0.01 | -0.01 | -0.02 | -0.02 | -0.01 | -0.03 | -0.02 | -0.03 | -0.00 |
| $CC_{Laue}$ | -0.02 | -0.06 | -0.01 | -0.05 | -0.10 | -0.07 | -0.03 | -0.04 | -0.03 |
| $CC_{1/2}$ | -0.04 | -0.11 | -0.03 | -0.09 | -0.18 | -0.15 | -0.05 | -0.07 | -0.04 |
| $CC_{Rep}$ | -0.05 | -0.08 | -0.11 | -0.04 | -0.02 | -0.04 | -0.01 | -0.02 | -0.01 |
| $CC_{LLM}$ | +0.01 | -0.01 | +0.00 | -0.01 | -0.01 | -0.06 | -0.01 | -0.04 | -0.01 |
| F. Water ring scale factor, without the radial profile variance removal step | | | | | | | | | |
| $CC_{Friedel}$ | -0.00 | -0.00 | +0.01 | -0.01 | +0.00 | -0.01 | -0.01 | -0.02 | +0.00 |
| $CC_{Laue}$ | -0.01 | -0.04 | +0.01 | -0.03 | -0.07 | -0.04 | -0.02 | -0.02 | -0.02 |
| $CC_{1/2}$ | -0.02 | -0.07 | +0.01 | -0.05 | -0.13 | -0.08 | -0.03 | -0.04 | -0.02 |
| $CC_{Rep}$ | -0.02 | -0.01 | -0.04 | -0.01 | +0.00 | -0.00 | -0.00 | -0.01 | -0.00 |
| $CC_{LLM}$ | +0.00 | -0.00 | -0.01 | -0.01 | -0.01 | -0.03 | -0.01 | -0.03 | -0.01 |
| G. Overall scale factor, without the radial profile variance removal step | | | | | | | | | |
| $CC_{Friedel}$ | +0.00 | -0.00 | +0.01 | -0.01 | +0.00 | -0.00 | -0.00 | -0.01 | +0.00 |
| $CC_{Laue}$ | -0.01 | -0.04 | +0.01 | -0.02 | -0.07 | -0.03 | -0.02 | -0.01 | -0.01 |
| $CC_{1/2}$ | -0.02 | -0.07 | +0.01 | -0.04 | -0.12 | -0.06 | -0.02 | -0.03 | -0.02 |
| $CC_{Rep}$ | -0.01 | -0.01 | -0.03 | -0.01 | -0.01 | -0.01 | -0.00 | -0.01 | -0.00 |
| $CC_{LLM}$ | +0.00 | -0.00 | -0.02 | -0.01 | -0.01 | -0.03 | -0.01 | -0.02 | -0.01 |
| H. Bragg intensity scale factor, without the radial profile variance removal step | | | | | | | | | |
| $CC_{Friedel}$ | -0.04 | -0.08 | -0.09 | -0.15 | -0.03 | -0.20 | -0.10 | -0.17 | -0.08 |

| | | | | | | | | | |
|---|---|---|---|---|---|---|---|---|---|
| $CC_{Laue}$ | -0.40 | -0.46 | -0.10 | -0.36 | -0.47 | -0.33 | -0.36 | -0.30 | -0.38 |
| $CC_{1/2}$ | -0.79 | -1.00 | -0.27 | -0.85 | -0.98 | -0.83 | -0.73 | -0.64 | -0.77 |
| $CC_{Rep}$ | -0.52 | -0.46 | -0.64 | -0.74 | -0.86 | -0.67 | -0.38 | -0.63 | -0.39 |
| $CC_{LLM}$ | -0.13 | -0.27 | -0.28 | -0.26 | -0.35 | -0.48 | -0.09 | -0.20 | -0.12 |

**TABLE S8.** Statistics of the LLM model (up to 1.4Å resolution) for each dataset using the Refmac5-refined protein structures.[S2]

| Sample | WT-1 | WT-2 | WT-3 | G150A-1 | G150A-2 | G150A-3 | G150T-1 | G150T-2 | G150T-3 |
|---|---|---|---|---|---|---|---|---|---|
| Zero ADP | | | | | | | | | |
| $CC_{LLM}$ | 0.67 | 0.68 | 0.67 | 0.69 | 0.66 | 0.69 | 0.70 | 0.70 | 0.72 |
| $\gamma$ (Å) | 6.7 | 6.4 | 7.0 | 6.9 | 6.9 | 7.1 | 7.4 | 7.4 | 7.4 |
| $\sigma$ (Å) | 0.40 | 0.40 | 0.41 | 0.42 | 0.41 | 0.44 | 0.43 | 0.43 | 0.44 |
| Isotropic ADP | | | | | | | | | |
| $CC_{LLM}$ | 0.71 | 0.71 | 0.72 | 0.71 | 0.70 | 0.74 | 0.78 | 0.77 | 0.80 |
| $\gamma$ (Å) | 7.9 | 7.5 | 8.1 | 8.2 | 7.9 | 8.0 | 8.8 | 9.2 | 8.5 |
| $\sigma$ (Å) | <0.01 | <0.01 | <0.01 | <0.01 | <0.01 | <0.1 | <0.01 | <0.01 | <0.01 |
| Anisotropic ADP | | | | | | | | | |
| $CC_{LLM}$ | 0.70 | 0.71 | 0.67 | 0.70 | 0.68 | 0.73 | 0.76 | 0.75 | 0.80 |
| $\gamma$ (Å) | 7.9 | 7.2 | 8.8 | 8.4 | 8.3 | 7.6 | 11 | 11 | 9.1 |
| $\sigma$ (Å) | <0.01 | <0.1 | 0.15 | <0.01 | <0.1 | <0.1 | <0.01 | <0.01 | <0.1 |

**TABLE S9.** Statistics of the LLM model (up to 1.4Å resolution) for each dataset using the PHENIX-refined protein structures.[S1]

| Sample | WT-1 | WT-2 | WT-3 | G150A-1 | G150A-2 | G150A-3 | G150T-1 | G150T-2 | G150T-3 |
|---|---|---|---|---|---|---|---|---|---|
| Zero ADP | | | | | | | | | |
| $CC_{LLM}$ | 0.67 | 0.68 | 0.68 | 0.69 | 0.66 | 0.69 | 0.71 | 0.71 | 0.73 |
| $\gamma$ (Å) | 6.9 | 6.7 | 7.2 | 7.2 | 7.1 | 7.3 | 7.6 | 7.8 | 7.7 |
| $\sigma$ (Å) | 0.39 | 0.40 | 0.40 | 0.41 | 0.40 | 0.43 | 0.42 | 0.43 | 0.44 |
| Isotropic ADP | | | | | | | | | |
| $CC_{LLM}$ | 0.71 | 0.71 | 0.72 | 0.71 | 0.70 | 0.73 | 0.78 | 0.77 | 0.80 |
| $\gamma$ (Å) | 8.3 | 8.0 | 8.5 | 8.6 | 8.3 | 8.5 | 9.3 | 9.9 | 8.9 |
| $\sigma$ (Å) | <0.01 | <0.01 | <0.01 | <0.01 | <0.01 | <0.1 | <0.01 | <0.01 | <0.01 |
| Anisotropic ADP | | | | | | | | | |
| $CC_{LLM}$ | 0.61 | 0.62 | 0.61 | 0.70 | 0.62 | 0.69 | 0.74 | 0.75 | 0.77 |
| $\gamma$ (Å) | 9.7 | 9.3 | 10 | 8.6 | 9.4 | 8.7 | 14 | 13 | 12 |
| $\sigma$ (Å) | 0.18 | 0.23 | 0.23 | <0.01 | 0.15 | 0.12 | <0.1 | <0.01 | 0.11 |

**Table S10.** The R-factors calculated in CCTBX[S9] without bulk solvent or anisotropic scaling for Refmac5 and PHENIX models against the WT-1 dataset.

| | $R_{13}$ | $R_{23}$ | $R_{12}$ |
|---|---|---|---|
| (1) Refmac5; (2) PHENIX; (3) Experimental data | | | |
| Zero ADP | 0.499 | 0.500 | 0.107 |
| Iso ADP | 0.235 | 0.242 | 0.056 |
| Aniso ADP | 0.202 | 0.219 | 0.108 |
| (1) Refmac5; (2) PHENIX + $U_{ztr}$[1]; (3) Experimental data | | | |
| Zero ADP | 0.499 | 0.500 | 0.107 |
| Iso ADP | 0.235 | 0.242 | 0.056 |
| Aniso ADP | 0.202 | 0.207 | 0.061 |
| (1) Refmac5 - $U_{ztr}$; (2) PHENIX; (3) Experimental data | | | |
| Zero ADP | 0.499 | 0.500 | 0.107 |
| Iso ADP | 0.235 | 0.242 | 0.056 |
| Aniso ADP | 0.216 | 0.219 | 0.061 |

[1]$U_{ztr}$ is the zero-trace U matrix that relates Refmac5- and PHENIX-refined PDBs.

**TABLE S11.** $CC_{Bragg}$ and $CC_{LLM}$ between experimental data and calculated Refmac5, PHENIX models of the WT-1 dataset. The $CC_{Bragg}$ is calculated between Refmac5 and PHENIX models from 10 to 1.4Å resolution. The $CC_{Bragg}$ and $CC_{LLM}$ values were computed in a similar way using only the anisotropic component of the data and are comparable indicators of the sensitivity to the rescaling of the model anisotropic ADPs using $U_{ztr}$. The changes in $CC_{LLM}$ upon rescaling are much larger than the changes in $CC_{Bragg}$. Like the $CC_{Bragg}$ values, the $CC_{Bragg,total}$ values, computed using the Bragg intensities prior to subtracting the isotropic component, are insensitive to changes in the model ADPs.

| | Refmac5 | PHENIX | Refmac5-$U_{ztr}$[1] | PHENIX+$U_{ztr}$ |
|---|---|---|---|---|
| $CC_{Bragg,total}$ | 0.931 | 0.930 | 0.929 | 0.932 |
| $CC_{Bragg}$ | 0.892 | 0.893 | 0.890 | 0.895 |
| $CC_{LLM}$ | 0.709 | 0.605 | 0.608 | 0.707 |
| Refmac5 | | 0.989 | 0.994 | 0.995 |
| PHENIX | | | 0.995 | 0.995 |
| Refmac5-$U_{ztr}$ | | | | 0.990 |

[1]$U_{ztr}$ is the zero-trace U matrix that relates Refmac5- and PHENIX-refined PDBs.

**TABLE S12.** Statistics of the rigid-body translational motions (RBT) model (up to 1.4Å resolution) for each dataset using the Refmac5-refined protein structures.[S2]

| Sample | WT-1 | WT-2 | WT-3 | G150A-1 | G150A-2 | G150A-3 | G150T-1 | G150T-2 | G150T-3 |
|---|---|---|---|---|---|---|---|---|---|
| Zero ADP | | | | | | | | | |
| $CC_{RBT}$ | 0.53 | 0.52 | 0.55 | 0.54 | 0.54 | 0.55 | 0.49 | 0.49 | 0.50 |
| $\sigma$ (Å) | 0.37 | 0.40 | 0.38 | 0.39 | 0.38 | 0.40 | 0.35 | 0.37 | 0.37 |
| Isotropic ADP | | | | | | | | | |
| $CC_{RBT}$ | 0.59 | 0.58 | 0.61 | 0.60 | 0.59 | 0.60 | 0.55 | 0.56 | 0.57 |
| $\sigma$ (Å) | <0.1 | <0.1 | <0.1 | <0.01 | <0.1 | <0.1 | <0.01 | <0.01 | <0.01 |
| Anisotropic ADP | | | | | | | | | |
| $CC_{RBT}$ | 0.59 | 0.58 | 0.60 | 0.60 | 0.59 | 0.60 | 0.54 | 0.54 | 0.57 |
| $\sigma$ (Å) | <0.01 | 0.10 | <0.1 | <0.01 | <0.01 | <0.1 | <0.01 | <0.01 | <0.01 |

**TABLE S13.** Statistics of the RBT model (up to 1.4Å) for each dataset using the PHENIX-refined protein structures.[S1]

| Sample | WT-1 | WT-2 | WT-3 | G150A-1 | G150A-2 | G150A-3 | G150T-1 | G150T-2 | G150T-3 |
|---|---|---|---|---|---|---|---|---|---|
| Zero ADP | | | | | | | | | |
| $CC_{RBT}$ | 0.54 | 0.53 | 0.56 | 0.55 | 0.54 | 0.56 | 0.50 | 0.50 | 0.51 |
| $\sigma$ (Å) | 0.37 | 0.39 | 0.38 | 0.39 | 0.38 | 0.40 | 0.35 | 0.36 | 0.36 |
| Isotropic ADP | | | | | | | | | |
| $CC_{RBT}$ | 0.60 | 0.59 | 0.62 | 0.61 | 0.60 | 0.61 | 0.55 | 0.56 | 0.54 |
| $\sigma$ (Å) | <0.01 | <0.01 | <0.01 | <0.01 | <0.01 | <0.01 | <0.01 | <0.01 | <0.01 |
| Anisotropic ADP | | | | | | | | | |
| $CC_{RBT}$ | 0.55 | 0.55 | 0.57 | 0.60 | 0.56 | 0.59 | 0.52 | 0.54 | 0.55 |
| $\sigma$ (Å) | <0.1 | 0.15 | 0.15 | <0.1 | <0.1 | <0.1 | <0.01 | <0.01 | <0.01 |

### III. Conversion between Refmac5- and PHENIX-refined PDBs

Refmac5 and PHENIX programs choose different overall anisotropic scaling parameters, and these numbers are saved as #REMARK information in the headers of Refmac5-refined PDBs ($B^R_{HKL}$), and in PHENIX-refined PDBs ($B^P_{HKL}$) output by PDB-REDO after zero cycles of refinement[S8].

The overall anisotropic scale matrices ($B^R_{HKL}$ and $B^P_{HKL}$) in #REMARK are in the unit cell basis, and can be converted into the PDB Cartesian basis ($B^R_{XYZ}$ and $B^P_{XYZ}$) using the equations:

$$B^R_{XYZ} = A^{-T}S^{-T}B^R_{HKL}S^{-1}A^{-1}$$
$$B^P_{XYZ} = A^{-T}S^{-T}B^P_{HKL}S^{-1}A^{-1},$$

where the $A$ and $S$ matrices are defined as follows,

$$A = \begin{pmatrix} a^*_x & b^*_x & c^*_x \\ a^*_y & b^*_y & c^*_y \\ a^*_z & b^*_z & c^*_z \end{pmatrix}; \quad S = \begin{pmatrix} 1/|a^*| & 0 & 0 \\ 0 & 1/|b^*| & 0 \\ 0 & 0 & 1/|c^*| \end{pmatrix}.$$

We can derive the U matrix as the difference between two anisotropic scaling matrices in Cartesian coordinates, scaled by $2\pi^2$, using the equation,

$$\mathrm{U} = \frac{1}{2\pi^2}(B^P_{XYZ} - B^R_{XYZ}),$$

Three examples of this conversion for WT-1, G150A-1, and G150T-1 are shown in Table S4.

In addition, we can interconvert anisotropic ADPs in the Refmac5- and PHENIX-refined models by adding or subtracting the traceless U matrix ($U_{ztr}$) to the ANISOU arrays in the PDB file. We use a zero-trace $U_{ztr}$ matrix, calculated by removing trace/3 from the diagonal of the U matrix, to avoid changing $B_{eq}$ values. Addition of the $U_{ztr}$ matrix to the PHENIX anisotropic ADP transforms them into ADPs that closely resemble the Refmac5 values for most atoms (i.e., PHENIX+$U_{ztr}$ ⇔ Refmac5; Fig. S4). Conversely, subtraction of the $U_{ztr}$ matrix from Refmac5 anisotropic ADPs converts them to PHENIX-like values (i.e., Refmac5-$U_{ztr}$ ⇔ PHENIX; Fig. S5). Although this transformation does not make two PDB files identical in all aspects, it does make the anisotropy distributions of both models more similar to each other, especially for protein atoms, as shown in Fig. S1, S4, and S5. Importantly, it also fixes the discrepancy in $CC_{LLM}$ values while keeping the $CC_{Bragg}$ values largely unchanged for both models, as shown in Table S11 and Fig. 9. Therefore, the major differences in the anisotropic ADP models produced by the two programs arise from different initial choices of anisotropic scale matrices that are applied before restrained coordinate and ADP refinement. While the differing anisotropic scale matrices produced by Refmac5 and PHENIX have little effect on the model statistics against the Bragg data (similar $R_{free}/R_{work}$ and nearly identical $CC_{Bragg}$ values), they do affect the agreement of the LLM models with diffuse scattering data. We therefore conclude that diffuse scattering data appear to be more sensitive than Bragg data to the details of the anisotropic ADP model, at least for these ICH datasets.

### IV. Liquid-like motions model of diffuse scattering with individual atomic B factors

Here we copy the derivation from Ref. S10 showing that individual atomic B factors can be included in the LLM model of diffuse scattering by eliminating the overall Debye-Waller factor $\left(e^{-|q|^2\sigma^2}\right)$, and replacing $I_0(q)$, the squared structure factor of the unperturbed crystal, with $I_B(q)$, the Bragg intensity computed from the participating atoms.

The total squared structure factor of a crystal can be computed using the following equation:

$$I = \sum_{jk}\sum_{nm} f_m f_n^* e^{iq\cdot(R_j - R_k + x_m - x_n)} e^{iq\cdot(d_{jm} - d_{kn})}, \qquad (1)$$

where $R_j$ is the position of the origin of unit cell $j$, $x_m$ is the reference position of atom $m$ within each unit cell, and $d_{jm}$ is the deviation of the position of atom $m$ from the reference in unit cell $j$. Assuming atom displacements are statistically homogeneous, we replace the phase factor of the deviations with an average over the unit cell, then sum over $k$, noting that the difference between two lattice vectors is also a lattice vector:

$$I = N\sum_{j}\sum_{nm} f_m f_n^* e^{iq\cdot(r_{jmn})} \langle e^{iq\cdot(d_{jm} - d_{kn})}\rangle_k, \qquad (2)$$

where

$$r_{jmn} = R_j + x_m - x_n \qquad (3)$$

Now, as is standard, e.g., in calculation of the Debye-Waller factors, use the harmonic approximation to evaluate the average over the phase factor due to deviations:

$$I = N\sum_{j}\sum_{nm} f_m f_n^* e^{iq\cdot(r_{jmn})} e^{-\frac{1}{2}\langle[q\cdot(d_{jm} - d_{kn})]^2\rangle_k}. \qquad (4)$$

Further evaluate the average to obtain

$$I = N \sum_j \sum_{nm} f_m f_n^* e^{iq\cdot(r_{jmn})} e^{-\frac{1}{2}q\cdot U_m \cdot q} e^{-\frac{1}{2}q\cdot U_n \cdot q} e^{q\cdot U_{mn}(r_{jmn})\cdot q} \tag{5}$$

The exponential factors correspond to Debye-Waller factors for individual atoms (involving $U_m$ and $U_n$) and a factor for the correlated displacements of two atoms (involving $U_{mn}$). We will make some assumptions about the $U_{mn}$ factor to derive the modified LLM model, but first, make sure the incoherent scattering ($j = 0$ and $m = n$) is properly accounted for when choosing an arbitrary $U_{mn}$ that might not precisely match the individual atomic displacements:

$$I = N \sum_j \sum_{nm} f_m f_n^* e^{iq\cdot(r_{jmn})} e^{-\frac{1}{2}q\cdot U_m \cdot q} e^{-\frac{1}{2}q\cdot U_n \cdot q} e^{q\cdot U_{mn}(r_{jmn})\cdot q} + N \sum_m |f_m|^2 \left(1 - e^{-q\cdot U_m \cdot q} e^{q\cdot U_{mm}(0)\cdot q}\right) \tag{6}$$

Now separate the Bragg term and isolate the diffuse intensity from both the incoherent correction and Bragg:

$$I = N \sum_j \sum_{nm} f_m f_n^* e^{iq\cdot(r_{jmn})} e^{-\frac{1}{2}q\cdot U_m \cdot q} e^{-\frac{1}{2}q\cdot U_n \cdot q} + N \sum_j \sum_{nm} f_m f_n^* e^{iq\cdot(r_{jmn})} e^{-\frac{1}{2}q\cdot U_m \cdot q} e^{-\frac{1}{2}q\cdot U_n \cdot q} \left(e^{q\cdot U_{mn}(r_{jmn})\cdot q} - 1\right) + N \sum_m |f_m|^2 \left(1 - e^{-q\cdot U_m \cdot q} e^{q\cdot U_{mm}(0)\cdot q}\right) \tag{7}$$

From which we identify the first term as corresponding to the Bragg:

$$I_B = N \sum_j \sum_{nm} f_m f_n^* e^{iq\cdot(r_{jmn})} e^{-\frac{1}{2}q\cdot U_m \cdot q} e^{-\frac{1}{2}q\cdot U_n \cdot q}, \tag{8}$$

the middle term corresponding to the diffuse:

$$I_D = N \sum_j \sum_{nm} f_m f_n^* e^{iq\cdot(r_{jmn})} e^{-\frac{1}{2}q\cdot U_m \cdot q} e^{-\frac{1}{2}q\cdot U_n \cdot q} \left(e^{q\cdot U_{mn}(r_{jmn})\cdot q} - 1\right), \tag{9}$$

and the last term corresponding to the incoherent correction:

$$I_{inc} = N \sum_m |f_m|^2 \left(1 - e^{-q\cdot U_m \cdot q} e^{q\cdot U_{mm}(0)\cdot q}\right). \tag{10}$$

Now, the key assumption of the LLM is that the $U_{mn}(r_{jmn})$ has the same form for all pairs of atoms, and only depends on the separation vector $r_{jmn}$. In this case $U_{mn}(r_{jmn})$ can be written as

$$U_{mn}(r_{jmn}) = Vc(r_{jmn}), \tag{11}$$

where $V$ is a (potentially anisotropic) matrix of variations and $c(r_{jmn})$ is a scalar function with values in the range [-1,1] describing the correlation vs separation. Making this substitution and using a Taylor expansion for the exponential yields

$$N\sum_{l=1}^{\infty}\frac{(q\cdot V\cdot q)^l}{l!}\sum_j\sum_{nm} e^{iq\cdot(r_{jmn})}f_m f_n^* e^{-\frac{1}{2}q\cdot U_m\cdot q}e^{-\frac{1}{2}q\cdot U_n\cdot q}c^l(r_{jmn}), \tag{12}$$

which, using the Dirac delta function, is equivalent to

$$\sum_{l=1}^{\infty}\frac{(q\cdot V\cdot q)^l}{l!}\int d^3r\, e^{iq\cdot r}c^l(r)N\sum_j\sum_{nm}\delta(r-r_{jmn})f_m f_n^* e^{-\frac{1}{2}q\cdot U_m\cdot q}e^{-\frac{1}{2}q\cdot U_n\cdot q}. \tag{13}$$

Note that

$$\int d^3r\, e^{iq\cdot r}N\sum_j\sum_{nm}\delta(r-r_{jmn})f_m f_n^* e^{-\frac{1}{2}q\cdot U_m\cdot q}e^{-\frac{1}{2}q\cdot U_n\cdot q}$$
$$= N\sum_j\sum_{nm} e^{iq\cdot(r_{jmn})}f_m f_n^* e^{-\frac{1}{2}q\cdot U_m\cdot q}e^{-\frac{1}{2}q\cdot U_n\cdot q} = I_B, \tag{14}$$

which identifies the double sum in Eq. (12) as the Patterson of the mean electron density of the crystal, $P(r)$. We therefore have

$$I_D = \sum_{l=1}^{\infty}\frac{(q\cdot V\cdot q)^l}{l!}\int d^3r\, e^{iq\cdot r}c^l(r)P(r), \tag{15}$$

or, by the convolution theorem,

$$I_D = \sum_{l=1}^{\infty}\frac{(q\cdot V\cdot q)^l}{l!}I_B * \int d^3r\, e^{iq\cdot r}c^l(r), \tag{16}$$

which, to first order, is

$$I_D = q\cdot V\cdot qI_B * FT[c(r)], \tag{17}$$

where FT represents the Fourier Transformation.

Compared to Eq. (2), this equation is missing the leading factor $e^{-q\cdot V\cdot q}$. Eq. (2) otherwise has the same form as the first order term in this expansion, but with the unperturbed squared structure factor $I_0$ (computed without individual atom B factors) replaced by the Bragg intensity $I_B$ (computed using the individual atom B factors).

As an example, the following is the equation for the isotropic LLM using individual B factors, in which the correlation decreases exponentially with distance:

$$I_D = |q|^2\sigma^2 I_B * FTe^{-\frac{|r|}{\gamma}}, \tag{18}$$

yielding

$$I_D(q) = |q|^2\sigma^2 I_B(q) * \frac{8\pi\gamma^3}{[1+(\gamma|q|)^2]^2}. \tag{19}$$
